# Fishing for new immunotherapy compounds to boost innate tumor rejection

**DOI:** 10.64898/2026.11.15.766748

**Authors:** A.B. Machado, C. Labão-Almeida, I. Oliveira, L. Leite, C. Rebelo de Almeida, B. Costa, J. Xavier de Carvalho, B. Sebo, D. Calafato, V. Póvoa, R. Couto, C. Tange, T. Rodrigues, G.J.L. Bernardes, R. Fior

## Abstract

Despite significant advances in cancer immunotherapy, tackling immunosuppression remains critical to achieve more effective responses. While developing the zebrafish xenograft model, we found that some human tumors engraft very efficiently (“progressors”), while others are cleared by the host (“regressors”). We showed that this was dependent on innate immunity, which recognizes and eliminates some cancer cells, while others are able to evade and suppress it. This opened an opportunity to perform an *in vivo* phenotypic drug screen to find compounds that induce clearance by boosting innate-tumor rejection.

Zebrafish xenografts of human colorectal (CRC) and breast cancer (BC) that show a high engraftment/low clearance phenotype were used to screen over 500 FDA-approved small-molecule compounds, with tumor clearance serving as the phenotypic readout. The screen identified 23 hit compounds, of which six were effective across both cancer models. Mechanism-of-action enrichment analysis revealed a significant overrepresentation of autonomic nervous system-targeting drugs among the hits, particularly cholinergic ones. Immune profiling in myeloid reporter lines showed that the lead hit, Tiotropium bromide, a muscarinic receptor antagonist, reprograms tumor-associated macrophages toward a pro-inflammatory, anti-tumoral state. Tiotropium significantly increased macrophage phagocytosis, and macrophage depletion completely abolished its pro-clearance activity, establishing these myeloid cells as essential effectors. In a syngeneic CT26 colorectal mouse model, Tiotropium synergized with anti-PD-1 to control tumor growth and improve survival, outperforming all monotherapy arms. This effect was immune-dependent, as it was lost in immunodeficient NSG mice.

Together, these findings demonstrate that *in vivo* phenotypic screening of FDA-approved compounds in zebrafish xenografts is a productive strategy for innate immune drug discovery in cancer. The identification of Tiotropium as a myeloid-reprogramming innate immune primer, together with the unbiased recovery of the neuro-immune axis as a druggable target, opens new avenues for overcoming immunotherapy resistance.

## Introduction

Immunotherapy, particularly immune checkpoint blockade (ICB), has transformed cancer treatment, establishing that the immune system can be durably engaged to eliminate tumors^1^. Yet, many patients fail to benefit from these therapies, as resistance arises due to tumor-intrinsic factors such as low immunogenicity and antigen loss, as well as insufficient T cell infiltration and engagement of alternative immune suppression pathways^2^. Critically, the tumor microenvironment (TME) of most solid cancers is dominated not by T cells, but by innate myeloid populations, particularly macrophages and neutrophils^3^. Innate cells constitute the earliest immune barrier encountered by emerging tumors and critically shape subsequent adaptive responses, but despite their central role in tumor evolution and surveillance, they remain underexplored as therapeutic targets^4^. While some innate–targeted strategies, such as CD47/SIRPα blockade^5^ and CSF1R inhibitors^6^, have shown compelling activity in preclinical models, clinical translation has been limited by toxicity, insufficient efficacy, and difficulty in engaging the complex cellular crosstalk of the human TME^7,8^. These observations underscore the need for therapeutic strategies that effectively harness innate immunity in cancer, but have controlled toxicity, improving the depth and durability of clinical responses.

A fundamental challenge in drug discovery for oncology is the high attrition rate, as more than 90% of cancer drugs that enter clinical trials fail to reach approval, and even those that succeed often provide only modest survival benefits^9–11^. This persistent gap between preclinical promise and clinical success has led to renewed interest in phenotypic drug discovery (PDD), an approach in which compounds are screened based on their effects in disease–relevant biological systems, as an alternative to target-based drug discovery (TDD), which focuses on modulating specific molecular targets^12^. Notably, analysis of U.S. Food and Drug Administration (FDA) approvals between 1999 and 2008 revealed that PDD contributed more first–in–class small–molecule drugs than the then-dominant TDD approaches^13^. One particularly efficient application of PDD is drug repurposing, in which libraries of FDA–approved, clinically used compounds are screened for new therapeutic applications. Because these compounds already have established safety profiles, pharmacokinetic properties, and established mechanisms of action, repurposing substantially accelerates clinical translation. In cancer immunotherapy, this approach offers an attractive opportunity to identify immune modulators with immediate translational potential. Yet, few phenotypic screens have directly explored immune–tumor interactions^14–16^ and, to the best of our knowledge, none have done it *in vivo*.

Zebrafish phenotypic screens have shown strong translational success across therapeutic areas, identifying several compounds now in advanced clinical testing^17–19^, such as prostaglandin E2 derivatives (umbilical cord blood transplantation and graft versus-host disease)^20–22^, bone morphogenetic protein receptor inhibitors (fibrodysplasia ossificans progressiva)^23–25^, and benzothiophene carboxamides (antibiotic-induced hearing loss prevention)^26,27^ for example. With ∼82% genetic conservation with human disease genes, plus high fecundity, rapid development, and low cost compared to mammalian models, zebrafish excel for large-scale screening^17,28,29^. They also absorb small molecules through their skin, simplifying compound administration. Zebrafish xenografts, in which tumor cells are transplanted into embryos a few days post-fertilization, provide a uniquely suited system for innate immune-targeted PDD, as larvae lack fully functional adaptive immunity until 3–6 weeks post-fertilization^30–32^, meaning all immune-tumor responses are exclusively innate–mediated. Larval optical transparency and transgenic reporter lines enable real-time, single-cell-resolution visualization of innate cell behavior^33^.

We previously established that human cancer cell lines exhibit contrasting engraftment phenotypes in zebrafish xenografts: some are recognized and cleared (“regressors”), whereas others persist by evading or suppressing host innate immunity (“progressors”)^34,35^. Thus, progressor xenografts provide the ideal platform for an *in vivo* phenotypic drug screen for innate immune modulators: a robust, reproducible baseline of tumor engraftment against which compounds that induce innate tumor-clearance can be identified.

Here, we report an *in vivo* phenotypic drug screen using zebrafish xenografts of two human cancer progressor cell lines, colorectal HCT116 and breast cancer Hs578T, against an FDA–approved compound library of over 500 drugs. Two rounds of screening identified 23 confirmed hits, six effective across both cancer models. Mechanism–of–action enrichment analysis revealed a significant representation of autonomic nervous system (ANS)–modulating drugs among the hits, with muscarinic receptor (MR) antagonists, dopaminergic agents, and phosphodiesterase (PDE) inhibitors constituting the most enriched pharmacological classes. One of the top hits, Tiotropium, a MR antagonist approved for chronic obstructive pulmonary disease, induced macrophage–dependent tumor clearance across multiple progressor xenograft models, increased pro–inflammatory macrophage activation at the tumor site, and synergized with anti–PD–1 immune checkpoint blockade to reduce tumor volume and improve survival in immune–competent mice. Together, these results establish *in vivo* zebrafish xenograft screening as an efficient platform for innate immune drug discovery, and identify repurposed autonomic drugs, Tiotropium in particular, as a new class of innate immune modulators with potential to sensitize cold tumors to checkpoint therapy.

## Results

### A zebrafish xenograft phenotypic drug screen identifies compounds that boost innate tumor clearance

To identify drugs that enhance innate anti-tumor immunity, we performed a phenotypic drug screen of an FDA-approved small compound library using zebrafish xenografts of two human progressor cell lines, CRC HCT116 and BC Hs578T, i.e., cancer cells that have low clearance rates (

### Main

Figure 1). Briefly, fluorescently labeled cancer cells were injected into the perivitelline space (PVS) of 2 days post-fertilization (dpf) zebrafish embryos. Compounds from the drug library were tested at 10 μM, a standard concentration for library screens^36^, with daily renewal from 1 to 4/6 days post-injection (dpi). Tumor clearance, defined as the absence of a detectable tumor, was quantified relative to vehicle-treated controls at the end of the assay and served as our phenotypic readout for immune-mediated rejection.

**Figure 1:**
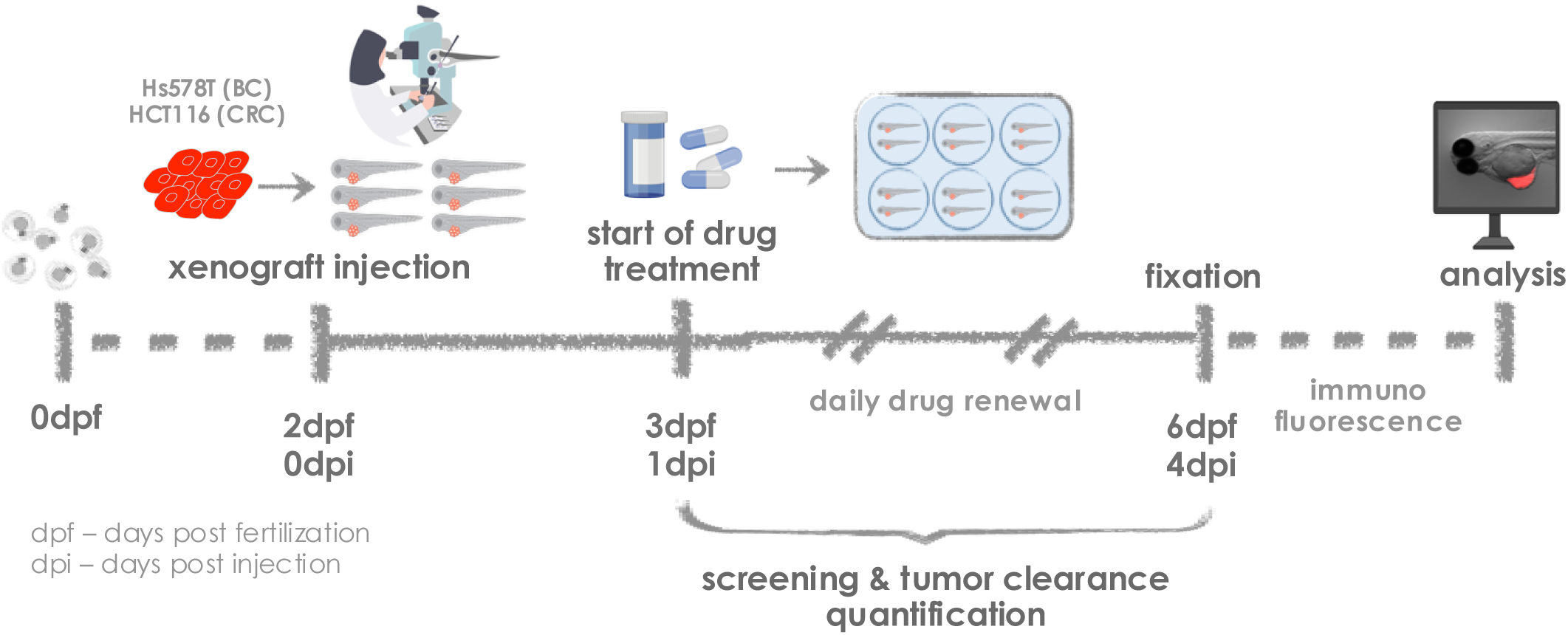
A phenotypic drug screen using zebrafish xenografts discovers new compounds that boost innate tumor clearance. Schematic representation of the phenotypic drug screen using zebrafish xenografts. CRC HCT116 or BC Hs578T cells were injected into the perivitelline space (PVS) of 2 days post-fertilization (dpf) zebrafish embryos. Compounds from the drug library were diluted in E3 fish media at 10uM and renewed daily. Tumor clearance was screened and quantified at 4 or 6 days post-injection (dpi), when embryos were fixed for immunofluorescence and further analysis.

We have previously shown that these progressor models are characterized by consistently low baseline rates of immune clearance (∼10% at 4 dpi)^35^ and that most compounds with tumor-autonomous effects, such as chemotherapy, lead to a reduction in tumor size but are not capable of inducing complete clearance in the 4 to 6-day time window of this assay^34^. Therefore, compounds capable of increasing tumor rejection in at least two independent rounds were considered screen hits, as they are expected to act through the immune system rather than by direct cytotoxicity.

Of a total of 556 screened compounds (**Supplementary Data 1)**, 81 were excluded due to toxicity and 64 unique drugs moved on to second-round re-testing, totaling 14 confirmed hits for HCT116 (2.56% hit rate) and 15 for Hs578T (3.84%) xenografts (**Table 1**, **Supplementary Figure 1**). Importantly, six drugs yielded effects in both models, suggesting they may act through broad immune-mediated mechanisms rather than cancer-type-specific ones.

**Table 1:** Summary statistics of the zebrafish xenograft phenotypic drug screen. Compounds causing developmental toxicity or lethality were excluded from the screen. First round candidates were re-tested in a second independent round, and confirmed hits were defined as compounds that reproducibly increased tumor clearance relative to control in both rounds. Common hits indicate compounds confirmed in both CRC and BC xenograft models. Hit rate was calculated as the number of confirmed hits divided by the total number of screened compounds per model.

| **Xenograft model** | **Total screened compounds** | **Excluded due to toxicity** | **Re-tested in 2<sup>nd</sup> round** |  | **Confirmed hits** |  | **Hit rate** |
|---|---|---|---|---|---|---|---|
| HCT116 | 546 | 81 | 40 | 8 | 14 | 6 | 2.56% |
| Hs578T | 391 |  | 35 |  | 15 |  | 3.84% |

### Mechanism-of-action enrichment reveals over-representation of cholinergic drugs

Because the drug library we used consists exclusively of FDA-approved compounds with known clinical indications and pharmacological targets, we sought to determine whether particular mechanisms of action (MOA) were overrepresented beyond what chance would expect among the 23 unique hit compounds (**Table 2**). Therefore, we performed an enrichment analysis comparing the distribution of MOA classes among the hits against the full screened library, which showed a combined hit rate of 4.1% for both models (**Figure 2**, **Supplementary Data 2**).

**Table 2:** Hit compounds identified in the drug screen. For each compound, the primary mechanism of action and clinical indication are shown. Average Δ tumor clearance represents the mean difference in the percentage of fish with no tumor at the end of the assay between treated and control conditions, at 4 or 6 dpi, pooled across independent experiments. COMT - catechol-O-methyltransferase, COX - cyclooxygenase, DDC - dopa decarboxylase, DHFR - dihydropteroate synthetase, NRTI - nucleoside reverse transcriptase inhibitors, NSAID - non-steroidal anti-inflammatory drug, ROS - reactive oxygen species, PBP - penicillin-binding protein, PDE - phosphodiesterase.

| **Compound** | **Mechanism of action** | **Indication** | **Average Δ tumor clearance (%)** |  |
|---|---|---|---|---|
|  |  |  | **HCT116** | **Hs578T** |
| Amrinone | PDE3 inhibitor | Vasodilator | 19.2 | 18.3 |
| Dicyclomine·HCl | Muscarinic receptor antagonist | Antispasmodic | 21.5 | 28.0 |
| Ketoprofen | COX inhibitor | NSAID / Analgesic | 18.7 | 30.3 |
| Methyldopa Sesquihydrate | α-adrenoceptor agonist | Antihypertensive | 14.0 | 16.0 |
| Tiotropium Bromide | Muscarinic receptor antagonist | Bronchodilator | 30.0 | 24.0 |
| Trimethoprim | DHFR inhibitor | Antibiotic | 17.5 | 27.0 |
| Ceftizoxim·Na | β-lactam (PBP inhibitor) | Antibiotic | 19.5 | — |
| Fexofenadine·HCl | H1 receptor antagonist | Antihistamine | 16.0 | — |
| Scopolamine·HBr | Muscarinic receptor antagonist | Antiemetic / Anticholinergic | 20.5 | — |
| Sumatriptan Succinate | 5-HT1B/1D receptor agonist | Antimigraine | 14.0 | — |
| Pentoxifylline | PDE inhibitor | Hematologic agent | 23.0 | — |
| Piperacillin | β-lactam (PBP inhibitor) | Antibiotic | 23.0 | — |
| Primaquine diphosphate | ROS generator | Antimalarial | 20.5 | — |
| Zalcitabine | NRTI | Antiviral / Anti-HIV | 18.0 | — |
| Bendamustine·HCl | DNA alkylating agent | Antineoplastic | — | 24.0 |
| Disopyramide | Na<sup>+</sup> channel blocker | Antiarrhythmic | — | 21.0 |
| Ethambutol Dihydrochloride | Arabinosyl transferase inhibitor | Antitubercular | — | 29.0 |
| Etodolac | COX inhibitor | NSAID / Analgesic | — | 30.0 |
| Fluconazole | Ergosterol synthesis inhibitor | Antifungal | — | 27.7 |
| Glycopyrrolate Iodide | Muscarinic receptor antagonist | Anticholinergic / Adjuvant | — | 26.0 |
| Metoprolol Tartrate | β-adrenoceptor antagonist | Antihypertensive | — | 12.7 |
| S(-)-Carbidopa | DDC inhibitor | Antiparkinsonian | — | 22.5 |
| Tolcapone | COMT inhibitor | Antiparkinsonian | — | 16.5 |

**Figure 2:**
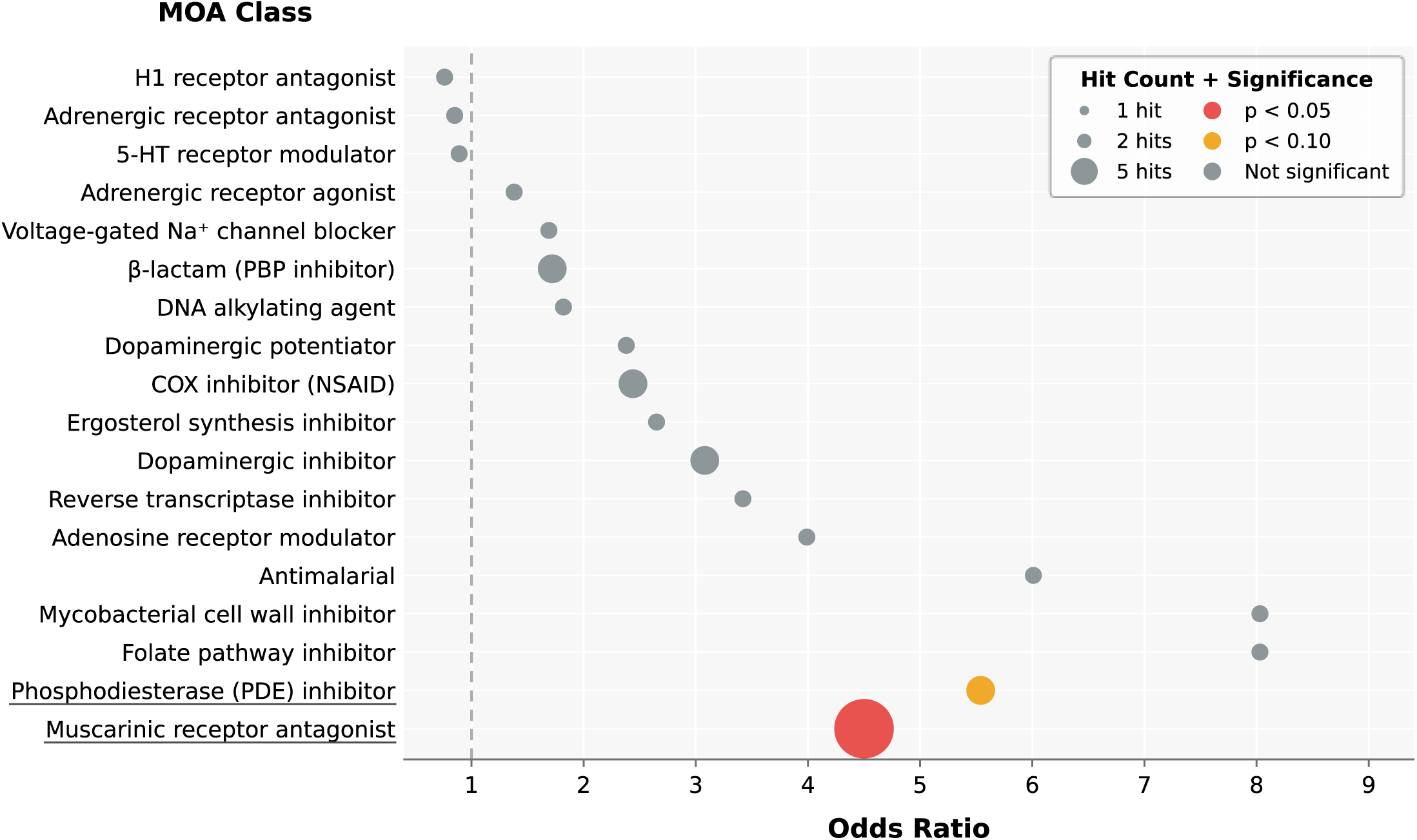
Hit compounds show enrichment of certain mechanisms of action. Dot plot showing the enrichment of mechanism of action (MOA) classes among hit compounds, quantified by odds ratio. Dot size is proportional to the number of hit compounds within that class (1-4 hits) and color indicates statistical significance, assessed by Fisher’s exact test: red (p < 0.05), orange (p < 0.10), gray (not significant).

Muscarinic Receptor (MR) antagonists were the only significantly enriched class, with 5 of 36 library compounds confirmed as hits (Dicyclomine, Disopyramide, Glycopyrrolate Iodide, Scopolamine·HBr, and Tiotropium Bromide; 13.9% hit rate, p=0.012), compared with the 1.5 expected by chance. PDE inhibitors showed a trend toward enrichment, with 2 hits among 11 compounds (Amrinone and Pentoxifylline; 18.2%, *p*=0.072).

Additionally, non–steroidal anti–inflammatory drugs (NSAIDs), Ketoprofen and Etodolac, were also represented in the enrichment analysis. Cyclooxygenase (COX) inhibition has already been described to counteract prostaglandin–driven immunosuppression of myeloid populations^37^, with several NSAIDs, such as aspirin, currently being explored in combination with ICB^38,39^. The recovery of this drug class serves as an internal positive control, validating our overall screen rationale.

Several of the represented classes among hit compounds act on neurotransmitter signaling (**Figure 2**), prompting us to ask whether this reflected a broader pattern. We therefore grouped all screened compounds by the neurotransmitter system their molecular target belongs to, irrespective of MOA class or clinical indication, dividing them into cholinergic, dopaminergic, adrenergic, purinergic, histaminergic, serotonergic, glutamatergic, GABAergic, or opioidergic (**Supplementary Data 3**).

Compounds acting on the autonomic nervous system (ANS), comprising the cholinergic, dopaminergic, and adrenergic classes, accounted for 9 of the 23 hits, representing 17.3% of the library (*p*=0.0093). Individually, cholinergic compounds were significantly enriched among the hits (5 of 40, 12.5%, *p*=0.019), dopaminergic compounds showed an enrichment trend (3 of 29, 10.3%, *p*=0.11), while adrenergic compounds, the largest system in the library, returned a hit rate at baseline (2 of 46, 4.3%). The remaining systems did not return hit rates significantly different from the library one, including the serotonergic (1 of 39, 2.6%) and the histaminergic (1 of 34, 2.9%), or were too sparsely represented for a meaningful test. These results highlight an autonomic signature in our screen, driven mainly by cholinergic compounds, pointing to parasympathetic signaling as a candidate regulator of innate anti-tumor immunity.

To assess how the composition of the library shapes the MOA enrichment analysis, we examined the distribution of the overall approved pharmacopeia. MR antagonists act on aminergic G protein-coupled receptors (GPCR), which are the target of 31-36% of approved drugs^40–42^. Our library mirrors this composition, with 149 of 556 compounds (26.8%) acting on GPCRs and 110 of those (74%) on aminergic receptors, raising the possibility that enrichment could simply reflect class abundance. Enrichment, however, did not reflect class size. Despite MR antagonists being the largest MOA class in the library (36 compounds, 13.9% hit rate), none of the seven classes of comparable size reached significance, including other aminergic classes such as H1 receptor antagonists (1 of 31, 3.2%) and adrenergic receptor antagonists (1 of 28, 3.6%), as well as non-aminergic glucocorticoid receptor agonists (0 of 24) and sex hormone receptor modulators (0 of 23) (**Supplementary Data 2**). The highest hit rate among these comparably sized classes belonged to the NSAIDs (2 of 22, 9.1%, p=0.229), consistent with their role as an internal positive control. The same applied to PDE inhibitors, whose 18.2% hit rate (2 of 11) was the highest of any class of comparable size (8 to 15 compounds).

Sampling depth nonetheless limits what this analysis can detect. With 23 hits among 556 compounds, a class must contain at least 24 compounds before even one hit would be expected by chance, and only 5 of the 115 MOA classes are that large. For all the others, an absence of hits is uninformative. Only nuclear receptor ligands (58 compounds) and transporters or pumps (32 compounds) were sampled deeply enough for their lack of hits to be interpretable, and neither yielded one.

### Top hit compounds induce tumor clearance across multiple progressor models

To prioritize compounds for further characterization, we re-screened high-potential compounds, including the six hits common to both CRC and BC models and a few others belonging to the enriched MOA classes, in additional progressor CRC zebrafish xenograft models (human SW620, and murine MC38 and CT26 cell lines, **Supplementary Figure 2**). As multiple drugs showed a consistent increase in tumor clearance across all models, we focused on two of the six common hits - Tiotropium Bromide (**Figure 3A**), a long-acting M3R antagonist, and Amrinone (**Figure 3B**), a PDE3 inhibitor - based on the strong enrichment of their respective MOA classes in the screen.

**Figure 3:**
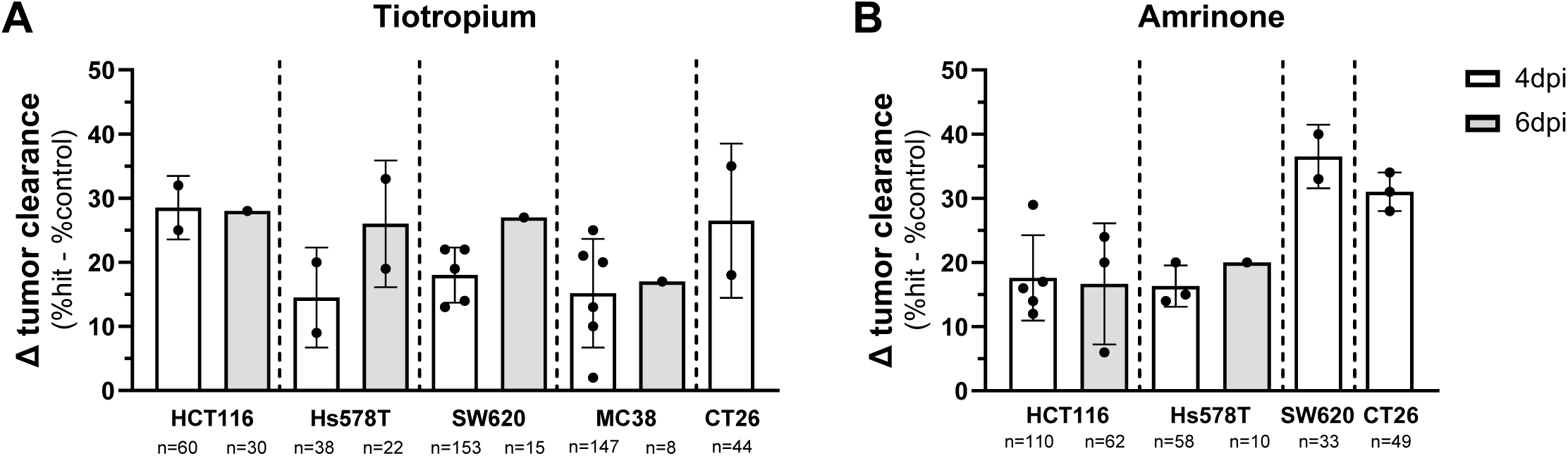
Top hit compounds induce tumor clearance in multiple progressor models. Induced tumor clearance of Tiotropium **(A)** and Amrinone **(B)** in multiple progressor zebrafish xenograft models at 4 and 6 dpi, quantified as the difference in tumor clearance between treated and control conditions. Tumor clearance is the percentage of fish with no tumor at the end of the assay. Graphs show mean ± STD, each dot represents one independent experiment, and n shows the total number of fish per condition.

To confirm that the clearance effects were not due to direct tumor-autonomous cytotoxicity, we performed in vitro viability assays with both lead candidates. Immunocytochemistry of HCT116 and CT26 cells exposed to 10 μM Tiotropium or Amrinone for 4 days, i.e., under the same treatment conditions used in the primary screen, showed no differences in apoptosis induction compared to DMSO controls (**Supplementary Figure 3**). Additionally, cell viability assays with increasing concentrations of Tiotropium did not impact the viability of any of the tested progressor cell lines (**Supplementary Figure 4**).

Collectively, these results reinforce the idea that the pro-clearance activity of these compounds is attributable to non-cell autonomous, immune-modulatory mechanisms rather than cancer cell-intrinsic vulnerabilities.

### Tiotropium and Amrinone promote anti-tumoral innate immunity

To determine whether the lead hit compounds modulate the innate TME, we performed immune profiling of progressor xenografts generated in myeloid-labeling transgenic reporter zebrafish lines (**Figure 4**, **Supplementary Figure 5**). Representative confocal images of CT26 xenograft tumors with DAPI counterstaining (**Figure 4A-C**) at 4 dpi illustrate the distribution of neutrophils (**Figure 4A’-C**’) and macrophages (**Figure 4A’’-C’’**) in control, Tiotropium-treated, and Amrinone-treated conditions.

**Figure 4:**
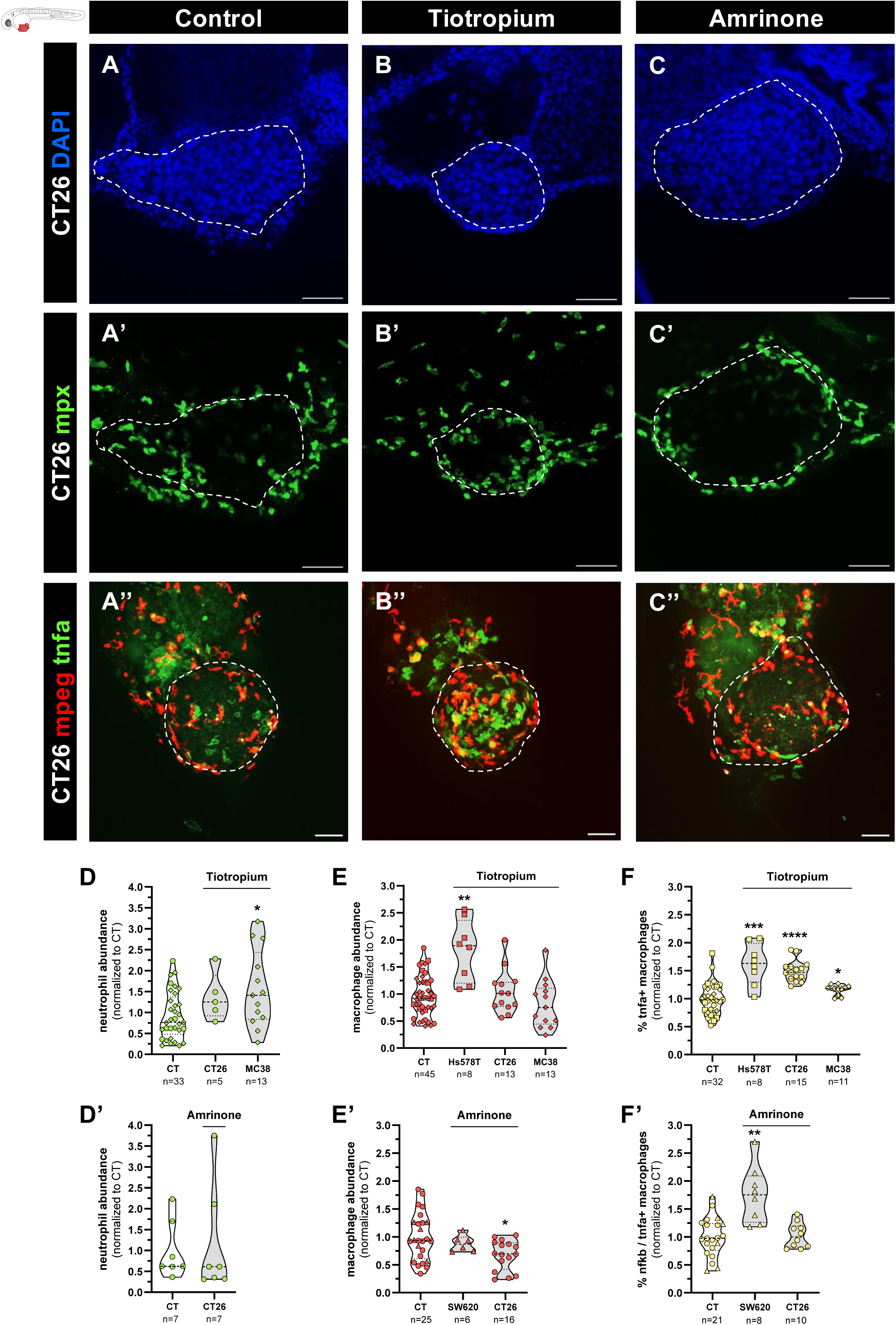
Selected hit compounds promote anti-tumoral innate immunity across multiple xenograft models. (A–C) Representative confocal images of CT26 xenografts at 4 dpi in control (A), Tiotropium-treated (B), and Amrinone-treated (C) conditions. White dashed lines outline the tumor. Row 1 (A–C): DAPI counterstaining labels cell nuclei in blue. Row 2 (A’–C’): neutrophils are labelled in green [Tg(mpx:GFP)]. Row 3 (A’’–C’’): macrophages are labelled in red and TNFα-expressing cells in green [Tg(mpeg1:mCherry-F; tnfa:eGFP-F)]. All images are oriented anterior to the left, posterior to the right, dorsal side up, and ventral side down. Scale bars represent 50 μm. **(D-E, D’-E’)** Quantification of neutrophil (D-D’) and macrophage (E-E’) abundance at the tumor site in Tiotropium-treated (D, E) and Amrinone-treated (D’, E’) xenografts across multiple progressor models, calculated as the ratio between the total number of innate cells and the tumor area of each xenograft, normalized to the control condition. **(F, F’)** Quantification of the percentage of inflammatory (NFκB/TNFα-positive) macrophages, normalised to the control. Violin plots show the interquartile range and the median. Each dot represents one zebrafish xenograft, and the total number analyzed (n) is indicated below each condition. According to data normality, comparisons between control and treated conditions were performed using the Welch’s t-test or the Mann-Whitney U test (*P ≤ 0.05, **P ≤ 0.01, ***P ≤ 0.001, ****P ≤ 0.0001).

Quantification across multiple progressor models revealed that both compounds variably modulate myeloid cell recruitment to the tumor site, with Tiotropium significantly increasing neutrophil abundance in MC38 xenografts (**Figure 4D**) and macrophage abundance in Hs578T xenografts (**Figure 4E**), while Amrinone decreased macrophage abundance in CT26 xenografts (**Figure 4E’**). However, as myeloid presence in the TME is not necessarily associated with tumoricidal activity, we focused on the polarization state of macrophages, which more reliably determines their pro/anti-tumoral orientation^43,44^. Notably, Tiotropium significantly expanded the proportion of TNFα-positive macrophages in HCT116, CT26, and MC38 xenografts (**Figure 4F**), and Amrinone elevated the proportion of NFκB-positive macrophages in SW620 xenografts (**Figure 4F**’). Of note, neither compound altered total myeloid cell numbers in non-tumor-bearing larvae (**Supplementary Figure 6**), suggesting that compound effects are TME-specific and do not reflect systemic perturbation of innate immune homeostasis.

In summary, Tiotropium and Amrinone promote anti-tumoral innate immunity mainly by reprogramming macrophages toward a pro-inflammatory state, which likely underlies the enhanced tumor clearance observed across models, and supports their potential as immunotherapies.

### Tiotropium promotes macrophage phagocytosis and requires macrophages for tumor clearance

To confirm that hit-induced macrophage polarization towards a pro-inflammatory state is reflected in increased anti-tumoral activity, we assessed phagocytosis in MC38 xenografts (**Figure 5A-F**). Consistently, Tiotropium treatment significantly increased both the absolute number and percentage of phagocytic macrophages at the tumor site at 4 dpi (**Figure 5E,F**). This indicates that Tiotropium promotes macrophage tumoricidal function, even in the absence of significant changes in total macrophage abundance.

**Figure 5:**
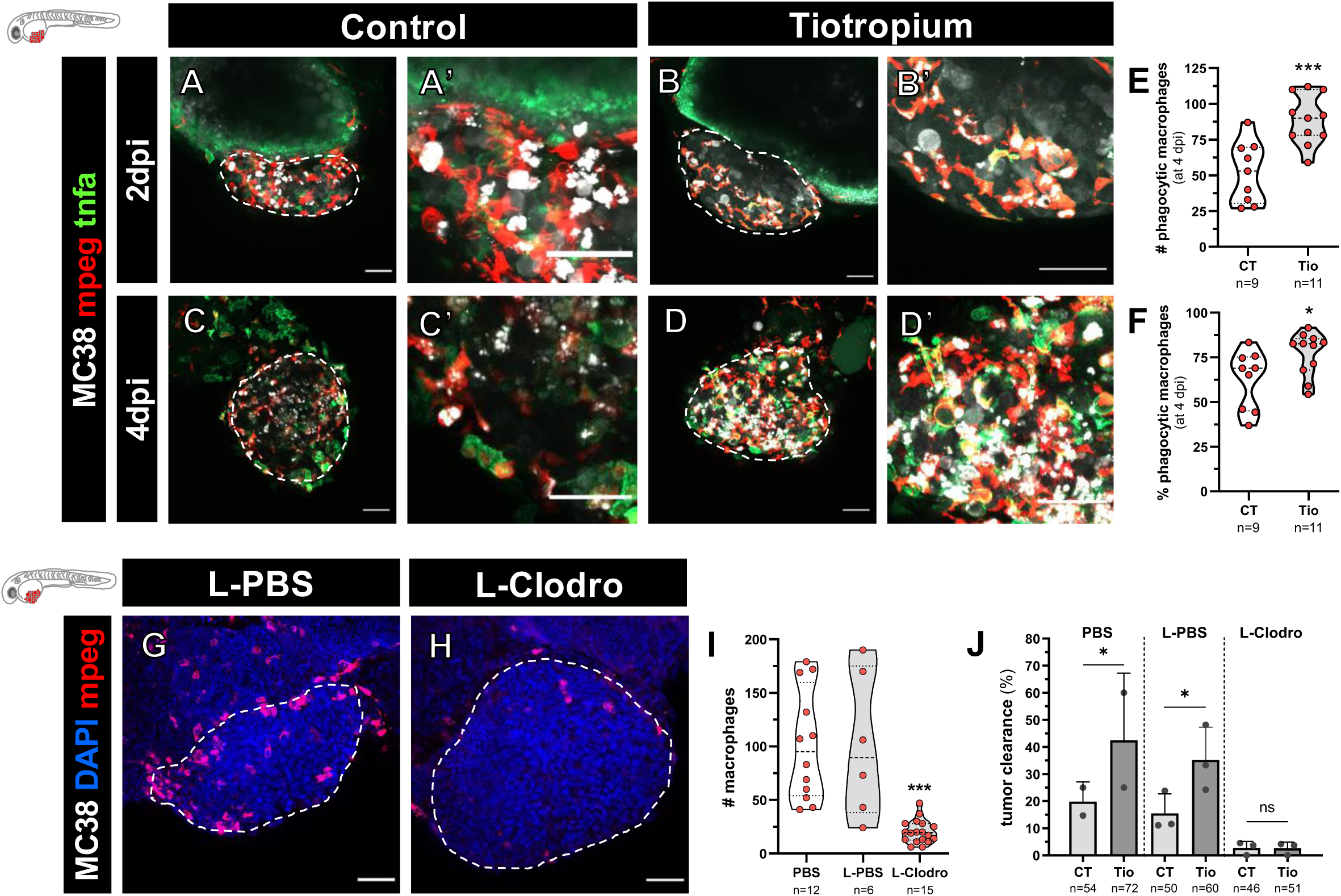
Tiotropium promotes macrophage phagocytosis and requires macrophages for tumor clearance. (A-D) Representative confocal images of MC38 xenografts at 2 and 4 dpi in control (A, C), and Tiotropium-treated (B, D) conditions. MC38 cells are presented in white, and dashed lines outline the tumor. Macrophages are labelled in red and TNFα-expressing cells in green [Tg(mpeg1:mCherry-F; tnfa:eGFP-F)]. All images are oriented anterior to the left, posterior to the right, dorsal side up, and ventral side down. Scale bars represent 50 μm. **(E,F)** Quantification of phagocytic macrophages at the tumor site in total number per xenograft (E) and percentage (F) in each condition. Violin plots show the interquartile range and the median. Each dot represents one zebrafish xenograft, and the total number analyzed (n) is indicated below each condition. Data were analyzed using the Welch’s t-test (*P≤ 0.05, ***P ≤ 0.001). **(G, H)** Representative confocal images in MC38 xenografts at 4 dpi in L-PBS (G) and L-Clodro (H) conditions. Macrophages were pharmacologically depleted by resuspending tumor cells in clodronate-encapsulated liposomes (L-Clodro) prior to injection. L-PBS-encapsulated liposomes (L-PBS) served as the vehicle control. Tumors are outlined in white, macrophages are labelled in red [Tg(mpeg1:mCherry-F)], and nuclei are stained with DAPI in blue. All images are oriented anterior to the left, posterior to the right, dorsal side up, and ventral side down. Scale bars represent 50 μm. **(I)** Quantification of total macrophage number per xenograft across PBS, L-PBS, and L-Clodro conditions. Violin plots show the interquartile range and the median. Each dot represents one zebrafish xenograft and the total number analyzed (n) is indicated below each condition. Data were analyzed using the Mann-Whitney U test (***P ≤ 0.001). **(J)** Tumour clearance percentage of control and Tiotropium-treated xenografts across conditions. Graph shows mean ± S.D. and each dot represents one independent experiment. The total number of xenografts analyzed per condition (n) is indicated. Data were analyzed using Fisher’s exact test (ns ≥ 0.05, *P ≤ 0.05).

To directly test whether these myeloid cells are required for Tiotropium-induced tumor clearance, we pharmacologically depleted macrophages with clodronate-encapsulated liposomes (L-Clodronate) prior to injection, with L-PBS-encapsulated liposomes serving as the vehicle control (**Figure 5G-J**). Successful local macrophage ablation was confirmed by quantification of total macrophages at the tumor site (**Figure 5I**). In the absence of macrophages, Tiotropium completely failed to induce tumor clearance, in stark contrast to PBS and L-PBS conditions, where it behaved as expected (**Figure 5J**). Together, these results establish macrophages as essential effectors of Tiotropium-induced tumor clearance, with their enhanced phagocytic and pro-inflammatory reprogramming likely underlying the observed anti-tumor response.

### Cholinergic neuronal projections are in close contact with SW620 tumors

Since Tiotropium is a muscarinic receptor antagonist, we asked whether acetylcholine (ACh)-mediated signaling could underlie its pro-clearance activity and sought to identify its likely source within the TME. ACh in tumors can originate from two sources: parasympathetic neuronal projections or the non-neuronal cholinergic system (NNCS), comprising tumor, stromal, and immune cells^45,46^. To analyze the former, we generated SW620 xenografts in the Tg(*chata:GFP*) cholinergic neuron reporter line. Confocal imaging at 4 dpi revealed extensive cholinergic neuronal architecture in the surrounding host tissue, with neuronal projections in close contact with the tumor mass (**Figure 6**). Further work is necessary to properly explore the mediators of cholinergic tone within the TME.

**Figure 6:**
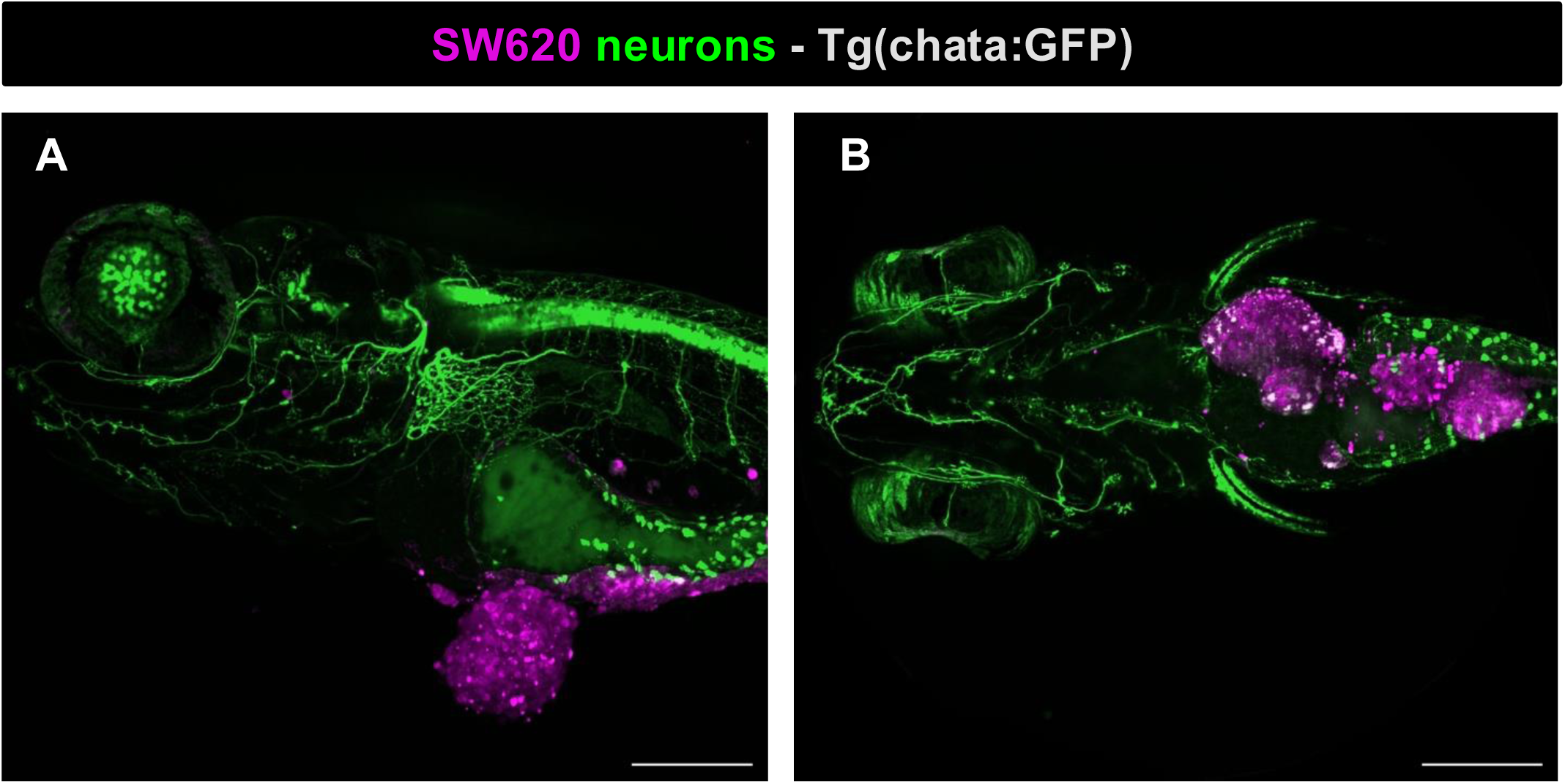
Cholinergic neuronal projections in SW620 xenografts. Representative confocal images of SW620 xenografts at 4 dpi generated in the *Tg(chata:GFP)* cholinergic neuron reporter line, shown in lateral **(A)** and dorsal **(B)** views. Cholinergic neurons are labeled in green and tumor cells in magenta. Scale bars represent 200 µm.

### Tiotropium shows efficacy as monotherapy and synergizes with ICB in a mouse tumor model

The final goal of our phenotypic screen was to identify innate-modulating compounds capable of switching the TME from suppressive to reactive, and that can then be translated to mammalian models and ultimately human patients. For this, we tested Tiotropium and Amrinone in the preclinical syngeneic CT26 CRC mouse model in the BALB/c background. In this model anti-PD-1 monotherapy shows limited efficacy, making it an ideal model to test synergy between our hit compounds with adaptive ICB^47^.

Initial tolerability and dose-response experiments tested Tiotropium at 1, 10, and 50 mg/kg and Amrinone at 1, 10, and 20 mg/kg as monotherapies, administered intratumorally (**Supplementary Figure 7A**). Amrinone showed no significant changes in animal body weight, a proxy for tolerability, nor in tumor growth, at any dose tested (**Supplementary Figure 7B,C**). In contrast, Tiotropium showed a trend toward tumor reduction, though accompanied by dose-dependent toxicity, with weight variation exceeding 10% at doses of 10 and 50 mg/kg. Based on these results, Tiotropium was selected as the sole focus for subsequent ICB combination experiments using doses below 1 mg/kg.

CT26 tumor-bearing mice were treated with intratumoral Tiotropium (0.1 or 0.5 mg/kg), alone or in combination with intravenous anti-PD-1 antibody (**Figure 7A**). All treatment groups significantly reduced tumor volume and improved survival relative to vehicle-treated controls (**Figure 7B,C**). Both doses of Tiotropium produced comparable tumor volume reduction and survival benefit as monotherapy and were not significantly different from anti-PD-1 alone in either readout.

**Figure 7:**
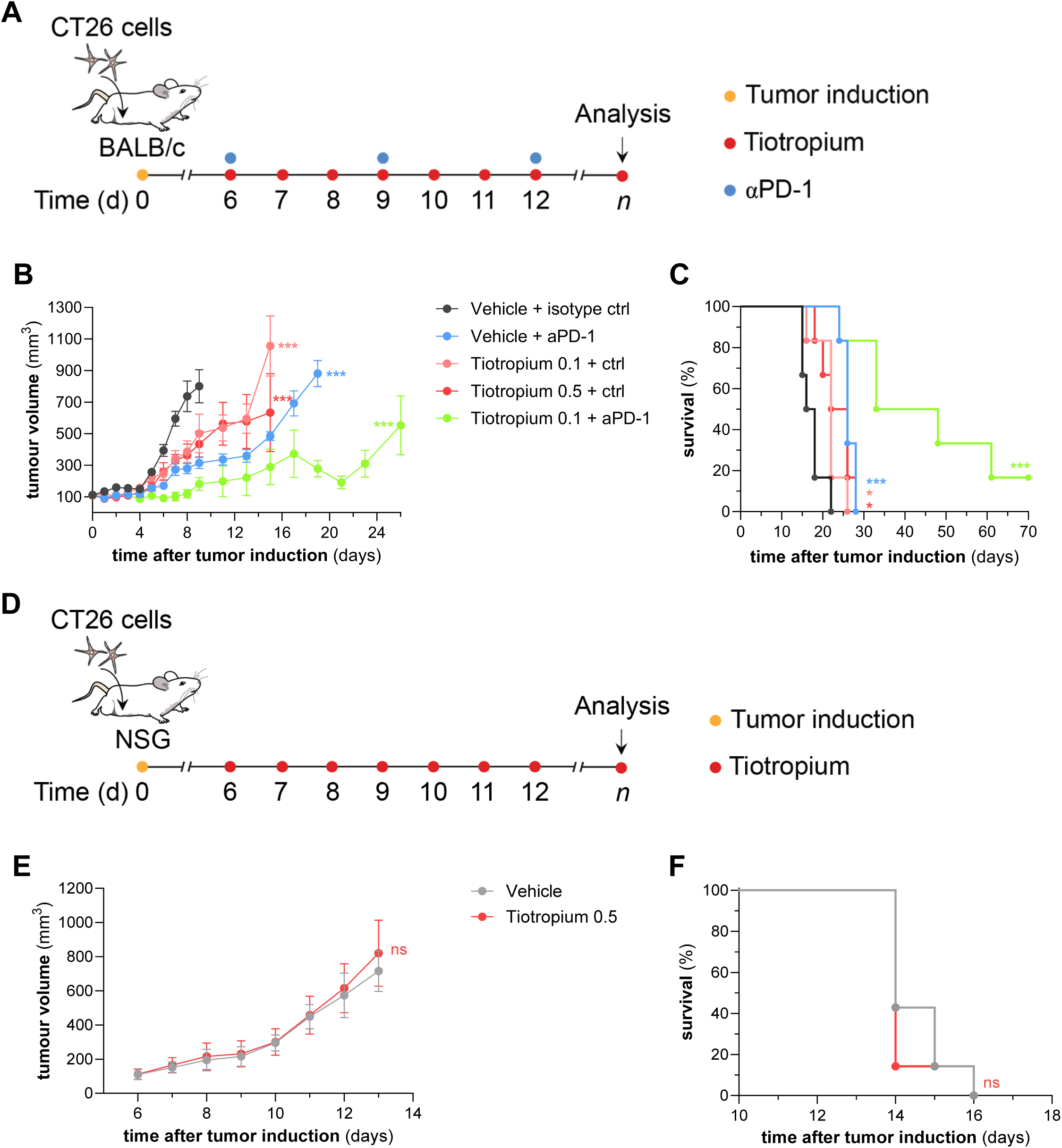
Tiotropium shows efficacy as a monotherapy and synergizes with ICB in a CT26 mouse tumor model. **(A)** Experimental design for testing synergy between Tiotropium and ICB. CT26 cells were inoculated subcutaneously into the abdomens of BALB/c mice. Treatments began once tumor volume reached 100-200 mm³. αPD-1: 200ug/mouse (IV); Tiotropium: 0.1 or 0.5 mg/kg (IT). n = 6 for each treatment group. IV, intravenous; IT, intratumoral. (**B)** Tumor volume curves. Treatment groups were compared with the vehicle + isotype control on day 8 of treatment (day 13 after tumor induction) using Two-way RM ANOVA. Data are presented as mean ± STD. Vehicle vs. Tiotropium 0.1 mg/kg *p*<0.0001, vehicle vs. Tiotropium 0.5 mg/kg *p*<0.0001, vehicle vs. αPD-1 *p*<0.0001, vehicle vs. combination *p*<0.0001; Tiotropium 0.1 vs. 0.5 mg/kg *p*=0.9593; αPD-1 vs. combination *p*<0.0001; Tiotropium 0.1 vs. combination *p*<0.0001; Tiotropium 0.5 vs. combination *p*<0.0001; αPD-1 vs. Tiotropium 0.5 *p*=0.2347; αPD-1 vs. Tiotropium 0.1 *p*=0.0114. (**C)** Survival curves. Treatment groups were compared with the vehicle + isotype control using the log-rank (Mantel-Cox) test. Vehicle vs. Tiotropium 0.1 mg/kg *p*=0.0282, vehicle vs. Tiotropium 0.5 mg/kg *p*=0.0156, vehicle vs. αPD-1 *p*=0.0006, vehicle vs. combination *p*=0.0006; Tiotropium 0.1 vs. 0.5 mg/kg *p*=0.4141; αPD-1 vs. combination *p*=0.0126; Tiotropium 0.1 vs. combination *p*=0.0021; Tiotropium 0.5 vs. combination *p*=0.0021; αPD-1 vs. Tiotropium 0.5 *p*=0.1255; αPD-1 vs. Tiotropium 0.1 *p*=0.1879. **(D)** Experimental design for determining if Tiotropium’s effect is immune-dependent. CT26 cells were inoculated subcutaneously into the abdomens of NSG mice. Treatment began once tumor volume reached 100-200 mm³. Tiotropium: 0.5 mg/kg (IT). n = 7 for each treatment group. **(E, F)** Tumor volume curves (E) and survival curves (F) for NSG experiments, analyzed as in (B) and (C), respectively. No significant differences were observed between the vehicle and the Tiotropium-treated groups.

Strikingly, the combination of Tiotropium 0.1 mg/kg with anti-PD-1 outperformed all monotherapy arms, producing the greatest tumor volume reduction and survival benefit across all treatment groups. These results were independently replicated in a second experiment (**Supplementary Figure 8**). The superiority of the combination over each agent alone argues that Tiotropium and anti-PD-1 engage complementary rather than overlapping immune mechanisms. Importantly, treatment was well tolerated across all arms, as confirmed by histopathological evaluation, which showed normal organ architecture with no treatment-related inflammatory, degenerative, necrotic, vascular, or proliferative lesions (**Supplementary Figure 9**).

To confirm Tiotropium’s immune-dependent efficacy, as observed in zebrafish, we made use of immunodeficient NOD scid gamma (NSG) mice, which lack T, B, and NK cells and present defective dendritic cell function and reduced macrophage phagocytic activity^48^ (**Figure 7D**). In stark contrast to CT26 tumors in immune-competent BALB/c mice, Tiotropium at 0.5 mg/kg failed to reduce tumor growth or extend survival in this context **(Figure 7E,F)**. Tiotropium-treated groups showed statistically significant but modest reductions in body weight relative to vehicle controls, remaining below the acceptable 10% threshold, with no treatment-related morbidity observed (**Supplementary Figure 10**).

### Tiotropium alters the myeloid and T cell compartments of ICB-treated tumors

To examine the immune changes accompanying the Tiotropium–anti-PD-1 combination, we repeated the experiment with Tiotropium at 0.1 mg/kg, harvested CT26 tumors on day 12 and profiled viable CD45⁺ leukocytes by spectral flow cytometry (**Supplementary Figure 11A**). Unsupervised clustering of 1,093,830 cells across 18 markers resolved 20 immune populations (**Figure 8A**, **Supplementary Figure 11B,C**). Overall immune composition was broadly similar between arms (**Figure 8B**, **Supplementary Figure 11D,E**).

**Figure 8:**
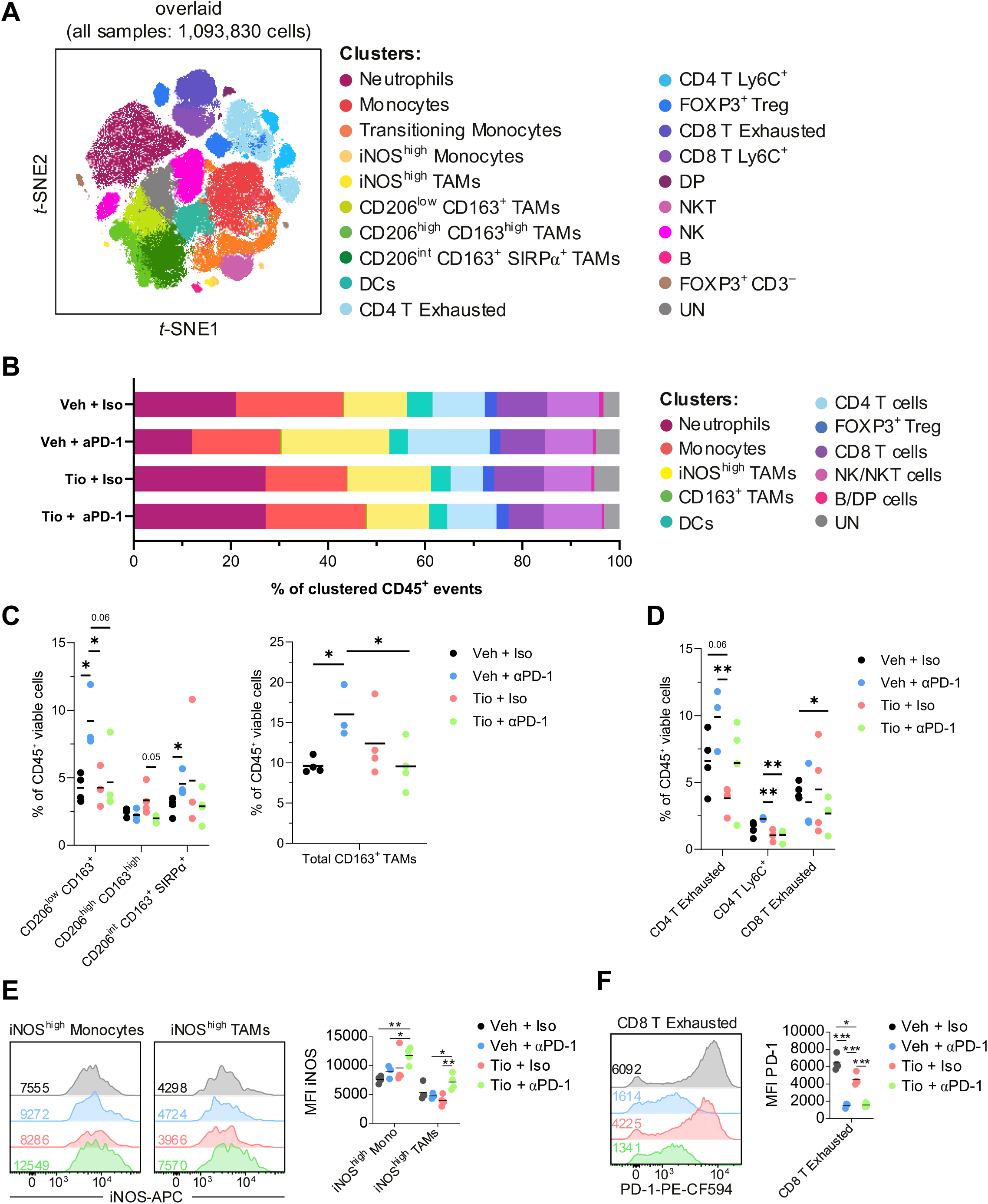
Tiotropium alters the myeloid and T cell compartments of CT26 tumors under checkpoint blockade. Tumors were harvested on day 12 from BALB/c mice treated with vehicle + isotype control (n = 4), vehicle + anti-PD-1 (n = 3), Tiotropium 0.1 mg/kg + isotype control (n = 4) or Tiotropium 0.1 mg/kg + anti-PD-1 (n = 4), and analyzed by spectral flow cytometry. **(A)** Unsupervised t-distributed stochastic neighbor embedding (t-SNE) of viable CD45⁺ cells from all samples (1,093,830 cells), showing the 20 clusters identified by FastPhenoGraph. (**B)** Immune composition per treatment, expressed as a percentage of clustered CD45⁺ events. Bars show the mean of per-animal percentages, with the 20 clusters grouped into 11 lineages. **(C)** Frequencies of the CD163⁺ TAM clusters (left) and their per-animal sum (right). **(D)** Frequencies of CD4 T exhausted, CD4 T Ly6C⁺, and CD8 T exhausted clusters. **(E)** iNOS expression in iNOS^high^ monocytes and iNOS^high^ TAMs, shown as representative histograms (left) and mean fluorescence intensity (MFI) across all animals (right). **E)** PD-1 expression in CD8 exhausted T cells, shown as representative histograms (left) and mean fluorescence intensity (MFI) across all animals (right). Horizontal lines in graphs indicate group means and each dot represents one animal. Data were analyzed using unpaired two-tailed *t*-tests (ns ≥ 0.05, *P ≤ 0.05, **P ≤ 0.01, ***P ≤ 0.001). TAMs: tumor-associated macrophages; DP: double-positive (CD4+CD8+) T cells; DCs: dendritic cells; Treg: regulatory T cells; NKT: natural killer T cells; NK: natural killer cells; UN: unassigned.

Within the myeloid compartment, anti-PD-1 monotherapy was associated with an expansion of CD163⁺ TAMs, a phenotype linked to immunosuppression, driven by the CD206^low^ CD163⁺ and CD206^int^ CD163⁺ SIRPα⁺ TAM clusters (**Figure 8C**). This expansion was not observed under combination treatment, where CD163⁺ TAM frequency returned to control levels. A comparable pattern was seen among CD4 T cells, with exhausted and Ly6C⁺ frequencies higher under anti-PD-1 monotherapy than in Tiotropium-treated animals, and a decrease in CD8 exhausted T cells under combination treatment versus controls (**Figure 8D**).

Regarding myeloid activation, combination treatment increased iNOS mean fluorescence intensity (MFI) in monocytes relative to both vehicle and anti-PD-1 monotherapy, and in TAMs relative to both monotherapies (**Figure 8E**), with correspondingly higher frequencies of iNOS^high^ monocytes and TAMs than under Tiotropium alone (**Supplementary Figure 11F**). Additionally, Tiotropium monotherapy reduced PD-1 expression in CD8 exhausted T cells relative to controls, suggesting a decrease in exhaustion.

Together, these data indicate that the Tiotropium–anti-PD-1 combination is associated with a myeloid compartment skewed toward iNOS expression and away from CD163⁺ TAM accumulation, accompanied by lower frequencies of exhausted T cells, relative to anti-PD-1 alone. Overall, Tiotropium’s efficacy as monotherapy, its synergy with anti-PD-1, and its loss of activity in immunodeficient mice establish it as a translationally relevant innate immune modulator and support its repurposing as an agent capable of sensitizing cold tumors to ICB.

## Discussion

Here, we report the first *in vivo* phenotypic drug screen specifically designed to identify innate immune modulators in cancer. Using zebrafish xenografts to screen an FDA-approved library in a two-round design, we identified 23 compounds capable of boosting innate tumor clearance across two cancer models. This yielded hit rates of 2.56% and 3.84% for HCT116 and Hs578T respectively, comparable with other large zebrafish phenotypic screens^49–51^. The zebrafish xenograft model is uniquely suited for this screen as it builds on the well-established strengths of zebrafish PDD, such as scalability, optical transparency, and whole–organism pharmacology^17^, while adding the crosstalk between tumor cells and the innate immune system. Furthermore, the use of progressor models with a reproducible low-clearance baseline, short assay window, and a stringent selection criterion of tumor clearance collectively filters against tumor-autonomous drug effects. Six of the 23 confirmed hits showing activity across both CRC and BC models is particularly noteworthy, as it suggests the drugs target conserved innate immune regulatory mechanisms rather than cancer type-specific biology. Together, these features establish an innate immunity–focused screening platform that can be readily applied to additional tumor types and drug libraries.

### Macrophage reprogramming underlies the clearance phenotype

To better understand how the screened compounds enhance tumor clearance, we further characterized two representative hits, Tiotropium and Amrinone, from the six that showed activity across both CRC and BC models. It is well established that innate immunity, namely macrophages and neutrophils, is a key modulator of tumor control at the larval stage of zebrafish xenografts^35,52^. In fact, progressor tumors can predominantly suppress the clearance of co–injected regressors in polyclonal xenografts, demonstrating the strongly immunosuppressive microenvironment of this model^35^.

Immune profiling of Tiotropium– and Amrinone–treated xenografts in myeloid-reporter lines showed that both compounds variably modulate macrophage and neutrophil recruitment to the tumor site but converge on a macrophage-activating effect, reflected in the expansion of inflammatory TNFα- or NFκB-positive tumor-associated macrophages (TAMs) across multiple xenograft models. TAMs are among the most abundant immune populations in the TME of most solid cancers and are predominantly skewed toward an M2-like immunosuppressive, pro-tumoral phenotype^53,54^. However, TAM abundance alone is not a reliable surrogate for prognosis, as its clinical impact is highly context-dependent and varies widely across tumor types^55,56^. Macrophage activation state, often inferred from the balance between pro-inflammatory M1-like and anti-inflammatory M2-like cells, has been described as a much more informative proxy for survival and therapy response^57,58^, including in CRC and BC^59,60^. Accordingly, therapeutic strategies to repolarize TAMs are being actively explored as a means to overcome immunosuppression and therapy resistance^57,61^. In further support of a functional tumoricidal response, Tiotropium significantly increased the number and proportion of tumor cell-phagocytic macrophages, consistent with the link between inflammatory polarization and enhanced tumor cell killing^62^. The direct requirement for macrophages in Tiotropium’s effect was established by pharmacological depletion, which revealed a complete abrogation of tumor clearance in the absence of this myeloid population.

The heterogeneity of responses across models, namely in myeloid recruitment, likely reflects differences in the baseline immunosuppressive landscape of each progressor cell line^35^. This variation in response patterns that nevertheless lead to a tumor-clearing phenotypic output, suggests that Tiotropium and Amrinone act on a generalized immune regulatory axis. Importantly, neither Tiotropium nor Amrinone showed direct cancer cell toxicity *in vitro* or altered total myeloid numbers in non–tumor–bearing larvae, pointing to TME-specific immune effects. Together, these findings position anti-tumor macrophage repolarization, rather than simple modulation of myeloid recruitment, as the primary shared effector mechanism underlying the pro-clearance activity of the lead hit compounds.

From a translational perspective, these observations support macrophage reprogramming, rather than depletion, as a more desirable therapeutic strategy. While TAM-depleting approaches, such as CSF1R inhibition, can dampen their immunosuppressive activity, they will also impair antigen phagocytosis, presentation, and cytokine production, all crucial for effective anti-tumoral immunity^63^. This is of particular importance in the context of immunotherapy, where myeloid-rich tumors that often exhibit primary resistance to ICB, a major unmet clinical challenge, could benefit from strategies capable of converting a “cold” TME into a responsive one^4,64,65^. Most CRC patients, particularly those with microsatellite stability (MSS), fall into this category, presenting M2-dominated and low T-cell-infiltrated cold tumors that derive minimal benefit from single-agent ICB^66,67^.

### Innate priming enhances checkpoint blockade

Motivated by this, we asked whether our lead hit compounds could act as innate immune primers to improve the efficacy of adaptive-targeting ICB. To model this setting experimentally, we tested Tiotropium and Amrinone in a CT26 murine CRC model that recapitulates key features of MSS CRC, including a TAM–dominated immunosuppressive TME and moderate responsiveness to anti–PD–1 monotherapy^68^. Strikingly, the combination of Tiotropium 0.1 mg/kg with anti-PD-1 outperformed all monotherapy groups, yielding the greatest benefit in terms of both tumor reduction and survival. This argues against simple additive effects and suggests instead that Tiotropium and ICB engage complementary immune mechanisms, and that lower, potentially better-tolerated doses of each agent may suffice in combination. This is in line with the emerging view that combination strategies are key avenues for overcoming primary resistance to ICB, with examples such as oleuropein-driven myeloid reprogramming, innate-priming STING agonists, and cytokine-based regimens like IL–21 showing robust synergy with anti–PD–1 in preclinical models^69–72^. The loss of Tiotropium efficacy in immunodeficient NSG mice provides additional evidence that its anti-tumor activity is immune-dependent, consistent with the macrophage dependence observed in zebrafish.

To identify immune changes accompanying this synergy, we profiled CT26 tumors by spectral flow cytometry. Anti-PD-1 monotherapy expanded CD163⁺ TAMs, a phenotype associated with immunosuppression and whose abundance correlates with poor prognosis across solid tumors, including CRC^55,73^. Myeloid expansion following ICB has been described as a compensatory resistance mechanism, whereby relief of T-cell inhibition is offset by recruitment or polarization of suppressive myeloid populations that constrain the response^74,75^. CD163⁺ TAM expansion was absent under combination treatment, suggesting that part of the benefit of combining Tiotropium may lie in limiting a myeloid counter-response to ICB, rather than simply augmenting T-cell activation. Accordingly, the combination was also associated with higher iNOS expression in monocytes and TAMs, consistent with a shift toward classically activated M1-like macrophages^76^. Among T cells, exhausted CD4 and CD8 populations were less frequent in Tiotropium-containing arms than under anti-PD-1 monotherapy.

Altogether, our data support a model in which Tiotropium’s myeloid conditioning sensitizes cold tumors to subsequent anti-PD-1-mediated T-cell reinvigoration, resulting in complementary engagement of both arms of the immune system to induce durable tumor control and overcome therapy resistance.

### An FDA-approved drug library enables mechanistic inference

The use of an FDA-approved library not only accelerates drug development and clinical translation, thanks to established safety profiles and pharmacokinetics, but it also facilitates mechanistic studies, as all compounds have known targets and indications. As such, we performed an MOA enrichment analysis to identify drug classes overrepresented among the 23 hits relative to the full library, revealing MR antagonists as the only significantly enriched class, with PDE inhibitors showing a similar trend. Notably, the hit list also contained COX–inhibiting NSAIDs, such as Ketoprofen^77,78^ and Etodolac^79,80^, a drug class already described as myeloid-modulating^81^, validating our original screen rationale. The recovery of this established immunomodulatory drug class serves as a screen positive control and underscores the idea that the identified enriched MOA classes capture clinically relevant axes of immune regulation rather than incidental pharmacology.

### Cyclic nucleotide signaling in immune modulation

Two PDE inhibitors scored in our screen, the PDE3-specific Amrinone and the non-selective Pentoxifylline, a class that trended toward enrichment. Blocking PDE-mediated catalytic breakdown to elevate intracellular cyclic nucleotides is a well-established way to modulate immune cell functions. In monocytic cells, most studies describe PDE inhibition as having anti-inflammatory effects, as increased cAMP and cGMP levels generally dampen inflammatory cytokine production and shift transcriptional programs away from NFκB–driven TNFα responses^82–84^. However, PDE isoforms matter as they modulate partially distinct cAMP pools, with PDE3 contributing more to basal cAMP degradation and PDE4 becoming the dominant regulator under conditions of elevated, stimulus–induced cAMP^85^. Conversely, in oncological contexts, increased PDE activity has been shown to contribute to tumor–driven immune evasion: PDE4 overexpression reduces T-cell activation and promotes TAM M2-like polarization^86–88^, while PDE5 inhibition can limit myeloid–derived suppressor cell (MDSC) function and restore T–cell–mediated tumor control^89,90^. Our finding that the PDE3–targeting inhibitor Amrinone increases innate tumor clearance by inducing a more activated, pro–inflammatory macrophage state further reflects how the impact of decreased cyclic nucleotide signaling can be highly isoform–specific and context–dependent, ranging from restraint of excessive systemic inflammation to relief of local immune suppression within the TME. Further work is required to understand the precise mechanisms by which PDE inhibition promotes innate immune activation in this experimental context.

### Neuro-immune signature among hits

A striking outcome of our unbiased screen was the convergence of hit compounds on drugs targeting autonomic neurotransmission, such as muscarinic, dopaminergic, and adrenergic modulators, which accounted for 9 of the 23 hits (39.1%) while representing only 17.3% of the full library. This aligns with the growing recognition that the nervous system is an active regulator of the TME. The emerging field of cancer neuroscience shows that sympathetic, parasympathetic, and sensory fibers innervate most solid tumors, which in turn promote neuro-immune crosstalk that can shape disease progression and therapy response^91,92^. Additionally, many immune cell types express adrenergic, cholinergic, and dopaminergic receptors, and their engagement reshapes cytokine output, antigen presentation, and effector function^93,94^. In the context of our model, this opens two possibilities: the hit compounds may act indirectly by exerting effects on neural circuits that will in turn affect the TME, or directly, by targeting the neurotransmitter receptors expressed by myeloid cells.

### A dopaminergic trend without a shared direction

Dopaminergic signaling is increasingly recognized as a neuro–immune axis in cancer. Peripheral dopamine acts on immune cell-expressed D-like receptors (DRD) to modulate cytokine production, antigen presentation, and cytotoxic activity in a dose– and receptor type–dependent manner^95^. In several tumor models, dopamine has been shown to restrain MDSC activity and restore NK and T-cell cytotoxic responses^96,97^. Conversely, DRD2 antagonism can reprogram TAMs toward an inflammatory state^98,99^. Our screen recovered three compounds acting on dopaminergic signaling. Two are used clinically for Parkinson’s disease: the aromatic L-amino acid decarboxylase (AADC) inhibitor Carbidopa, which limits dopamine synthesis, and the catechol-O-methyltransferase (COMT) inhibitor Tolcapone, which slows dopamine degradation. The third is the antihypertensive Methyldopa, which depletes central catecholamines through false neurotransmitter formation. That these three compounds perturb catecholamine handling in different directions suggests that the relevant variable for immune modulation in this context may be turnover rather than absolute dopaminergic tone level, which would be consistent with the opposing effects reported for dopamine itself and for DRD2 blockade. Although we did not mechanistically pursue these hits, their independent identification points to dopaminergic modulation as an under-explored therapeutic axis in cancer that warrants dedicated follow-up.

Interestingly, Tolcapone offers an additional mechanistic lead that does not require dopamine. In a glioma model, COMT inhibition has been shown to cause mitochondrial dysfunction and double-stranded RNA leakage in tumor cells, activating a type I interferon response that stimulates microglia-mediated phagocytosis^100^. Whether Tolcapone acts through this route or through dopaminergic signaling in our model remains to be tested.

### Opposing adrenergic modulators promote clearance

The screen also uncovered two adrenergic modulators, the α2–adrenergic agonist Methyldopa Sesquihydrate and the β–blocker Metoprolol Tartrate, although the adrenergic system as a whole was not enriched. Sympathetic adrenergic signaling is nonetheless the most extensively characterized ANS arm in this context. In fact, β–adrenergic antagonism has emerged as an immunomodulatory strategy, with preclinical and clinical data indicating that β–blockers can relieve stress–induced immunosuppression and enhance T–cell function^101,102^. Notably, β–blocker use has been associated with survival benefits across several cancers^103–105^ and specifically with improved outcomes under immunotherapy, prompting a phase I trial of propranolol with pembrolizumab^106,107^. Likewise, α2–adrenergic receptor activation has recently been shown to upregulate both innate and adaptive anti-tumor immune responses, including reprogramming of TAMs toward a pro-inflammatory phenotype^108,109^. In autoimmune settings, Methyldopa was specifically demonstrated to modulate antigen presentation by blocking MHC class II interactions and dampening T–cell responses^110^. That two compounds with opposing adrenergic actions both promote clearance, in a system showing no overall enrichment, is consistent with the context-dependent, “yin-yang” effects attributed to sympathetic signaling in tumors^111^, and argues against a simple directional model for this arm.

### Cholinergic signaling as a dominant axis in the screen

MR antagonists represent the most enriched MOA class in our screen, with five confirmed hits - Tiotropium, Dicyclomine, Disopyramide, Scopolamine, and Glycopyrrolate - in one or both xenograft models. The fact that these five compounds display different MR subtype selectivity profiles, yet converge on a pro-clearance outcome, argues for a class effect of muscarinic blockade rather than a receptor subtype-specific mechanism. Acetylcholine, an essential neurotransmitter and the canonical ligand for MRs, can be produced not only by neurons but also by a non-neuronal cholinergic system (NNCS), comprising tumor, stromal, and immune cells, all of which can express muscarinic receptors and engage in autocrine or paracrine signaling^46^. The immunological effects of MR antagonism are, therefore, complex and context-dependent, shaped by the opposing G-protein coupling profiles of the five MR subtypes: M1R, M3R, and M5R signal through Gq/11 to activate phospholipase C and downstream calcium mobilization, while M2R and M4R couple to Gi/o to suppress cAMP^112^. In a model of acute inflammation, this complexity manifests as subtype-specific opposing effects: selective M1R blockade reduces TNFα and improves survival in septic shock, whereas M2R blockade promotes M1-like macrophage polarization and worsens outcome^113^. In zebrafish larvae specifically, pharmacological inhibition of MRs increases neutrophil recruitment and pro-inflammatory cytokine expression through downregulation of arginases 1/2^114^. This picture becomes even more complex by evidence from cancer models, where the TME can markedly influence cholinergic tone, which has in turn been linked to both tumor-promoting and tumor-suppressive outcomes^45^. M2R and M4R signaling in macrophages is inherently immunosuppressive, as the resulting reduction in intracellular cAMP removes the inhibitory constraint on NFκB-driven pro-inflammatory activation^115^. In cancer models, M4R on TAMs drives MAPK/AKT-mediated M2 polarization and upregulation of immune checkpoint markers in prostate cancer^116^, while M1R-dependent parasympathetic input suppresses pancreatic tumorigenesis by limiting myeloid expansion and cancer stemness^117^. Reconciling this data and invoking the principle that macrophage responses are exquisitely sensitive to their pre-existing polarization state and transcriptional history^118^, we hypothesize that in the chronically M2-skewed TME, where constitutive cholinergic signaling through M2R and M4R sustains immunosuppression, broad MR antagonism would be expected to relieve this suppressive brake, whereas the directionally opposite effect would be expected in an acutely inflamed, M1-polarised context.

### Tiotropium working model

Tiotropium, our main hit, is a long-acting muscarinic antagonist approved for the treatment of chronic obstructive pulmonary disease (COPD), where it acts by blocking M1R- and M3R-mediated bronchoconstriction and mucus secretion in the airway. Although Tiotropium binds M1R, M2R, and M3R with equal affinity in the lung, it shows kinetically selective, prolonged antagonism of M1R in the nerve ganglia and of M3R in smooth muscle and mucous glands, with rapid dissociation from M2R on postganglionic nerve fibers^119^. Beyond its canonical function, Tiotropium has documented immunomodulatory properties in multiple contexts: it inhibits TNFα-dependent alveolar macrophage-mediated neutrophil chemotaxis^120^, reduces M2 macrophage development in allergic inflammation^121^, inhibits ACh-driven generation of pro-inflammatory microparticles^122^, and has clinical evidence of anti-inflammatory effects in COPD^123^. In the context of cancer specifically, M3R has emerged as a key mediator of tumor immune evasion. In gastric cancer, M3R activation by tumor-derived ACh supports tumorigenesis and stemness by driving EGFR/ERK/AKT signaling^124^. In CRC, tumor–derived ACh acting on M3R upregulates PD-L1/2 expression in immune cells^125^ and pharmacological blockade has been shown to improve anti-tumor immune responses in *in vivo* models^126,127^.

Based on this body of evidence, we propose three non-mutually exclusive hypotheses for how Tiotropium and broader MR antagonism promote macrophage pro-inflammatory reprogramming and tumor clearance. The first is that tumor/TME-derived ACh acting through MRs on TAMs suppresses their pro-inflammatory potential, and that MR blockade relieves this NNCS-mediated brake. The second is that disrupting ACh–M3R/M4R signaling in tumor cells reduces the expression of immunosuppressive markers, thereby improving immune clearance. The third is that MR blockade may cause local ACh accumulation within the TME, shifting cholinergic signaling towards nicotinic receptors on macrophages, particularly α7nAChR, whose activation on TAMs has been shown to promote anti-tumor activity in CRC and enhance macrophage phagocytosis^128,129^.

The presence of cholinergic neuronal projections in the vicinity of SW620 xenograft tumors in the Tg(chata:GFP) line leaves both neurons and the NNCS as possible relevant ACh sources across all three hypotheses. Future work is necessary to resolve and distinguish the contributions of each of these mechanisms, such as quantification of TME ACh levels and identification of its prime source, genetic ablation or chemical blockade of specific MR subtypes, and nicotinic receptor perturbation experiments.

### Sampling depth constrains the enrichment analysis

The main limitation of our screen is sampling depth. Most MOA classes represented in the library contain too few compounds for any hit to be expected by chance, so the absence of hits in a given class carries little information about the biology of that target family. Crucially, class size in an approved-drug library is not arbitrary, as druggability has historically been determined by binding site architecture and ease of targeting^130^. As a result, approved drugs cluster into a small number of privileged families, mainly GPCRs, ion channels, kinases, and nuclear receptors^131^, and only a few hundred of the roughly 4,500 human genes estimated to be druggable have an approved drug against them^132^. The classes we could interpret here are therefore those that happen to be well represented in clinical pharmacology, which reflects the history of drug development rather than the relevance of a target to innate immunity. This approach renders whole target families invisible, regardless of their biological importance.

The analysis is further limited by what the annotation records and by what it can establish. Older drugs are not only more promiscuous, having emerged from an era of limited chemical diversity and no capacity to counterscreen for specificity, but also the best characterized, so their off-target activities are documented in a detail that recent approvals lack^133^. As such, these drugs often populate more than one MOA class, irrespective of how selective they actually are. More fundamentally, assigning a compound to a class does not establish which activity produces the phenotype. Disopyramide and Methyldopa each contribute to two classes, and attributing their pro-clearance effect to either would require targeted pharmacology that we did not perform. We therefore treat this analysis as hypothesis-generating. Its value was in directing attention to the cholinergic axis, and the mechanistic weight of this study rests on the experiments that follow rather than on the enrichment statistics themselves.

### Conclusions

In summary, this work presents the first *in vivo* phenotypic drug screen specifically designed to identify innate immune modulators in cancer, establishing the zebrafish xenograft model as a productive platform for this purpose. The screen yielded a clinically actionable hit list spanning diverse pharmacological classes and identified Tiotropium as a myeloid-reprogramming innate immune primer and proof-of-concept repurposing candidate that can sensitize cold tumors to checkpoint blockade. Beyond the individual compounds, the enrichment of autonomic nervous system-targeting drugs positions the neuro-immune interface as an underexplored but pharmacologically tractable axis for cancer immunotherapy that warrants dedicated investigation. Together, these findings highlight the untapped potential of *in vivo* phenotypic approaches to navigate the complexity of the tumor immune microenvironment and deliver immediately translatable candidates for clinical testing.

## Materials and Methods

### Cancer cell lines and culture

Human breast cancer cell line Hs578T was kindly provided by Mónica Bettencourt Dias Lab (Gulbenkian Institute for Molecular Medicine). Human colorectal cancer cell line SW620 was purchased from American Type Culture Collection (ATCC), whereas HCT116 was kindly provided by Ângela Relógio Lab (Charité Medical University of Berlin). Mouse colorectal cancer cell lines CT26 (ATCC) and MC38 (Kerafast) were kindly provided by Gonçalo Bernardes Lab (Gulbenkian Institute for Molecular Medicine).

Cells were cultured in Dulbecco’s Modified Eagle Medium (DMEM) High Glucose (Biowest), supplemented with 10% FBS (Sigma-Aldrich) and antibiotics (100 U ml−1 penicillin and 100 μg ml−1 streptomycin, Hyclone), maintained in a humidified atmosphere of 5% CO_2_ at 37 °C.

### *In vitro* cytotoxicity and viability assays

Sterile round coverslips (VWR, 13mm #1.5) were placed in 12-well plate wells and seeded with HCT116 or CT26 cells (3 x 10^4^ cells/well) in complete cell culture medium [DMEM supplemented with 10% fetal bovine serum (FBS) and 1% Penicillin-Streptomycin (P/S)]. For days 1 to 4 after seeding, DMSO or the drug (Tiotropium or Amrinone) was freshly diluted in complete medium to 10 µM and added to the wells, with the solution changed daily. Four replicates were seeded for each condition, and the plates were incubated at 37°C and 5% CO_2_. After 4 days, the media were discarded, wells were washed with 1x DPBS, fixed with 4% FA for 10 min at RT, and washed again. The cells were then permeabilized with PBS-Triton 0.1% (PBSTx) for 5 min at RT and blocked for 1 h at RT with PBDX-GS blocking solution (1% bovine serum albumin, 0.05% Triton, 1.5% goat serum, 1% DMSO in 1X DPBS). The cells were then stained with cleaved caspase-3 primary antibody overnight at 4 °C in a humid chamber. After washing steps with PBSTx, the cells were incubated overnight at 4 °C in a humid chamber with anti-rabbit 650 and conjugated anti-phalloidin 488, protected from light. Finally, cells were counterstained with DAPI (50 μg/ml) for 15min at RT, washed with ddH2O, and coverslips were mounted on glass microscope slides using Mowiol mounting medium (Sigma-Aldrich) and stored at 4 °C. Images were acquired using an Andor BC43 Confocal Microscope (20x objective, 512 x 512 px, FUSION Software, Oxford Instruments). Six to eight different random fields of view were acquired per coverslip (4 per condition). References for the antibodies used are provided in **Supplementary Table 1**.

Additionally, the sensitivity of all used cancer cell lines - CT26, MC38, HCT116, SW620, Hs578T - to increasing Tiotropium concentrations (1nM, 10nM, 100nM, 1uM, 10uM, 100uM, 1mM) was assessed using CellTiter Blue (Promega), according to the manufacturer’s instructions. All cell lines were seeded into a flat-bottom 96-well plate at 1.5 x 10^4^ cells per well and allowed to settle overnight. The cells were then treated with different concentrations of Tiotropium or DMSO for 24 and resorufin levels were measured using an Infinite M200 plate reader (TECAN). Cell viability (%) values were normalized to the ones obtained for the DMSO control.

### Animal care

All animals used in this study were handled and maintained according to the standard protocols of the European Animal Welfare Legislation, Directive 2010/63/EU (European Commission, 2016), and either the Champalimaud Foundation Fish Platform or the GIMM Rodent Platform. All protocols were approved by the necessary Portuguese institutional organizations - Órgão de Bem-Estar e Ética Animal (ORBEA) and Direção Geral de Alimentação e Veterinária (DGAV) - and by the Champalimaud Animal Ethical Committee or the GIMM Lisboa Ethical Committee.

### Zebrafish husbandry and lines

Zebrafish between 3 and 18 months of age were reared in 3.5 L tanks filled with E3 medium, at a density of 10 fish/L. The rearing temperature was 28°C and the animals were kept in a light/dark cycle of 14 h/10 h (lights on from 08:00 until 22:00). Zebrafish were fed three times per day, artemia in the mornings, and powder (Sparos 400-600, Techniplast) in the afternoons and evenings.

### Zebrafish transgenic lines

According to the purpose of each experiment, different genetically modified zebrafish lines, in the Tübingen background, were used in this study: Tg(*mpx:GFP*) i114Tg^134^, Tg(*mpeg1:mCherry*) ump2^135^, Tg(*csf1ra:GFP*) sh377Tg^136^, Tg(*mpeg1:mCherry-F; tnfa:GFP-F*) ump2Tg;ump5Tg^43^, Tg(*mpeg1:mCherry; nfkb:GFP*) ump2Tg;nc1Tg^137^, Tg(*fli:GFP*) y1Tg^138^, Tg(*chata:GFP*) [TgBAC*(chata:GAL4-VP16)* mpn202Tg^139^ x Tg(*UAS:GFP*)].

### Cell staining

Tumor cells were grown to 70% confluence in T-75 flasks, washed with 1x DPBS (Biowest), and enzymatically detached with TrypLE (Thermo Fisher Scientific). Cell suspensions were collected in 15 ml centrifuge tubes, centrifuged at 300 g for 4 min, and resuspended in 1x DPBS. If not transfected with fluorescent markers, cells were then stained in 1.5 ml microcentrifuge tubes using lipophilic dyes – Vybrant CM-DiI (4 µl/ml in DPBS 1x) or Deep Red Cell Tracker (1 µl/ml in DPBS 1x, 10 mM stock) (Life Technologies) – for 15 min at 37°C, while protected from light. Cells were then centrifuged at 300 g for 5 min and resuspended in complete medium. Viability was assessed by the Trypan Blue exclusion method, and cell number was determined by hemocytometer counting. Cells were resuspended in DPBS 1x to a final concentration of 0.25 × 10^6^ cells/μl, except for the Hs578T cell line, which was resuspended at a final concentration of 0.5 × 10^6^ cells/μl.

### Zebrafish macrophage ablation with clodronate liposomes

For the selective depletion of macrophages, liposome-encapsulated PBS (L-PBS) and liposome-encapsulated clodronate (L-Clodronate, 5 mg/ml) were purchased from Liposoma (CP-005-005). Immediately prior to injection into zebrafish, cells were resuspended in either PBS, L-PBS, or L-Clodronate.

### Zebrafish xenograft generation

On injection day, 2 dpf zebrafish embryos were separated from unhatched eggs, and Pronase (Roche) 1x was added to the E3 of the eggs to boost hatching. During injection, embryos were anesthetized by incubation in 1x Tricaine (Sigma-Aldrich) for 5 min, then transferred to an agarose plate and carefully aligned with a hairpin loop. Fluorescently labeled cancer cells were injected using a microinjection needle under a fluorescence scope (Zeiss Axio Zoom V16) with a milli-pulse pressure injector (Applied Scientific Instrumentation, MPPI-3). Approximately 500–1000 cells were injected into the perivitelline space (PVS) of each embryo, creating a tumor of a size similar to the embryo’s eye. After injection, embryos were kept in Tricaine for 10 min to allow wound closure, then transferred to fresh E3 medium and incubated at 34°C. At 1-day post-injection (dpi), zebrafish xenografts were screened for the presence or absence of a tumor under a fluorescence microscope. Xenografts with edema, cells in the yolk sac, and cellular debris were discarded, whereas successfully injected ones were grouped by tumor size, classified by comparison with the embryos’ eye size.

### Drug screening

For drug screening, we used an FDA-approved drug library (Screen-Well® FDA Approved Drug Library V2, version 1.2, BML-2843, Enzo Life Sciences). 556 compounds were tested (**Supplementary Data 1**) in “progressor” CRC and BC zebrafish xenografts: HCT116 and Hs578T cell lines, respectively. At 1 dpi, xenografts were divided into groups of at least 30 per condition, balanced for initial tumor size, and maintained in 6-well plates at 34 °C. Drugs or vehicle (DMSO) were diluted in E2 medium, at a standard concentration of 10 uM, and refreshed daily from 1 to 4-6 dpi. The effect of the tested compounds was determined phenotypically by quantifying tumor clearance as follows:

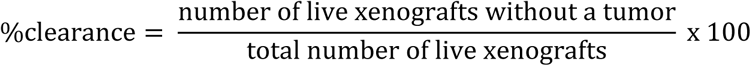

At the end of the assay, xenografts were euthanized with a Tricaine overdose, fixed with 4% Formaldehyde (FA, Thermo Scientific) overnight at 4 °C, and preserved at - 20 °C in 100% methanol. Compounds capable of increasing tumor clearance, when compared to controls, in at least one of the cell lines, were rescreened in a second round and considered hits if the effect was maintained. Toxic compounds at the standard concentration were excluded from the screen. Hit compounds were then purchased from new sources (**Supplementary Table 2**) for further experiments, to ensure reagent authenticity.

### Mechanism-of-action enrichment analysis

Compounds were annotated with a primary MOA class using the manufacturer’s descriptions and manually audited against the literature. Secondary classes were assigned when the activity was either independently responsible for a documented clinical or adverse effect, or a recognized mechanistic contributor to the approved pharmacological action. For example, the class Ia antiarrhythmic Disopyramide was assigned a primary class of voltage-gated Na⁺ channel blocker and a secondary class of muscarinic receptor antagonist, since its anticholinergic activity is independently responsible for its characteristic adverse effects. Conversely, the antihypertensive Methyldopa was assigned a secondary class of adrenergic receptor agonist, since its conversion to α-methylnorepinephrine and subsequent α2-adrenergic agonism is the mechanism underlying its approved action.

Compounds were counted in every class they carry, so class totals exceed the 556 screened compounds and classes are not independent. Classes containing fewer than three compounds were omitted.

Enrichment was assessed per class by two-sided Fisher exact test against the remainder of the library, with odds ratios also computed (**Supplementary Data 2**). P values are nominal, with no multiple-testing correction, given the small number of hits and the non-independence of classes. The analysis was repeated using primary MOA class alone as a sensitivity control.

For the neurotransmitter-system analysis, compounds were regrouped by the neurotransmitter system of their molecular target, irrespective of MOA class or clinical indication, and tested by the same procedure (**Supplementary Data 3**). All analyses were performed using Python.

### Zebrafish whole-mount immunofluorescence

Xenografts stored in 100% methanol were rehydrated through a decreasing percentage series (75% > 50% > 25% methanol in PBSTx). Next, xenografts were permeabilized with ice-cold acetone at −20°C for 7 min, then blocked in PBDX_GS for 1 h at RT. Staining with primary antibodies was performed overnight, followed by an additional overnight incubation with a secondary antibody and DAPI for nuclear counterstaining. Washing was performed between all steps. Stained xenografts were then mounted between two rectangular coverslips using Mowiol mounting medium (Sigma-Aldrich), enabling double-sided imaging. References and dilutions of the used antibodies are provided in **Supplementary Table 3**.

### Imaging and analysis of zebrafish xenografts

All images, both from fixed and live xenografts, were obtained using an Andor BC43 Confocal Microscope (20x objective, FUSION Software, Oxford Instruments), in Z-stacks of 3 (live) or 5 (fixed) μm intervals. For live imaging, tricaine-anesthetized xenografts were mounted in 0.8% low-melting agarose in a FluoroDish (WPI) filled with Tricaine 1x in E3, and placed inside the microscope’s incubator at 34°C. Generated images were analyzed using the Fiji (ImageJ, v1.54r) software^140^. Tumor area was quantified by measuring the area occupied by the fluorescently labeled tumor cells in all slices of the Z-stack of each xenograft. Macrophages and neutrophils were quantified in each slice using the cell counter plugin, and myeloid cell abundance was calculated as the ratio of the total number of each cell type to the tumor area of each xenograft. Macrophage engulfment events were defined as colocalization of fluorescent tumor cells within macrophages.

### Mouse xenograft generation and treatment

Eight- to twelve-week-old BALB/c mice were obtained from Charles River. NSG mice^141^ were a generous gift from João Barata Lab (Gulbenkian Institute for Molecular Medicine). Cultured CT26 cells at 70-80% confluency were detached with TrypLE Express (Thermo Fisher Scientific) for 3-5 minutes, centrifuged, and placed in PBS. On day 0, 5×10^5^ cells were injected subcutaneously into the abdomen of BALB/c or NSG mice, in 100 μl of a 1:1 mixture of Dulbecco’s modified Eagle medium and Matrigel (Corning). Once palpable, tumors were measured using a caliper, and volume was calculated as:

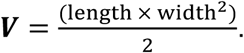

Once tumor volumes reached 100-200 mm^3^, Tiotropium (0.1 or 0.5 mg/kg) or DMSO was administered daily by intratumoral injection in 50 μl of PBS. For checkpoint blockade, mice were treated intravenously every three days with InVivoMAb anti-mouse PD-1 (clone RMP1-1, Bio X Cell) or mouse IgG2a (2A3) isotype control in PBS. Toxicity was monitored by daily weighing. Mice were euthanized when tumor volume reached 1 cm^3^. Efficacy and survival experiments were performed twice independently (**Figure 7**, **Supplementary Figure 8**).

### Toxicity assessment by histopathology

At study termination, representative samples of liver, kidney, lung, heart and spleen were collected from control (n = 4), anti-PD-1-treated (n = 4), Tiotropium-treated (n = 8; 0.1 and 0.5 mg/kg pooled) and anti-PD-1 plus Tiotropium-treated (n = 4) mice. Tissues were fixed in neutral-buffered formalin, routinely processed, embedded in paraffin, sectioned at 4 µm, and stained with hematoxylin and eosin. Sections were evaluated by a veterinary pathologist blinded to the experimental groups for alterations in tissue architecture and for inflammatory, degenerative, necrotic, vascular, and proliferative lesions, according to standardized diagnostic criteria for lesions in laboratory rodents^142–146^.

### Tumor dissociation and spectral flow cytometry

At day 12 post-inoculation, tumors were excised, minced, and digested using a mix of collagenase I, collagenase IV (Worthington) and DNase I (Roche) in a shaker for 20 min, 250 rpm. at 37 °C. After digestion, samples were filtered through a 100-μm cell strainer, erythrocytes were lysed using RBC Lysis Buffer (BioLegend), and cells were washed and counted.

Cells were then incubated with TruStain FcX™ PLUS (1:100 dilution, BioLegend), True-Stain Monocyte Blocker (1:50 dilution, BioLegend), and a viability dye (Zombie NIR, 1:2000 dilution, BioLegend) in PBS for 15 min at 4°C. Staining for extracellular markers - CD45, CD3, CD4, CD8, CD19, CD11c, F4/80, Ly6C, Ly6G, CD49b, MHC-II, SIRPα, PD-1 - was performed for 30 min at RT in PBS with 2% FBS and 1 mM EDTA. BD Horizon Brilliant Stain Buffer PLUS (Waters) was included in any mix that included at least two different BD Horizon Brilliant dyes, according to the manufacturer’s instructions. Cell suspensions were then fixed, permeabilized and stained for intracellular markers - CD206, FOXP3, CD163, iNOS - overnight at 4 °C using the eBioscience Foxp3 Transcription Factor Staining Buffer Set (ThermoFisher Scientific). Clones and dilutions of the antibodies are provided in **Supplementary Table 4.**

The samples were then analyzed in a FACSymphony A5 SE flow cytometer (BD Biosciences), equipped with BD FACSDiva software. Analysis was performed using FlowJo (v10.10.2), taking advantage of the t-SNE feature^147^ for unsupervised dimensionality reduction and the FastPhenoGraph plugin^148^ for unsupervised clustering and population identity. Group comparisons used unpaired two-tailed *t*-tests.

### Statistical Analysis

Unless stated otherwise, statistical analysis was performed using the GraphPad Prism software version 10.4.1. The ROUT method was used with a Q=1% to identify outliers, which were excluded from further analysis. Data were assessed for normality using the D’Agostino & Pearson and Shapiro-Wilk tests. In general, data sets with a Gaussian distribution were analyzed by parametric unpaired two-tailed t-test or ordinary one-way ANOVA with multiple comparisons, whereas the sets that did not pass the normality tests were analyzed by non-parametric unpaired Mann–Whitney U test. Most often, each experimental data set was compared with its respective control, and two-sided tests were used with a 95% confidence level. For mouse studies, tumor growth values and weight variation were analyzed by two-way ANOVA, and survival comparisons were performed using the log-rank Mantel–Cox test. Spectral flow cytometry data were compared between treatment arms using unpaired two-tailed t-tests. Statistical differences were considered significant whenever P-value (*P*)<0.05 and statistical output was represented by stars as follows: non-significant (*ns*)>0.05, \**P*<0.05; \*\**P*<0.01; \*\*\**P*<0.001; \*\*\*\**P*<0.0001. Violin plots show the median and interquartile range, while column plots present the data as the mean ± standard deviation (STD).

## Acknowledgements

We are grateful to all members of the Fior Lab for their support and critical discussion. To the Gonçalo Bernardes Lab for their collaboration and handling of mouse experiments. To the Champalimaud Foundation (CF) Fish Platform (Catarina Certal, Joana Monteiro and team) for excellent animal care and the CF Glass Wash and Media Preparation Platform (Maria Vito) for their technical support. To the Flow Cytometry Facility of Gulbenkian Institute for Molecular Medicine for their technical support, funded by FLxFlow-HDSpect-LISBOA2030-FEDER-01318900.

## Author contributions

Conceptualization by R.F, zebrafish experiments performed by A.B.M., I.O., C.R.A., B.C., L.L., B.S., R.F., and V.P.O., mouse experiments performed by C.L.A., R.C., and C.T., data analysis performed by A.B.M., I.O., J.X.C., and C.L.A., writing by A.B.M. with revisions by R.F.

## List of Abbreviations

AADC: aromatic L-amino acid decarboxylase
ACh: acetylcholine
ANS: autonomic nervous system
AKT: protein kinase B
BC: breast cancer
COMT: catechol-O-methyltransferase
COPD: chronic obstructive pulmonary disease
COX: cyclooxygenase
CRC: colorectal cancer
CSF1R: colony-stimulating factor 1 receptor
DMSO: dimethyl sulfoxide
DPBS: Dulbecco’s phosphate-buffered saline
dpf: days post-fertilization
dpi: days post-injection
DRD: dopamine receptor
EGFR: epidermal growth factor receptor
ERK: extracellular signal-regulated kinase
FA: formaldehyde
FBS: fetal bovine serum
FDA: United States Food and Drug Administration
GPCR: G protein-coupled receptor
ICB: immune checkpoint blockade
IL: interleukin
IT: intratumoral
IV: intravenous
JAK2: Janus kinase 2
LPS: lipopolysaccharide
MAPK: mitogen-activated protein kinase
MDSC: myeloid-derived suppressor cell
MHC: major histocompatibility complex
MOA: mechanism of action
MR: muscarinic receptor
MSS: microsatellite stable
NFκB: nuclear factor kappa B
nAChR: nicotinic acetylcholine receptor
NK: natural killer
NNCS: non-neuronal cholinergic system
NSAID: non-steroidal anti-inflammatory drug
NSG: NOD scid gamma
PBSTx: PBS Triton 0.1%
PD-1: programmed cell death protein 1
PD-L1: programmed cell death ligand 1
PD-L2: programmed cell death ligand 2
PDD: phenotypic drug discovery
PDE: phosphodiesterase
PVS: perivitelline space
RT: room temperature
SIRPα: signal regulatory protein alpha
STAT: signal transducer and activator of transcription
STING: stimulator of interferon genes
TAM: tumor-associated macrophage
TDD: target-based drug discovery
TME: tumor microenvironment
TNFα: tumor necrosis factor alpha

## Supplementary

**Supplementary Figure 1 -** Tumor clearance induced by the screen hit compounds in HCT116 and Hs578T xenografts.

**Supplementary Figure 2 -** Re-screening of hit compounds shows sustained tumor clearance in multiple progressor models.

**Supplementary Figure 3 -** CT26 and HCT116 cell lines show no i*n vitro* cytotoxicity to Tiotropium or Amrinone treatment.

**Supplementary Figure 4 -** Tiotropium does not change the viability of multiple cancer cell lines *in vitro*.

**Supplementary Figure 5 -** Extended immune characterization of Tiotropium and Amrinone across multiple xenograft models.

**Supplementary Figure 6 -** Drug treatments have no significant effects on macrophage or neutrophil distribution in non-tumor-bearing zebrafish larvae.

**Supplementary Figure 7 -** Dose-response and toxicity assessment of Tiotropium and Amrinone as monotherapies in a CT26 mouse tumor model.

**Supplementary Figure 8 -** Tiotropium synergizes with αPD-1 checkpoint blockade in an independent CT26 replicate experiment.

**Supplementary Figure 9 -** Representative histopathology of selected organs following anti-PD-1, Tiotropium, or combination treatment.

**Supplementary Figure 10 -** Mouse body weight variation in mouse experiments.

**Supplementary Figure 11 -** Immunophenotyping of CT26 tumors by spectral flow cytometry.

**Supplementary Table 1 -** List of antibodies used for immunofluorescence of *in vitro* cultures.

**Supplementary Table 2 -** List of screen hit compounds purchased from independent sources.

**Supplementary Table 3 -** List of antibodies used for immunofluorescence of zebrafish xenografts.

**Supplementary Table 4 -** List of antibodies used for spectral flow cytometry analysis of mouse xenograft tumors.

