## Supplementary Figures and Tables for "Fishing for new immunotherapy compounds to boost innate tumor rejection"

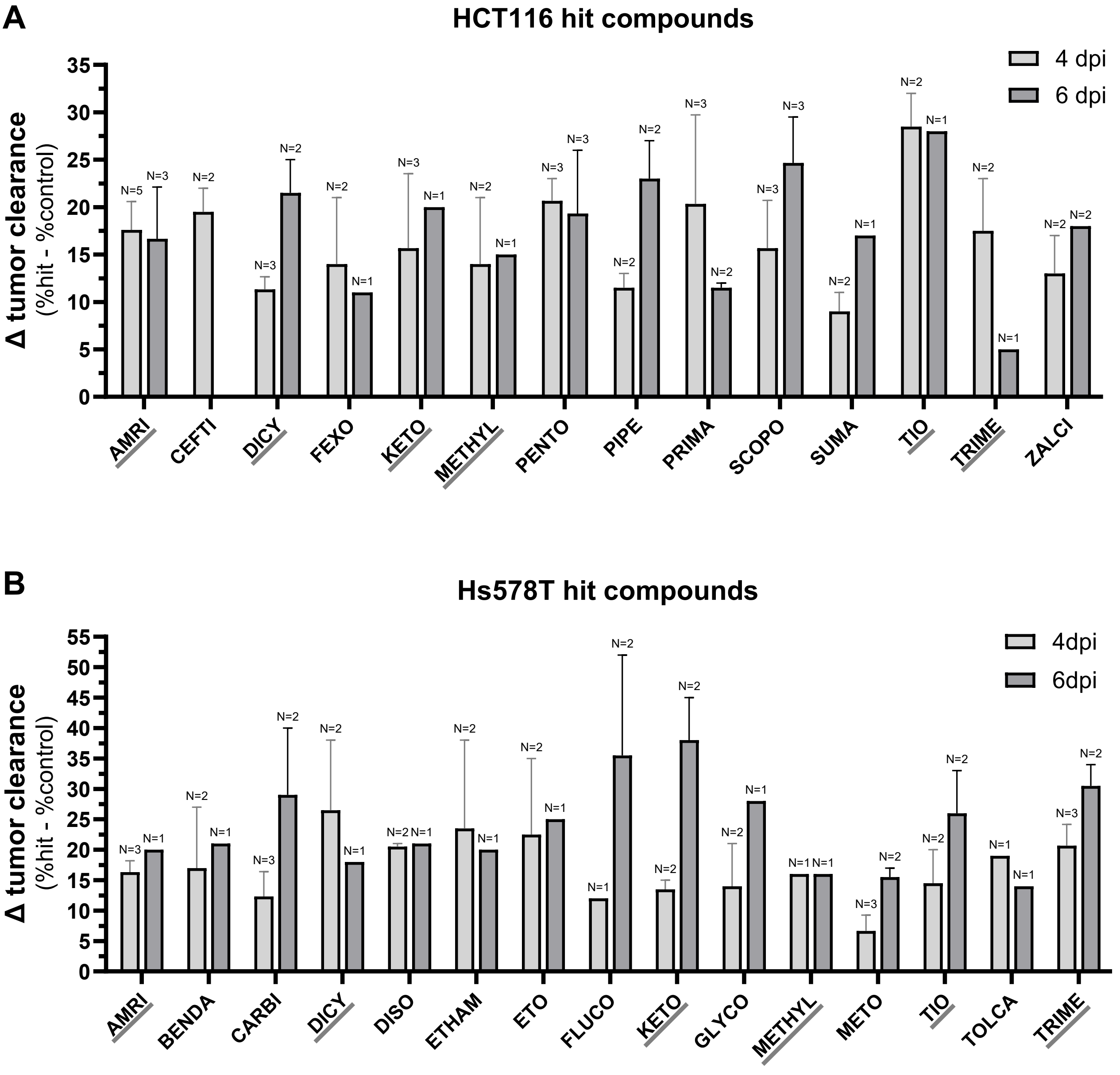


**Supplementary Figure 1 – Tumor clearance induced by the screen hit compounds in HCT116 and Hs578T xenografts.**

Induced tumor clearance of screen hit compounds in **(A)** HCT116 and **(B)** Hs578T zebrafish xenograft models at 4 and 6 dpi, quantified as the difference in tumor clearance between treated and control conditions. Tumor clearance is the percentage of fish with no tumor at the end of the assay. Compounds confirmed as hits in both models are underlined in grey. Graph shows mean ± STD and N indicates the number of independent experiments.

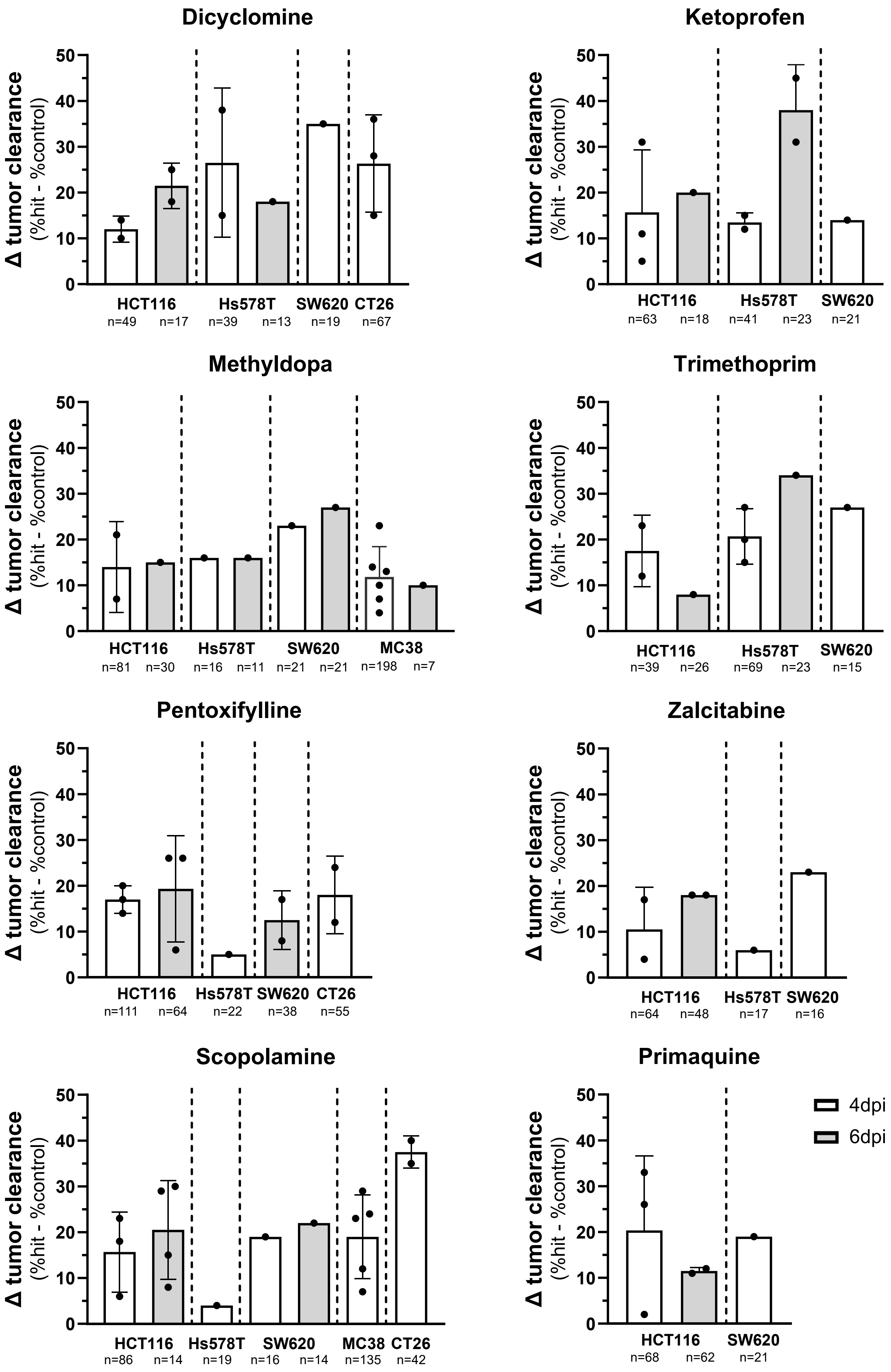
Dicyclomine
Δ tumor clearance
(%hit - %control)
50
40
30
20
10
0
HCT116
n=49 n=17
Hs578T
n=39 n=13
SW620
n=19
CT26
n=67
Ketoprofen
HCT116
n=63 n=18
Hs578T
n=41 n=23
SW620
n=21
Methyldopa
HCT116
n=81 n=30
Hs578T
n=16 n=11
SW620
n=21 n=21
MC38
n=198 n=7
Trimethoprim
HCT116
n=39 n=26
Hs578T
n=69 n=23
SW620
n=15
Pentoxifylline
HCT116
n=111 n=64
Hs578T
n=22
SW620
n=38
CT26
n=55
Zalcitabine
HCT116
n=64 n=48
Hs578T
n=17
SW620
n=16
Scopolamine
HCT116
n=86 n=14
Hs578T
n=19
SW620
n=16 n=14
MC38
n=135
CT26
n=42
Primaquine
HCT116
n=68 n=62
SW620
n=21
4dpi
6dpi

**Supplementary Figure 2 – Re-screening of hit compounds shows sustained tumor clearance in multiple progressor models.**

Induced tumor clearance of hit compounds in multiple progressor zebrafish xenograft models at 4 and 6 dpi, quantified as the difference in tumor clearance between treated and control conditions. Tumor clearance is the percentage of fish with no tumor at the end of the assay. Graphs show mean ± STD, each dot represents one independent experiment, and n shows the total number of fish per condition.

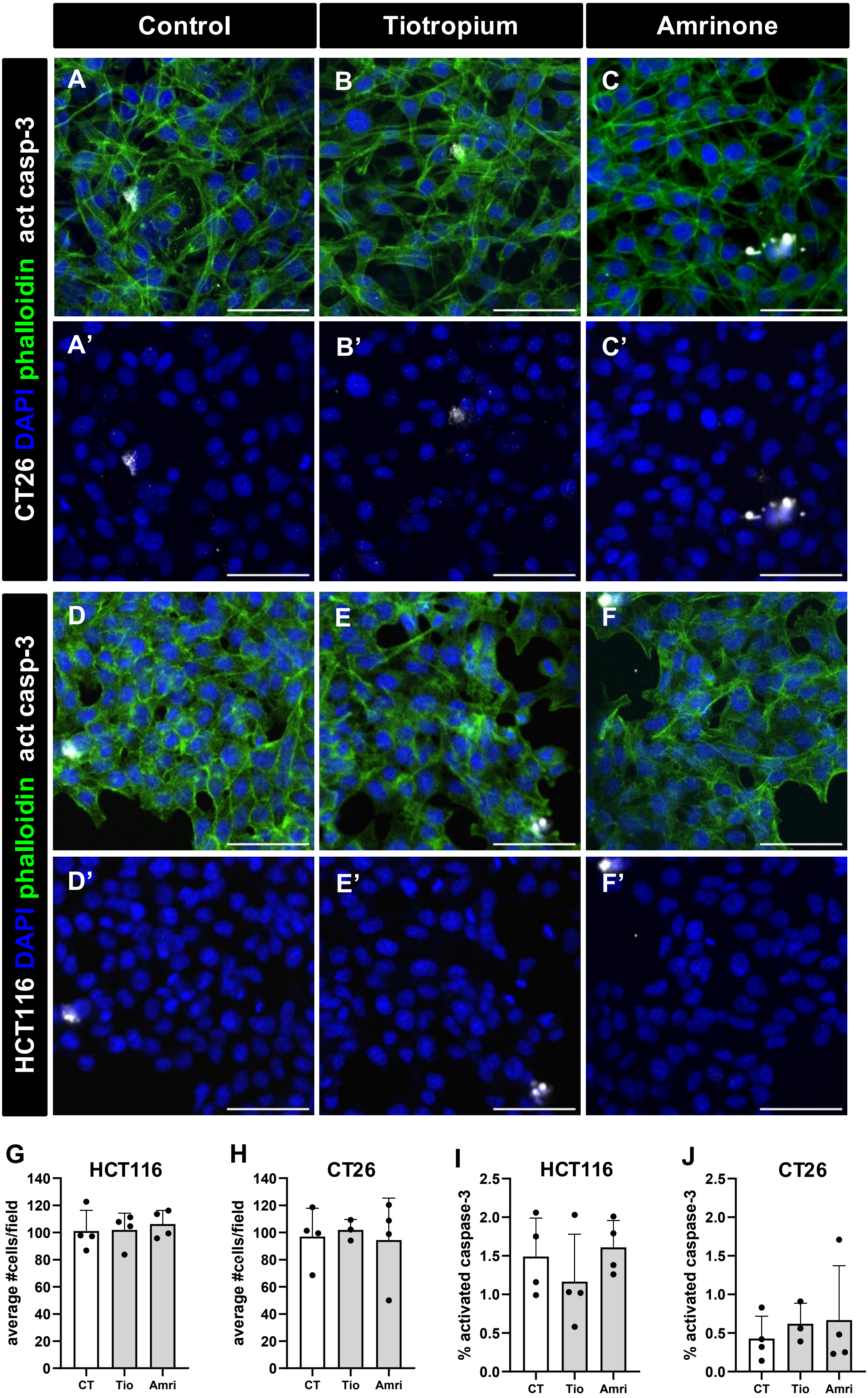
Control
Tiotropium
Amrinone
CT26 DAPI phalloidin act casp-3
HCT116 DAPI phalloidin act casp-3
A
B
C
A'
B'
C'
D
E
F
D'
E'
F'
G
HCT116
average #cells/field
CT
Tio
Amri
H
CT26
average #cells/field
CT
Tio
Amri
I
HCT116
% activated caspase-3
CT
Tio
Amri
J
CT26
% activated caspase-3
CT
Tio
Amri

**Supplementary Figure 3 - CT26 and HCT116 cell lines show no i*n vitro* cytotoxicity to Tiotropium or Amrinone treatment.**

Representative confocal images of CT26 (**A**-**C**) and HCT116 (**D**-**F**) cells at 4 days post-seeding, after immunocytochemistry. Phalloidin in green, activated caspase 3 in white, nuclei (DAPI) in blue. To determine possible direct cytotoxicity of the Tiotropium and Amrinone (10uM), average number of cells per field of view (**G**-**H**) and percentage of apoptotic cells (positive for activate caspase-3, **I**-**H**) were quantified and compared to control (DMSO). Graphs show mean ± STD. Each dot represents one coverslip (3-4), which represents the average of 6-8 different fields of view. All data was analyzed using the Kruskal-Wallis test (Control vs Tiotropium *ns*, Control vs Amrinone *ns*, in all graphs). Scale bars represent 50 µm.

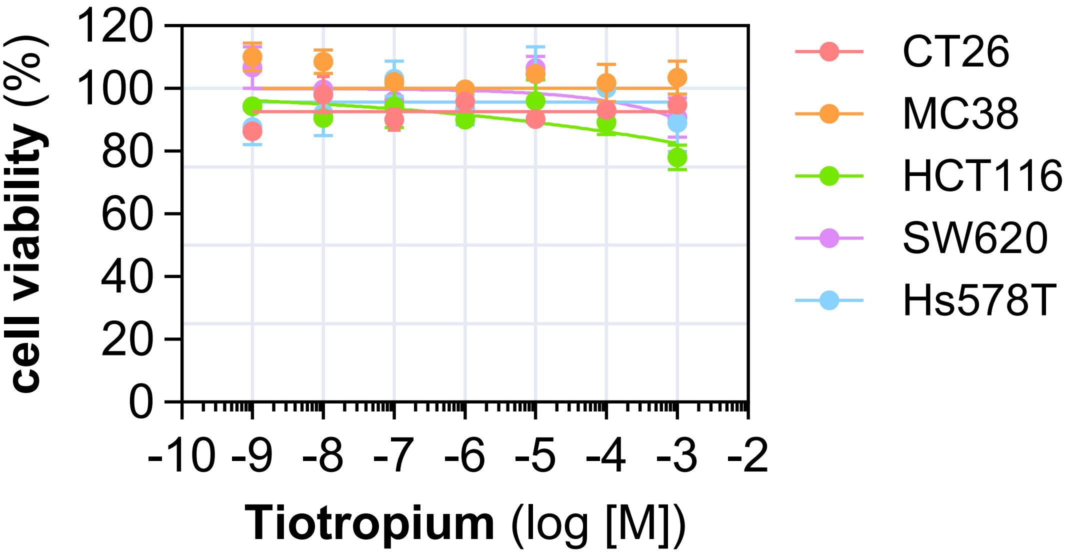


**Supplementary Figure 4 – Tiotropium does not change the viability of multiple cancer cell lines *in vitro.***

Multiple cancer cell lines – CT26, MC38, HCT116, SW620, Hs578T – were exposed to increasing concentrations of Tiotropium, and viability was assessed by Cell Titer Blue assay. Data is shown as mean ± STD.

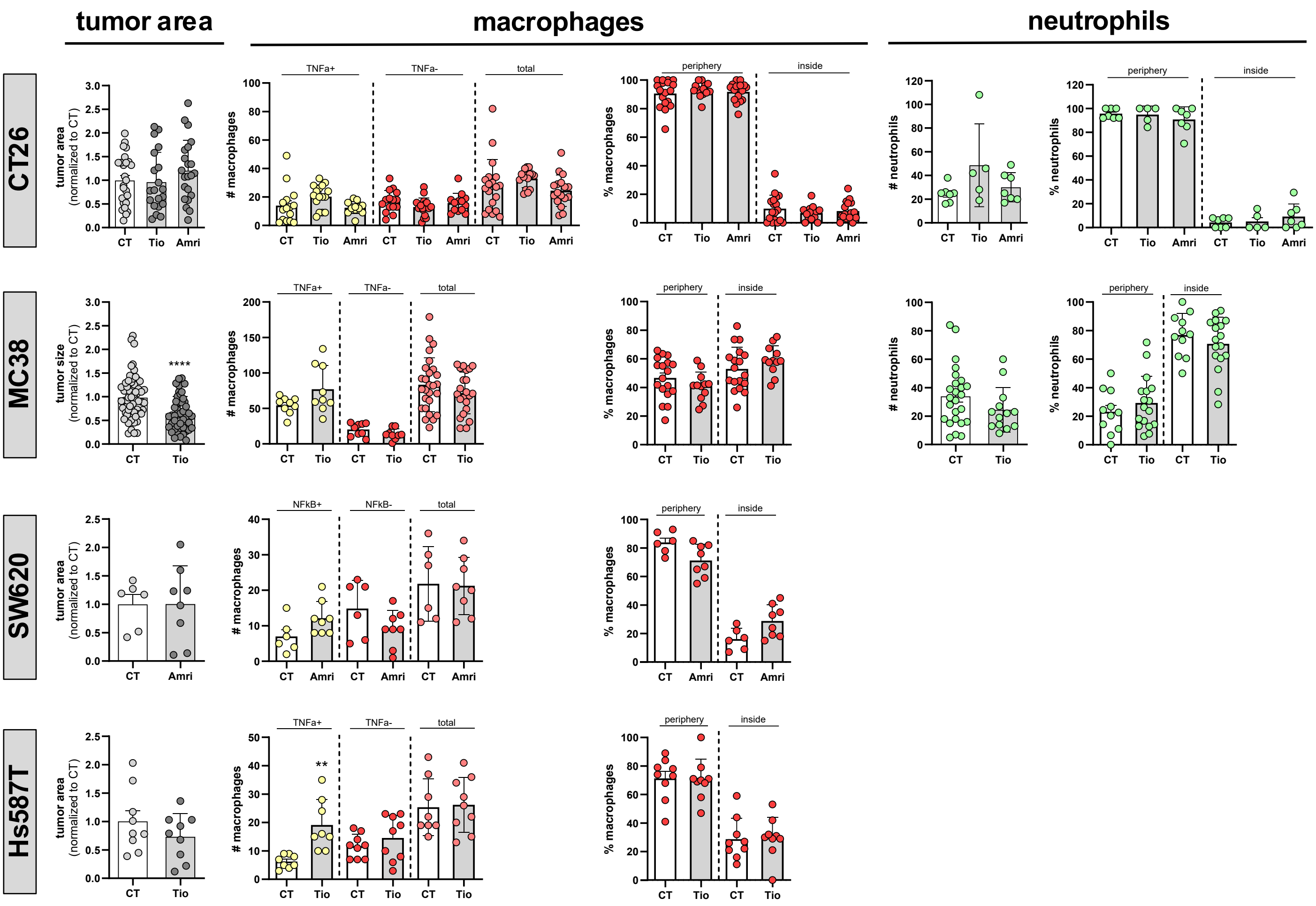


**Supplementary Figure 5 – Extended immune characterization of Tiotropium and Amrinone across multiple xenograft models.**

Quantification of immune characterization features in control (CT), Tiotropium-treated (Tio), and Amrinone-treated (Amri) zebrafish xenografts of CT26, MC38, SW620, and Hs578T progressor cell lines, at 4 dpi. Within each row, graphs show from left to right, when available: tumor area normalized to the control condition; absolute number of macrophages per xenograft (TNFα/NFκB-positive, TNFα/NFκB-negative, and total); macrophage distribution, as the percentage of cells localized at the periphery versus inside the tumor; absolute number of neutrophils; and neutrophil distribution. All graphs show mean ± S.D. and each dot represents one zebrafish xenograft. According to data normality, comparisons between control and treated conditions were performed using Welch's t-test or Mann-Whitney U test (**P ≤ 0.01, ****P ≤ 0.0001).

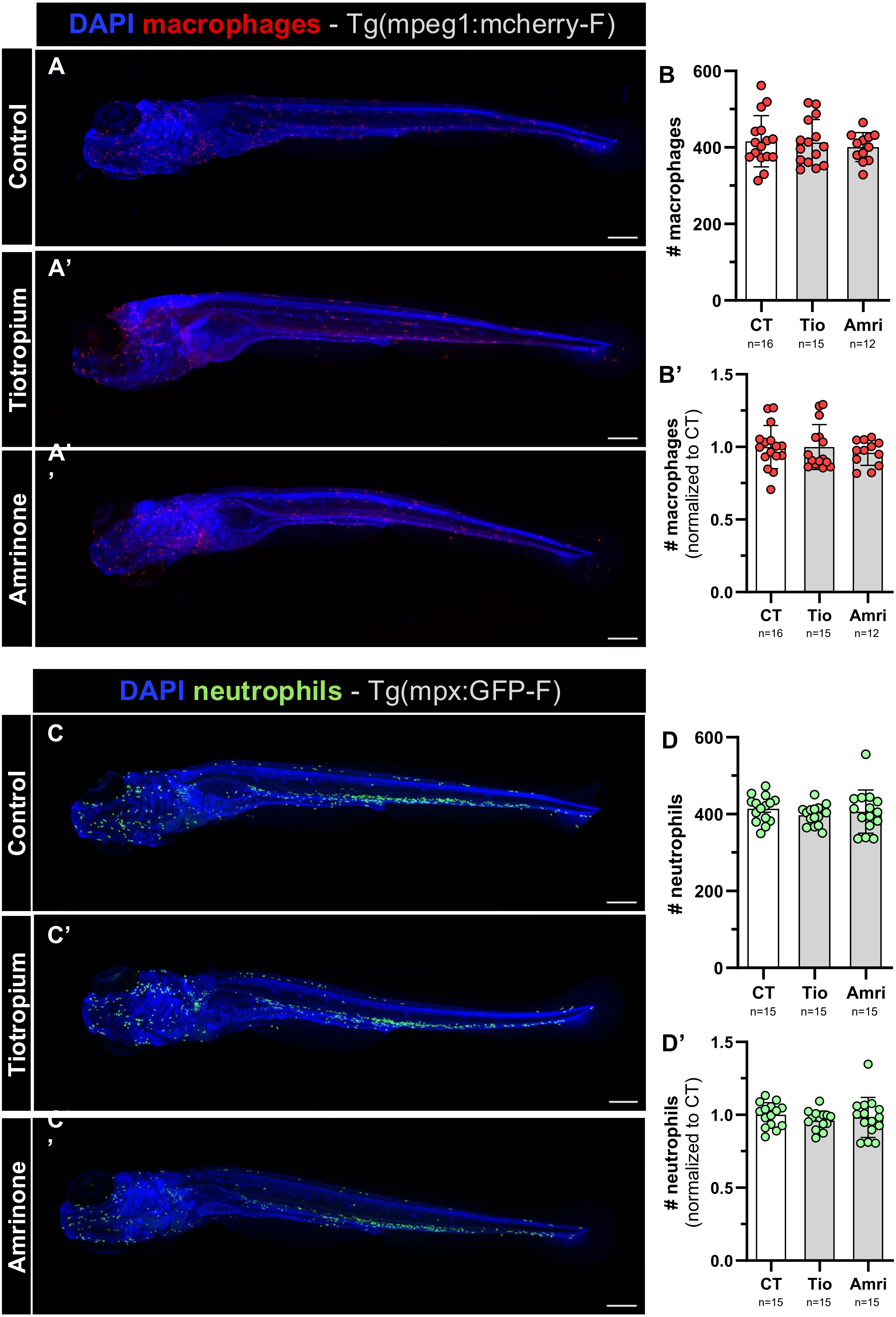

DAPI macrophages - Tg(mpeg1:mcherry-F)
A
Control
A'
Tiotropium
A''
Amrinone
B
### macrophages
600
400
200
0
CT
n=16
Tio
n=15
Amri
n=12
B'
### macrophages (normalized to CT)
1.5
1.0
0.5
0.0
CT
n=16
Tio
n=15
Amri
n=12
DAPI neutrophils - Tg(mpx:GFP-F)
C
Control
C'
Tiotropium
C''
Amrinone
D
### neutrophils
600
400
200
0
CT
n=15
Tio
n=15
Amri
n=15
D'
### neutrophils (normalized to CT)
1.5
1.0
0.5
0.0
CT
n=15
Tio
n=15
Amri
n=15

**Supplementary Figure 6 - Drug treatments have no significant effects on macrophage or neutrophil distribution in non-tumor-bearing zebrafish larvae.**

Representative full-body confocal images of macrophages (**A**, red) and neutrophils (**C**, green) of control and Tiotropium or Amrinone-treated larvae at 6 days post-fertilization. Quantification of total body macrophages (**B**, absolute number; **B'**, normalized to CT*)* and neutrophils (**D**, absolute number; **D'**, normalized to CT). Bars indicate the results as mean ± STD, and each dot represents one embryo pooled from two independent experiments. All data was analyzed using ANOVA or the Kruskal-Wallis test, according to normality (Control vs Tiotropium *ns*, Control vs Amrinone *ns*, in all graphs). All images are oriented with the left side anterior, the right side posterior, the dorsal side up, and the ventral side down. Scale bars represent 200µm.

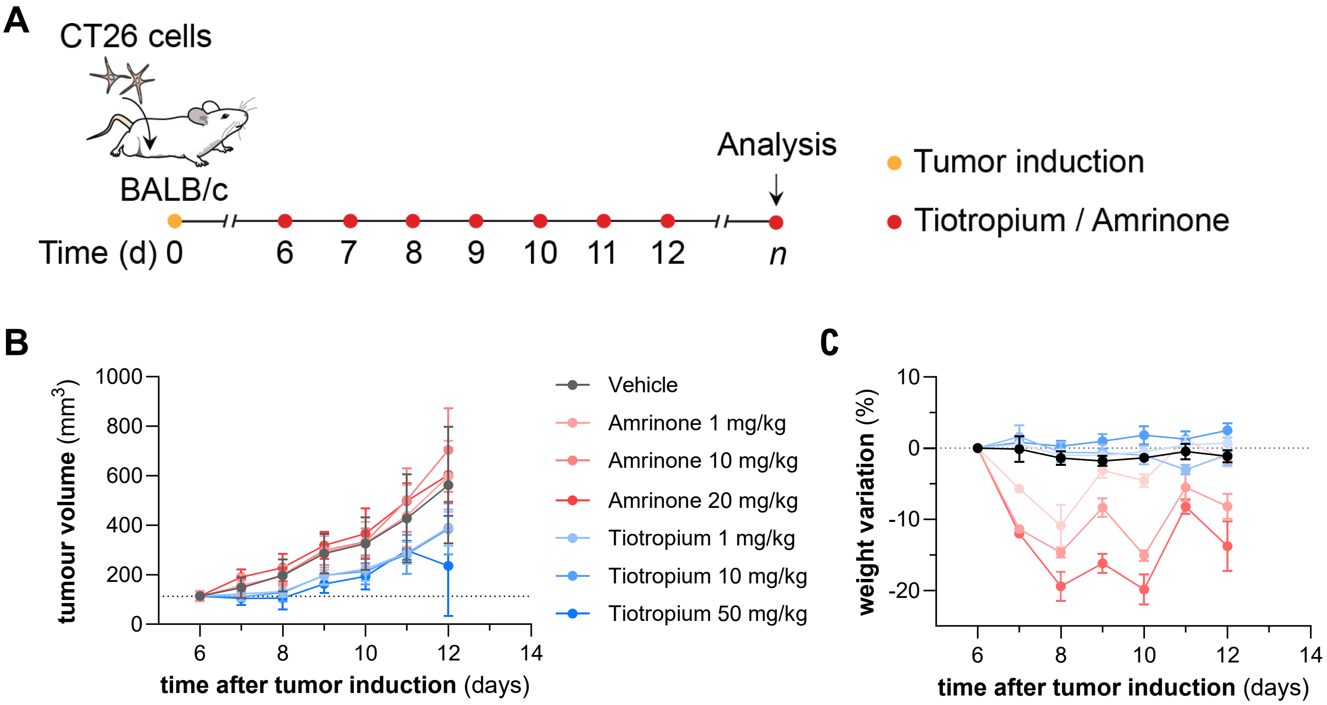


**Supplementary Figure 7 – Dose-response and toxicity assessment of Tiotropium and Amrinone as monotherapies in a CT26 mouse tumor model.**

**(A)** Experimental design for testing increasing dosages of Tiotropium and Amrinone. CT26 cells were inoculated subcutaneously into the abdomens of BALB/c mice. Treatments began once tumor volume reached 100-200 mm³. Amrinone: 1, 10, or 20 mg/kg (IT), Tiotropium: 1, 10, or 50 mg/kg (IT). n = 5 for each treatment group. IV, intravenous; IT, intratumoral. (**B**) Tumour volume curves. Treatment groups were compared to vehicle at day 7 of treatment (day 12 after tumor induction) using the Kruskal-Wallis test with Dunnett's T3 multiple comparisons test. Data are presented as mean ± STD. (**C**) Mouse body weight curves. Data are presented as mean ± SEM.

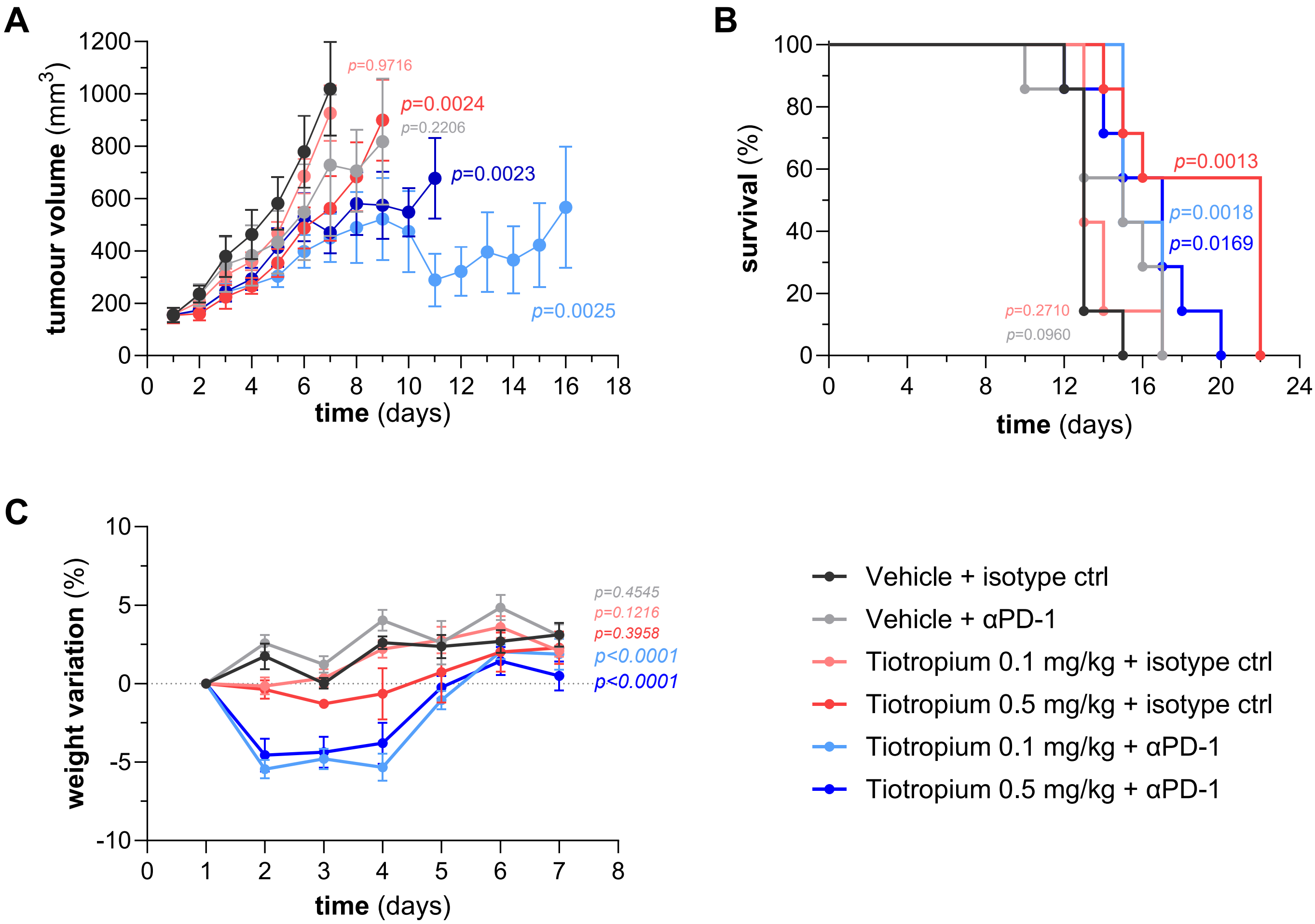


**Supplementary Figure 8 – Tiotropium synergizes with αPD-1 checkpoint blockade in an independent CT26 replicate experiment.**

CT26 tumor-bearing BALB/c mice were treated with intratumoral Tiotropium (0.1 or 0.5 mg/kg) and intravenous αPD-1 (200 µg/mouse) alone or in combination, following the same experimental design as Figure 5 and serving as an independent replicate experiment. n = 7 per treatment group. **(A)** Tumor volume curves. Treatment groups were compared to vehicle + isotype control using Brown-Forsythe and Welch one-way ANOVA with Dunnett's T3 multiple comparisons test. Data are presented as mean ± S.D. **(B)** Survival curves were compared using log-rank Mantel-Cox tests. **(C)** Mouse body weight variation, presented as mean ± S.E.M. The *p*-values shown in the plots correspond to each treatment group (color-coded) compared to vehicle + isotype control.

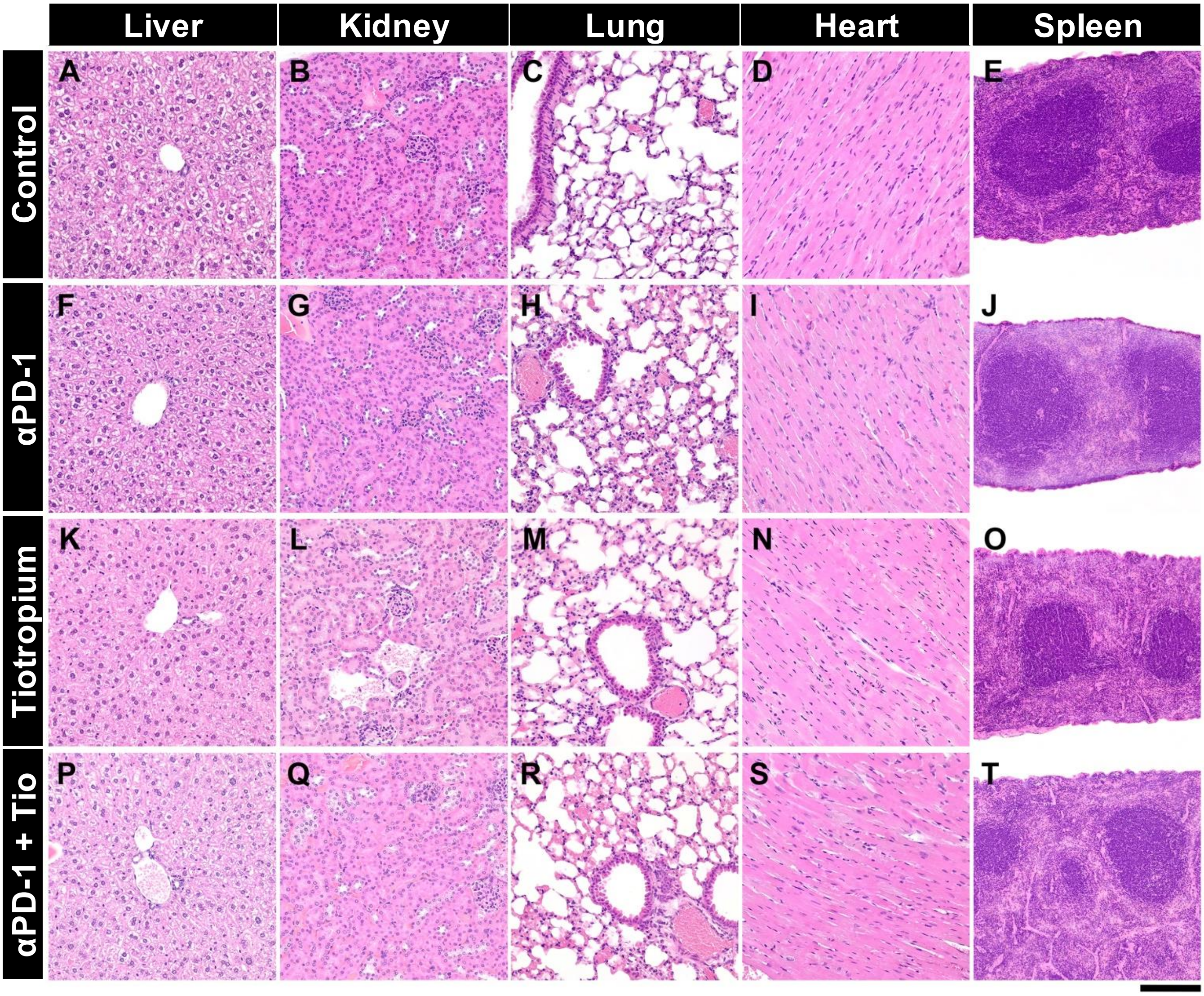


**Supplementary Figure 9 - Representative histopathology of selected organs following anti-PD-1, Tiotropium, or combination treatment.**

Representative H&E-stained sections of liver, kidney, lung, heart, and spleen from control (**A-E**; *n* = 4), αPD-1-treated (**F-J**; *n* = 4), Tiotropium-treated (**K-O**; *n* = 8), and combination-treated (**P-T**; *n* = 4) mice. Across all groups, the examined organs retain normal histological architecture, without treatment-related inflammatory, degenerative, necrotic, vascular, or proliferative lesions. H&E - Hematoxylin and Eosin. Original magnification: 20x for liver, kidney, lung, and heart (scale bar = 100 μm); 10x for spleen (scale bar = 200 μm).

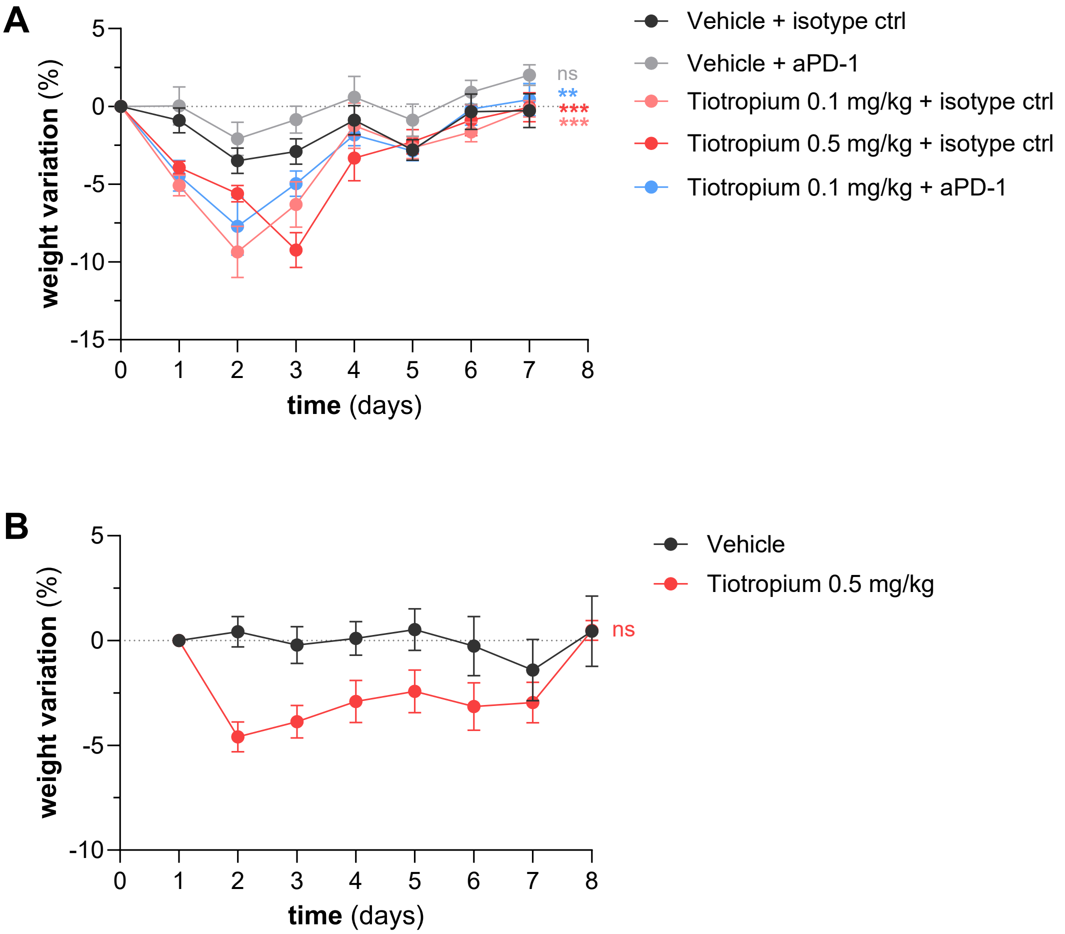


**Supplementary Figure 10 – Mouse body weight variation in mouse experiments.**

**(A)** Mouse body weight variation corresponding to the Tiotropium and αPD-1 combination experiment (Figure 5A–C), presented as mean ± S.E.M. Body weight variation was compared between treatment groups and vehicle control using two-way ANOVA: vehicle vs. Tiotropium 0.1 mg/kg $p<0.0001$, vehicle vs. Tiotropium 0.5 mg/kg $p<0.0001$, vehicle vs. αPD-1 $p=0.5965$, vehicle vs. combination $p=0.0038$. **(B)** Mouse body weight variation corresponding to the NSG experiment (Figure 5D–F), presented as mean ± S.E.M. No significant differences were observed.

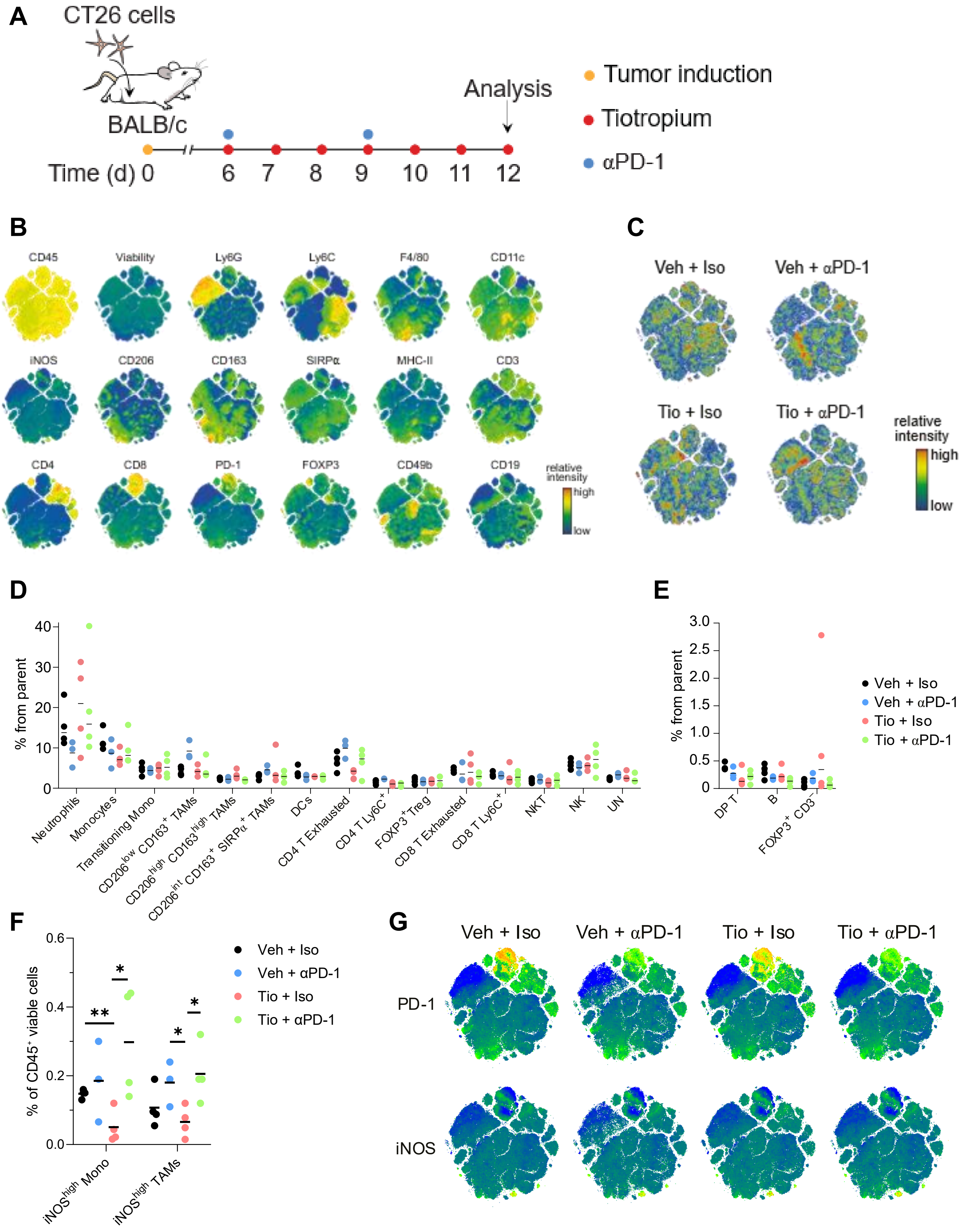

A
CT26 cells
BALB/c
Analysis
Time (d) 0 6 7 8 9 10 11 12
Tumor induction
Tiotropium
αPD-1
B
CD45
Viability
Ly6G
Ly6C
F4/80
CD11c
iNOS
CD206
CD163
SIRPα
MHC-II
CD3
CD4
CD8
PD-1
FOXP3
CD49b
CD19
relative intensity
high
low
C
Veh + Iso
Veh + αPD-1
Tio + Iso
Tio + αPD-1
relative intensity
high
low
D
% from parent
Neutrophils
Monocytes
Transitioning Mono
CD206low CD163+ TAMs
CD206high CD163high TAMs
CD206int CD163+ SIRPα+ TAMs
DCs
CD4 T Exhausted
CD4 T Ly6C+
FOXP3+ Treg
CD8 T Exhausted
CD8 T Ly6C+
NKT
NK
UN
E
% from parent
DP T
B
FOXP3+ CD3−
Veh + Iso
Veh + αPD-1
Tio + Iso
Tio + αPD-1
F
% of CD45+ viable cells
iNOShigh Mono
iNOShigh TAMs
Veh + Iso
Veh + αPD-1
Tio + Iso
Tio + αPD-1
G
Veh + Iso
Veh + αPD-1
Tio + Iso
Tio + αPD-1
PD-1
iNOS

**Supplementary Figure 10 - Immunophenotyping of CT26 tumors by spectral flow cytometry.**

**(A)** Experimental design for immunophenotyping of Tiotropium- and ICB-treated tumors by spectral flow cytometry. CT26 cells were inoculated subcutaneously into the abdomens of BALB/c mice. Treatments began once tumor volume reached 100-200 $mm^3$. αPD-1: 200ug/mouse (IV); Tiotropium: 0.1 mg/kg (IT). IV, intravenous; IT, intratumoral. **(B)** t-SNE heat plots of the 18 markers included in the analysis. **(C)** t-SNE heat plots showing the distribution and relative frequency of events per treatment condition. **(D-E)** Frequencies of immune clusters across treatment conditions. **(F)** Frequencies of $iNOS^{high}$ monocytes and TAMs. **(G)** t-SNE heat plots of PD-1 and iNOS expression per treatment condition. Horizontal lines in graphs indicate group means and each dot represents one animal. Data were analyzed using unpaired two-tailed *t*-tests (ns ≥ 0.05, *P ≤ 0.05, **P ≤ 0.01). TAMs: tumor-associated macrophages; DP: double-positive (CD4+CD8+) T cells; DCs: dendritic cells; Treg: regulatory T cells; NKT: natural killer T cells; NK: natural killer cells; UN: unassigned.

**Supplementary Table 1 -** List of antibodies used for immunofluorescence of *in vitro* cultures.

| Antibody | Species | Dilution | Brand | Reference |
|---|---|---|---|---|
| Anti-Cleaved Caspase-3 (Asp175) | Rabbit | 1:200 | Cell Signaling Technology | 9661 |
| Anti-Phalloidin AlexaFluor™ 488 | - | 1:500 | Thermo Fisher Scientific | A12379 |
| Anti-Rabbit IgG Dylight™ 650 | Goat | 1:500 | Thermo Fisher Scientific | 84546 |
| DAPI | - | 0.00015% in ddH20 | Sigma-Aldrich | 10236276001 |

**Supplementary Table 2 -** List of screen hit compounds purchased from independent sources.

| Compound name | Brand | Reference |
|---|---|---|
| Amrinone | MedChemExpress | HY-B1294 |
| | Merck | 1034308 |
| Carbidopa | APExBIO | B1904 |
| Ceftizoxime sodium | MedChemExpress | HY-B1596A |
| Dicyclomine hydrochloride | MedChemExpress | HY-B1339 |
| Disopyramide | MedChemExpress | HY-12533 |
| Ethambutol dihydrochloride | MedChemExpress | HY-B0535A |
| Etodolac | MedChemExpress | HY-76251 |
| Fexofenadine·HCl | MedChemExpress | HY-B0801 |
| Fluconazole | APExBIO | B2094 |
| Glycopyrrolate Iodide | MedChemExpress | HY-17465 |
| Ketoprofen | APExBIO | B1446 |
| L-(-)-α-Methyldopa (hydrate) | APExBIO | B1064 |
| Metoprolol Tartrate | APExBIO | B1339 |
| Pentoxifylline | MedChemExpress | HY-B0715 |
| Piperacillin sodium | MedChemExpress | HY-B1286 |
| Primaquine Diphosphate | APExBIO | B1820 |
| Scopolamine·HBr | APExBIO | N2570 |
| Sumatriptan succinate | MedChemExpress | HY-B0121 |
| Tiotropium Bromide | APExBIO | A3874 |
| | Supelco | PHR1890 |
| Tolcapone | APExBIO | A4383 |
| Trimethoprim | MedChemExpress | HY-B0510 |
| Zalcitabine | MedChemExpress | HY-17392 |

**Supplementary Table 3 -** List of antibodies used for immunofluorescence of zebrafish xenografts.

| Antibody | Species | Dilution | Brand | Reference |
|---|---|---|---|---|
| Anti-Cleaved Caspase-3 (Asp175) | Rabbit | 1:100 | Cell Signaling | 9661 |
| Anti-mCherry | Rabbit | 1:100 | Abcam | ab167453 |
| Anti-GFP | Mouse | 1:100 | Roche | ab1218 |
| Anti-Rabbit IgG AlexaFluor™ 488 | Goat | 1:400 | Thermo Fisher Scientific | A11034 |
| Anti-Rabbit IgG DyLight™ 594 | Goat | 1:400 | Thermo Fisher Scientific | 35560 |
| Anti-Rabbit IgG Dylight™ 650 | Goat | 1:400 | Thermo Fisher Scientific | 84546 |
| Anti-Mouse IgG Dylight™ 488 | Goat | 1:400 | Thermo Fisher Scientific | 35502 |
| DAPI | - | 50 µg/ml | Sigma-Aldrich | 10236276001 |

**Supplementary Table 4 -** List of antibodies used for spectral flow cytometry analysis of mouse xenograft tumors.

| Marker | Clones | Fluorochrome | Dilution |
|---|---|---|---|
| Zombie (viability) | - | NIR | 1:2000 |
| CD45 | 30-F11 | BV510 | 1:400 |
| CD3 | 17A2 | BV711 | 1:100 |
| CD8 | 53-6.7 | BV650 | 1:200 |
| CD4 | RM4-5 | BV786 | 1:800 |
| CD19 | 1D3 | BUV563 | 1:50 |
| F4/80 | BM8 | PECy7 | 1:800 |
| CD11c | N418 | BUV805 | 1:200 |
| Ly6C | HK1.4 | FITC | 1:400 |
| Ly6G | 1A8 | BV605 | 1:400 |
| CD49b | HMα2 | RB744 | 1:200 |
| MHC-II | M5/114.15.2 | PerCP | 1:400 |
| CD206 | C068C2 | PE-Fire700 | 1:500 |
| SIRPa | P84 | BUV737 | 1:500 |
| PD-1 | RMP1-30 | PE-CF594 | 1:100 |
| Foxp3 | FJK-16s | eFluor 450 | 1:50 |
| CD163 | S15049I | BV421 | 1:200 |
| iNOS | CXNFT | APC | 1:800 |
