## Supplementary Data for "Fishing for new immunotherapy compounds to boost innate tumor rejection"

**Supplementary Data 1 - All screened compounds.** Complete list of the 556 FDA-approved compounds screened in HCT116 and Hs578T zebrafish xenografts, with mechanisms of action (MOA), MOA class, clinical indication, and year of FDA approval.

| # | Compound | HCT116 | Hs578T | Mechanisms of action | Primary MOA class | Secondary MOA class(es) | Indication | FDA approval year |
|---|---|---|---|---|---|---|---|---|
| 1 | (±) Isoproterenol·HCl | No hit | No hit | β-adrenoceptor agonist | Adrenergic receptor agonist | | Bronchodilator | 1956 |
| 2 | (±)-Atenolol | No hit | No hit | β1-adrenoceptor antagonist (selective) | Adrenergic receptor antagonist | | Antihypertensive | 1981 |
| 3 | (S)-Timolol Maleate | No hit | No hit | β-adrenoceptor antagonist (non-selective) | Adrenergic receptor antagonist | | Antihypertensive | 1978 |
| 4 | 4-Aminosalicylic Acid | No hit | No hit | Bacterial dihydropteroate synthetase (DHPS) inhibitor | Folate pathway inhibitor | | Antitubercular | 1994 |
| 5 | Abacavir Sulfate | No hit | No hit | Nucleoside Reverse Transcriptase Inhibitor (NRTI) | Reverse transcriptase inhibitor | | Antiviral | 1998 |
| 6 | Acamprosate calcium | No hit | Not confirmed in 2nd round | N-methyl-D-aspartate (NMDA) receptor antagonist; GABA agonist | NMDA receptor antagonist | GABAergic modulator | Alcohol Deterrent | 2004 |
| 7 | Acarbose | No hit | No hit | α-glucosidase inhibitor | α-glucosidase inhibitor | | Antidiabetic | 1995 |
| 8 | Acebutolol·HCl | No hit | No hit | β1-adrenoceptor antagonist (selective) | Adrenergic receptor antagonist | | Antihypertensive | 1984 |
| 9 | Acetaminophen | No hit | No hit | COX inhibitor (NSAID) | COX inhibitor (NSAID) | | Analgesic | 1950 |
| 10 | Acetazolamide | No hit | No hit | Carbonic anhydrase inhibitor | Diuretic | | Diuretic | 1953 |

| | | | | | | | | |
|---|---|---|---|---|---|---|---|---|
| 11 | Acetohexamide | No hit | No hit | ATP-sensitive $K^+$ channel blocker (insulin secretagogue) | ATP-sensitive $K^+$ channel modulator | | Antidiabetic | 1964 |
| 12 | Acetohydroxamic Acid | No hit | No hit | Urease inhibitor | Other antimicrobial | | Antiurolithic | 1983 |
| 13 | Acetylcholine Chloride | No hit | No hit | Muscarinic receptor agonist; Nicotinic receptor agonist | Muscarinic receptor agonist | | Cholinergic agonist | 1973 |
| 14 | Acetylcysteine | No hit | No hit | Mucolytic (disulfide reduction); glutathione precursor | Mucolytic | | Expectorant | 1963 |
| 15 | Acitretin | Toxic | Not screened | Retinoid receptor agonist | Retinoid receptor agonist | | Dermatologic agent | 1996 |
| 16 | Acrivastine | No hit | No hit | H1 receptor antagonist (antihistamine) | H1 receptor antagonist | | Antihistamine | 1994 |
| 17 | Acyclovir (Acycloguanosine) Zovirax | No hit | No hit | Viral DNA polymerase inhibitor | Viral DNA polymerase inhibitor | | Antiviral | 1982 |
| 18 | Adapalene | No hit | Toxic | Retinoid receptor agonist | Retinoid receptor agonist | | Dermatologic agent | 1996 |
| 19 | Adefovir Dipivoxil | No hit | No hit | Nucleotide reverse transcriptase inhibitor (NtRTI) | Reverse transcriptase inhibitor | | Antiviral | 2002 |
| 20 | Adenosine | No hit | No hit | Adenosine receptor agonist | Adenosine receptor modulator | | Antiarrhythmic | 1989 |
| 21 | Albendazole | Toxic | Toxic | Helminth tubulin inhibitor | Antiparasitic | | Anthelmintic | 1996 |
| 22 | Alendronate·Na Trihydrate | No hit | No hit | Farnesyl pyrophosphate synthase inhibitor (bisphosphonate) | Bisphosphonate | | Bone resorption inhibitor | 1995 |
| 23 | Alfuzosin | No hit | No hit | α1-adrenoceptor antagonist | Adrenergic receptor antagonist | | Muscle relaxant | 2003 |
| 24 | Alitretinoin | Toxic | Not screened | Retinoid receptor agonist | Retinoid receptor agonist | | Antineoplastic | Unknown |
| 25 | Allopurinol | No hit | No hit | Xanthine oxidase inhibitor | Xanthine oxidase inhibitor | | Antiurolithic | 1966 |
| 26 | Almotriptan malate | No hit | No hit | 5-HT1B/1D receptor agonist | 5-HT receptor modulator | | Antimigraine | Unknown |

| | | | | | | | | |
|---|---|---|---|---|---|---|---|---|
| 27 | Alosetron·HCl | No hit | No hit | 5-HT3 receptor antagonist | 5-HT receptor modulator | | Antidiarrheal | 2000 |
| 28 | Alprostadil | No hit | No hit | Prostaglandin EP receptor agonist (PGE1) | Prostaglandin receptor agonist | | Vasodilator | 1995 |
| 29 | Altretamine | No hit | No hit | DNA alkylating agent | DNA alkylating agent | | Antineoplastic | 1990 |
| 30 | Amantadine·HCl | No hit | No hit | Viral M2 ion channel inhibitor; also promotes dopamine release | Viral ion channel inhibitor | Dopaminergic potentiator | Antiviral / Antiparkinsonian | 1986 |
| 31 | Ambrisentan | No hit | No hit | Endothelin ETA receptor antagonist | Vasoactive modulator | | Pulmonary antihypertensive | 2007 |
| 32 | Amcinonide | No hit | No hit | Glucocorticoid receptor agonist | Glucocorticoid receptor agonist | | Anti-inflammatory | 1979 |
| 33 | Amifostine | No hit | No hit | Radioprotective (free radical scavenger) | Radioprotective agent | | Radioprotective agent | 1995 |
| 34 | Amikacin Disulfate | No hit | Not screened | 30S ribosomal subunit inhibitor | Ribosomal subunit inhibitor | | Antibiotic | 1993 |
| 35 | Amiloride·HCl·2H2O | No hit | No hit | Epithelial $Na^+$ channel (ENaC) blocker | Diuretic | | Diuretic | 1981 |
| 36 | Aminocaproic Acid | No hit | No hit | Antifibrinolytic (plasminogen inhibitor) | Antifibrinolytic | | Hemostatic | 1964 |
| 37 | Aminohippurate·Na | No hit | No hit | Renal function diagnostic agent | Diagnostic agent | | Diagnostic agent | 1944 |
| 38 | Aminolevulinic Acid·HCl | No hit | No hit | Heme biosynthesis intermediate / photosensitiser | Heme biosynthesis modulator | | Antineoplastic | Unknown |
| 39 | Aminophylline | No hit | No hit | Phosphodiesterase (PDE) inhibitor; Adenosine receptor antagonist | Phosphodiesterase (PDE) inhibitor | Adenosine receptor modulator | Bronchodilator | 1982 |
| 40 | Amiodarone·HCl | Toxic | Not screened | Class III antiarrhythmic; voltage-gated $K^+$ channel blocker | Voltage-gated $K^+$ channel blocker | Adrenergic receptor antagonist; Thyroid modulator | Antiarrhythmic | 1985 |

| | | | | | | | | |
|---|---|---|---|---|---|---|---|---|
| 41 | Amitriptyline·HCl | Toxic | Not screened | NE / 5-HT / DA reuptake inhibitor | Serotonin reuptake inhibitor (SSRI/SNRI) | Adrenergic receptor antagonist; H1 receptor antagonist; Muscarinic receptor antagonist | Antidepressant | 1961 |
| 42 | Amlexanox | No hit | Not confirmed in 2nd round | TBK1 / IKKε inhibitor; mast cell stabiliser | Innate immune modulator | | Antihistamine | 1996 |
| 43 | Amlodipine | No hit | No hit | Voltage-gated $Ca^{2+}$ channel (VGCC) blocker | Voltage-gated $Ca^{2+}$ channel (VGCC) blocker | | Antihypertensive | Unknown |
| 44 | Amoxapine | Toxic | Not screened | NE / 5-HT / DA reuptake inhibitor | Serotonin reuptake inhibitor (SSRI/SNRI) | Dopaminergic inhibitor; Muscarinic receptor antagonist | Antidepressant | 1980 |
| 45 | Amoxicillin | No hit | Not screened | Penicillin-binding protein inhibitor (β-lactam) | β-lactam (PBP inhibitor) | | Antibiotic | 1974 |
| 46 | Amphotericin B | No hit | No hit | Ergosterol-binding polyene antifungal | Ergosterol-binding antifungal | | Antifungal | 1992 |
| 47 | Ampicillin Trihydrate | No hit | No hit | Penicillin-binding protein inhibitor (β-lactam) | β-lactam (PBP inhibitor) | | Antibiotic | 1998 |
| 48 | Amrinone | Hit | Hit | Phosphodiesterase 3 (PDE3) inhibitor | Phosphodiesterase (PDE) inhibitor | | Vasodilator | 1984 |
| 49 | Anagrelide | No hit | No hit | Phosphodiesterase 3 (PDE3) inhibitor | Phosphodiesterase (PDE) inhibitor | | Antiplatelet | 1997 |
| 50 | Anastrozole | No hit | No hit | Aromatase inhibitor | Aromatase inhibitor | | Antineoplastic | 1995 |
| 51 | Apomorphine·HCl Hemihydrate | Toxic | Not screened | Dopamine receptor agonist | Dopaminergic potentiator | | Antiparkinsonian | 2004 |
| 52 | Aprepitant | No hit | Not screened | NK1 receptor antagonist | NK1 receptor antagonist | | Antiemetic | 2003 |
| 53 | Argatroban | No hit | No hit | Direct thrombin inhibitor | Antiplatelet / Anticoagulant | | Anticoagulant | 2000 |

| | | | | | | | | |
|---|---|---|---|---|---|---|---|---|
| 54 | Aripiprazole | Toxic | Not screened | Dopamine D2 partial agonist; 5-HT1A partial agonist; 5-HT2A antagonist | Dopaminergic inhibitor | 5-HT receptor modulator | Antipsychotic | 2002 |
| 55 | Arsenic Trioxide | No hit | No hit | PML-RARA degradation inducer; thioredoxin reductase inhibitor | Thioredoxin reductase inhibitor | | Antineoplastic | 2000 |
| 56 | Artemether | No hit | Not screened | Reactive oxygen species generator (haem-activated) | Antimalarial | | Antimalarial | 2009 |
| 57 | Articaine·HCl | No hit | No hit | Voltage-gated $Na^+$ channel blocker | Voltage-gated $Na^+$ channel blocker | | Anesthetic | 2000 |
| 58 | Asenapine Maleate | No hit | No hit | Dopamine D2 receptor antagonist; H1 receptor antagonist; α1-adrenoceptor antagonist; 5-HT1A receptor antagonist | Dopaminergic inhibitor | 5-HT receptor modulator; Adrenergic receptor antagonist; H1 receptor antagonist | Antipsychotic | 2009 |
| 59 | Aspirin (Acetylsalicylic Acid) | No hit | No hit | COX inhibitor (NSAID) | COX inhibitor (NSAID) | | Analgesic | Unknown |
| 60 | Atazanavir | No hit | No hit | HIV protease inhibitor | HIV protease inhibitor | | Antiviral | 2003 |
| 61 | Atomoxetine·HCl | No hit | No hit | Norepinephrine reuptake inhibitor (NET selective) | Monoamine reuptake inhibitor (NE/DA) | | Nootropic / Antidepressant | 2002 |
| 62 | Atorvastatin Calcium | No hit | No hit | HMG-CoA reductase inhibitor (statin) | HMG-CoA reductase inhibitor (statin) | | Lipid-lowering agent | 1996 |
| 63 | Atovaquone | No hit | Toxic | Cytochrome bc1 complex inhibitor | Antimalarial | | Antimalarial | 1992 |
| 64 | Atracurium Besylate | No hit | No hit | Nicotinic receptor antagonist (NMJ) | Cholinergic antagonist | | Neuromuscular blocking agent | 1983 |
| 65 | Atropine Sulfate Monohydrate | No hit | No hit | Muscarinic receptor antagonist | Muscarinic receptor antagonist | | Antispasmodic | Unknown |
| 66 | Auranofin | Toxic | Not screened | Thioredoxin reductase inhibitor | Thioredoxin reductase inhibitor | | Antirheumatic | 1985 |

| | | | | | | | | |
|---|---|---|---|---|---|---|---|---|
| 67 | Azacitidine | No hit | No hit | DNA methyltransferase inhibitor | DNA methyltransferase inhibitor | | Antineoplastic | 2004 |
| 68 | Azathioprine | No hit | No hit | Purine synthesis inhibitor | Antimetabolite | | Immunosuppressant | 1968 |
| 69 | Azelaic Acid | No hit | No hit | Tyrosinase / thioredoxin reductase inhibitor | Thioredoxin reductase inhibitor | | Dermatologic agent | 1995 |
| 70 | Azelastine·HCl | No hit | No hit | H1 receptor antagonist (antihistamine) | H1 receptor antagonist | Leukotriene / arachidonate pathway | Antihistamine | 1996 |
| 71 | Azithromycin | Not screened | No hit | 50S ribosomal subunit inhibitor | Ribosomal subunit inhibitor | | Antibiotic | 1991 |
| 72 | Aztreonam | Not screened | No hit | Penicillin-binding protein inhibitor (β-lactam) | β-lactam (PBP inhibitor) | | Antibiotic | 1986 |
| 73 | Bacitracin | No hit | No hit | Bactoprenol recycling inhibitor | Other antimicrobial | | Antibiotic | 1948 |
| 74 | Baclofen | No hit | No hit | GABAB receptor agonist | GABAergic modulator | | Muscle relaxant | 1977 |
| 75 | Balsalazide | No hit | No hit | COX inhibitor (NSAID) | COX inhibitor (NSAID) | | Anti-inflammatory | Unknown |
| 76 | Beclomethasone Dipropionate | No hit | No hit | Glucocorticoid receptor agonist | Glucocorticoid receptor agonist | | Anti-inflammatory | Unknown |
| 77 | Benazepril·HCl | No hit | No hit | Angiotensin-converting enzyme (ACE) inhibitor | RAAS modulator | | Antihypertensive | 1991 |
| 78 | Bendamustine·HCl | No hit | Hit | DNA alkylating agent | DNA alkylating agent | | Antineoplastic | 2008 |
| 79 | Bendroflumethiazide | No hit | No hit | Thiazide diuretic (NCC inhibitor) | Diuretic | | Diuretic | 1959 |
| 80 | Benztropine Mesylate | No hit | No hit | Muscarinic receptor antagonist; also inhibits dopamine transporter | Muscarinic receptor antagonist | Monoamine reuptake inhibitor (NE/DA) | Antiparkinsonian | 1954 |
| 81 | Betaine | No hit | No hit | Homocysteine methyltransferase substrate | Other / Unclassified | | Other | 1996 |

| | | | | | | | | |
|---|---|---|---|---|---|---|---|---|
| 82 | Betamethasone | No hit | No hit | Glucocorticoid receptor agonist | Glucocorticoid receptor agonist | | Anti-inflammatory | 1961 |
| 83 | Betaxolol·HCl | No hit | No hit | β1-adrenoceptor antagonist (selective) | Adrenergic receptor antagonist | | Antihypertensive | 1985 |
| 84 | Bethanechol Chloride | Not confirmed in 2nd round | No hit | Muscarinic receptor agonist | Muscarinic receptor agonist | | Parasympathomimetic | 1948 |
| 85 | Bexarotene | No hit | Toxic | Retinoid receptor agonist | Retinoid receptor agonist | | Antineoplastic | 1999 |
| 86 | Bicalutamide | No hit | No hit | Androgen receptor antagonist | Sex hormone receptor modulator | | Antineoplastic | 1995 |
| 87 | Bimatoprost | Not confirmed in 2nd round | No hit | Prostaglandin FP receptor agonist | Prostaglandin receptor agonist | | Antihypertensive / Antiglaucomic | 2001 |
| 88 | Biperiden·HCl | No hit | No hit | Muscarinic receptor antagonist | Muscarinic receptor antagonist | | Antiparkinsonian | 1959 |
| 89 | Bisacodyl | Initial hit, not retested | No hit | Intestinal stimulant laxative | GI motility modulator | | Laxative | Unknown |
| 90 | Bisoprolol Fumarate | No hit | No hit | β1-adrenoceptor antagonist (selective) | Adrenergic receptor antagonist | | Antihypertensive | 1992 |
| 91 | Bleomycin Sulfate | No hit | No hit | DNA strand break inducer (metal chelation/ROS) | DNA strand break inducer | | Antineoplastic | 1973 |
| 92 | Bortezomib | No hit | Toxic | Proteasome inhibitor | Proteasome inhibitor | | Antineoplastic | 2003 |
| 93 | Bosentan | Not confirmed in 2nd round | No hit | Endothelin ETA/ETB receptor antagonist | Vasoactive modulator | | Antihypertensive | 2001 |
| 94 | Brimonidine | Not confirmed in 2nd round | No hit | α2-adrenoceptor agonist | Adrenergic receptor agonist | | Adrenergic agonist | 1997 |

| | | | | | | | | |
|---|---|---|---|---|---|---|---|---|
| 95 | Bromfenac | Not confirmed in 2nd round | Not screened | COX inhibitor (NSAID) | COX inhibitor (NSAID) | | Analgesic | 2005 |
| 96 | Bromocriptine Mesylate | Toxic | Not screened | Dopamine receptor agonist | Dopaminergic potentiator | 5-HT receptor modulator; Adrenergic receptor antagonist | Antiparkinsonian | 1982 |
| 97 | Brompheniramine Maleate | No hit | No hit | H1 receptor antagonist (antihistamine) | H1 receptor antagonist | Muscarinic receptor antagonist | Muscarinic receptor antagonist; Antiallergic | 1957 |
| 98 | Budesonide | No hit | No hit | Glucocorticoid receptor agonist | Glucocorticoid receptor agonist | | Anti-inflammatory | 1997 |
| 99 | Bumetanide | No hit | No hit | Loop diuretic (NKCC2 inhibitor) | Diuretic | | Diuretic | 1994 |
| 100 | Bupivacaine·HCl | No hit | Not screened | Voltage-gated $Na^+$ channel blocker | Voltage-gated $Na^+$ channel blocker | | Anesthetic | 1972 |
| 101 | Bupropion | No hit | No hit | NE / DA reuptake inhibitor | Monoamine reuptake inhibitor (NE/DA) | | Antidepressant | 2000's |
| 102 | Buspirone·HCl | No hit | No hit | 5-HT1A receptor agonist | 5-HT receptor modulator | | Anxiolytic | 1986 |
| 103 | Busulfan | No hit | No hit | DNA alkylating agent | DNA alkylating agent | | Antineoplastic | 1954 |
| 104 | Butenafine·HCl | Toxic | Not screened | Squalene epoxidase inhibitor | Ergosterol synthesis inhibitor | | Antifungal | 1996 |
| 105 | Butoconazole Nitrate | Toxic | Not screened | Ergosterol synthesis inhibitor (azole) | Ergosterol synthesis inhibitor | | Antifungal | 1995 |
| 106 | Cabergoline | No hit | No hit | Dopamine receptor agonist | Dopaminergic potentiator | 5-HT receptor modulator | Antiparkinsonian | 1996 |
| 107 | Caffeine | No hit | No hit | Adenosine receptor antagonist; Phosphodiesterase (PDE) inhibitor | Adenosine receptor modulator | Phosphodiesterase (PDE) inhibitor | Central Nervous System Stimulant | 1999 |
| 108 | Calcipotriene | No hit | Toxic | Vitamin D receptor agonist | Vitamin D receptor agonist | | Dermatologic agent | 1993 |

| | | | | | | | | |
|---|---|---|---|---|---|---|---|---|
| 109 | Calcitriol | Toxic | Not screened | Vitamin D receptor agonist | Vitamin D receptor agonist | | Antihypocalcemic agent | 1978 |
| 110 | Candesartan | No hit | No hit | Angiotensin II receptor antagonist (ARB) | RAAS modulator | | Antihypertensive | Unknown |
| 111 | Capecitabine | No hit | Not screened | Thymidylate synthase inhibitor (5-FU prodrug) | Antimetabolite | | Antineoplastic | 1998 |
| 112 | Capreomycin Disulfate | No hit | Not screened | 70S ribosome inhibitor | Ribosomal subunit inhibitor | | Antibiotic | 1971 |
| 113 | Capsaicin | No hit | No hit | TRPV1 receptor agonist | TRPV1 receptor agonist | | Analgesic | 1999 |
| 114 | Captopril | No hit | No hit | Angiotensin-converting enzyme (ACE) inhibitor | RAAS modulator | | Antihypertensive | 1981 |
| 115 | Carbachol (Carbamylcholine ) Chloride | No hit | Not confirmed in 2nd round | Muscarinic receptor agonist | Muscarinic receptor agonist | | Parasympathomimetic | 1972 |
| 116 | Carbamazepine | No hit | No hit | Voltage-gated $Na^+$ channel blocker | Voltage-gated $Na^+$ channel blocker | | Anticonvulsant | 1968 |
| 117 | Carbinoxamine Maleate | No hit | No hit | H1 receptor antagonist (antihistamine) | H1 receptor antagonist | Muscarinic receptor antagonist | Antihistamine | 1954 |
| 118 | Carboplatin | No hit | No hit | DNA crosslinker (platinum) | DNA crosslinker | | Antineoplastic | 1989 |
| 119 | Carglumic Acid | No hit | Not confirmed in 2nd round | Carbamoyl phosphate synthetase 1 activator | Urea cycle modulator | | Other | 2010 |
| 120 | Carmustine | No hit | No hit | DNA alkylating agent | DNA alkylating agent | | Antineoplastic | 1977 |
| 121 | Carvedilol | Toxic | Not screened | β-adrenoceptor antagonist; also α1-adrenoceptor antagonist | Adrenergic receptor antagonist | | Antihypertensive | 1995 |
| 122 | Cefaclor | No hit | No hit | Penicillin-binding protein inhibitor (β-lactam) | β-lactam (PBP inhibitor) | | Antibiotic | 1979 |
| 123 | Cefadroxil Monohydrate | No hit | No hit | Penicillin-binding protein inhibitor (β-lactam) | β-lactam (PBP inhibitor) | | Antibiotic | 1978 |

| | | | | | | | | |
|---|---|---|---|---|---|---|---|---|
| 124 | Cefazolin·Na | No hit | No hit | Penicillin-binding protein inhibitor (β-lactam) | β-lactam (PBP inhibitor) | | Antibiotic | 1973 |
| 125 | Cefdinir | No hit | Not confirmed in 2nd round | Penicillin-binding protein inhibitor (β-lactam) | β-lactam (PBP inhibitor) | | Antibiotic | 1997 |
| 126 | Cefditoren Pivoxil | No hit | No hit | Penicillin-binding protein inhibitor (β-lactam) | β-lactam (PBP inhibitor) | | Antibiotic | 2001 |
| 127 | Cefepime·2HCl Hydrate | No hit | No hit | Penicillin-binding protein inhibitor (β-lactam) | β-lactam (PBP inhibitor) | | Antibiotic | 1996 |
| 128 | Cefixime | No hit | No hit | Penicillin-binding protein inhibitor (β-lactam) | β-lactam (PBP inhibitor) | | Antibiotic | 1989 |
| 129 | Cefotaxime Acid | No hit | No hit | Penicillin-binding protein inhibitor (β-lactam) | β-lactam (PBP inhibitor) | | Antibiotic | Unknown |
| 130 | Cefotetan Disodium | No hit | Not confirmed in 2nd round | Penicillin-binding protein inhibitor (β-lactam) | β-lactam (PBP inhibitor) | | Antibiotic | 1985 |
| 131 | Cefoxitin·Na | No hit | No hit | Penicillin-binding protein inhibitor (β-lactam) | β-lactam (PBP inhibitor) | | Antibiotic | 1978 |
| 132 | Cefpodoxime Proxetil | No hit | No hit | Penicillin-binding protein inhibitor (β-lactam) | β-lactam (PBP inhibitor) | | Antibiotic | 1992 |
| 133 | Cefprozil | No hit | No hit | Penicillin-binding protein inhibitor (β-lactam) | β-lactam (PBP inhibitor) | | Antibiotic | 1991 |
| 134 | Ceftazidime | No hit | No hit | Penicillin-binding protein inhibitor (β-lactam) | β-lactam (PBP inhibitor) | | Antibiotic | 1985 |
| 135 | Ceftibuten | No hit | No hit | Penicillin-binding protein inhibitor (β-lactam) | β-lactam (PBP inhibitor) | | Antibiotic | 2009 |
| 136 | Ceftizoxim·Na | Hit | No hit | Penicillin-binding protein inhibitor (β-lactam) | β-lactam (PBP inhibitor) | | Antibiotic | Unknown |
| 137 | Ceftriaxone·Na | No hit | No hit | Penicillin-binding protein inhibitor (β-lactam) | β-lactam (PBP inhibitor) | | Antibiotic | 1984 |
| 138 | Cefuroxime Axetil | No hit | No hit | Penicillin-binding protein inhibitor (β-lactam) | β-lactam (PBP inhibitor) | | Antibiotic | 1987 |

| | | | | | | | | |
|---|---|---|---|---|---|---|---|---|
| 139 | Cefuroxime·Na | No hit | No hit | Penicillin-binding protein inhibitor (β-lactam) | β-lactam (PBP inhibitor) | | Antibiotic | 1983 |
| 140 | Celecoxib | No hit | No hit | COX inhibitor (NSAID) | COX inhibitor (NSAID) | | NSAID | 1998 |
| 141 | Cephalexin Monohydrate | No hit | No hit | Penicillin-binding protein inhibitor (β-lactam) | β-lactam (PBP inhibitor) | | Antibiotic | Unknown |
| 142 | Cetirizine 2HCl | No hit | No hit | H1 receptor antagonist (antihistamine) | H1 receptor antagonist | | Antihistamine | 1995 |
| 143 | Chenodiol (Chenodeoxycholic Acid) | No hit | No hit | Bile acid (cholesterol dissolution) | Other / Unclassified | | Cholagogue | 1983 |
| 144 | Chlorambucil | No hit | No hit | DNA alkylating agent | DNA alkylating agent | | Antineoplastic | 1957 |
| 145 | Chloramphenicol | Not confirmed in 2nd round | No hit | 50S ribosomal subunit inhibitor | Ribosomal subunit inhibitor | | Antibiotic | 1949 |
| 146 | Chlorhexidine Dihydrochloride | Not screened | Toxic | Bacterial cell membrane disruptor | Bacterial cell membrane disruptor | | Antiseptic | 1976 |
| 147 | Chloroquine Diphosphate | No hit | Not screened | Heme polymerisation inhibitor | Antimalarial | | Antimalarial | 1949 |
| 148 | Chlorothiazide | No hit | No hit | Thiazide diuretic (NCC inhibitor) | Diuretic | | Diuretic | 1958 |
| 149 | Chlorpheniramine Maleate | No hit | No hit | H1 receptor antagonist (antihistamine) | H1 receptor antagonist | Muscarinic receptor antagonist | Antihistamine | Unknown |
| 150 | Chlorpromazine·HCl | Toxic | Not screened | Dopamine D2 receptor antagonist | Dopaminergic inhibitor | 5-HT receptor modulator; Adrenergic receptor antagonist; H1 receptor antagonist; Muscarinic receptor antagonist | Antipsychotic | 1957 |
| 151 | Chlorpropamide | No hit | No hit | ATP-sensitive $K^+$ channel blocker (insulin secretagogue) | ATP-sensitive $K^+$ channel modulator | | Antidiabetic | 1958 |

| | | | | | | | | |
|---|---|---|---|---|---|---|---|---|
| 152 | Chlorthalidone | No hit | No hit | Thiazide-like diuretic (NCC inhibitor) | Diuretic | | Diuretic | 1960 |
| 153 | Chlorzoxazone | No hit | No hit | Centrally acting muscle relaxant (BK channel activator) | Other / Unclassified | | Muscle relaxant | 1958 |
| 154 | Ciclesonide | Toxic | Toxic | Glucocorticoid receptor agonist | Glucocorticoid receptor agonist | | Glucocorticoid | Unknown |
| 155 | Ciclopirox | Not screened | Toxic | Metal chelator (antifungal) | Other antifungal | | Antifungal | 1982 |
| 156 | Cidofovir | No hit | Not screened | Viral DNA polymerase inhibitor | Viral DNA polymerase inhibitor | | Antiviral | 1996 |
| 157 | Cilastatin·Na | No hit | No hit | Renal dehydropeptidase inhibitor | Other antimicrobial | | Antibiotic adjuvant | 1985 |
| 158 | Cilostazol | No hit | No hit | Phosphodiesterase 3 (PDE3) inhibitor; adenosine uptake inhibitor | Phosphodiesterase (PDE) inhibitor | Adenosine receptor modulator | Vasodilator / Antiplatelet | 1999 |
| 159 | Cimetidine | No hit | Toxic | H2 receptor antagonist | H2 receptor antagonist | | Antiulcer | 1977 |
| 160 | Cinacalcet·HCl | Not screened | Toxic | Calcium-sensing receptor positive allosteric modulator | Calcium-sensing receptor modulator | | Calcimimetic | 2004 |
| 161 | Ciprofloxacin | No hit | No hit | DNA gyrase / topoisomerase IV inhibitor | DNA gyrase / topoisomerase IV inhibitor | | Antibiotic | 1987 |
| 162 | Cisatracurium Besylate | No hit | No hit | Nicotinic receptor antagonist (NMJ) | Cholinergic antagonist | | Neuromuscular blocking agent | 1995 |
| 163 | Cisplatin (Cis-Diamineplatinum(Ii) Dichloride ) | No hit | No hit | DNA crosslinker (platinum) | DNA crosslinker | | Antineoplastic | Unknown |
| 164 | Citalopram·HBr | No hit | No hit | Serotonin reuptake inhibitor (SSRI/SNRI) | Serotonin reuptake inhibitor (SSRI/SNRI) | | Antidepressant | 1998 |
| 165 | Cladribine | No hit | No hit | Purine nucleoside analogue (adenosine deaminase resistant) | Antimetabolite | | Antineoplastic | 1993 |

| | | | | | | | | |
|---|---|---|---|---|---|---|---|---|
| 166 | Clarithromycin | No hit | No hit | 50S ribosomal subunit inhibitor | Ribosomal subunit inhibitor | | Antibiotic | 1991 |
| 167 | Clavulanate Potassium | No hit | No hit | β-lactamase inhibitor | Other antimicrobial | | Antibiotic adjuvant | 1984 |
| 168 | Clemastine Fumarate | Toxic | Not screened | H1 receptor antagonist (antihistamine) | H1 receptor antagonist | Muscarinic receptor antagonist | Antihistamine | 1977 |
| 169 | Clindamycin Palmitate·HCl | No hit | No hit | 50S ribosomal subunit inhibitor | Ribosomal subunit inhibitor | | Antibiotic | 1980 |
| 170 | Clindamycin·HCl | No hit | No hit | 50S ribosomal subunit inhibitor | Ribosomal subunit inhibitor | | Antibiotic | 1970 |
| 171 | Clobetasol Propionate | Not confirmed in 2nd round | No hit | Glucocorticoid receptor agonist | Glucocorticoid receptor agonist | | Anti-inflammatory | 1985 |
| 172 | Clofarabine | No hit | No hit | Purine nucleoside analogue; ribonucleotide reductase inhibitor | Antimetabolite | | Antineoplastic | 2004 |
| 173 | Clofazimine | Not screened | Toxic | Mycobacterial membrane redox cycling agent | Bacterial DNA template inhibitor | | Antibacterial | 1986 |
| 174 | Clomiphene Citrate | Toxic | Not screened | Estrogen receptor modulator | Sex hormone receptor modulator | | Fertility agent | 1967 |
| 175 | Clomipramine·HCl | Toxic | Not screened | NE / 5-HT / DA reuptake inhibitor | Serotonin reuptake inhibitor (SSRI/SNRI) | H1 receptor antagonist; Muscarinic receptor antagonist | Antidepressant | 1989 |
| 176 | Clonidine·HCl | No hit | No hit | α2-adrenoceptor agonist | Adrenergic receptor agonist | | Antihypertensive | Unknown |
| 177 | Clopidogrel Bisulfate | No hit | No hit | P2Y12 receptor antagonist | Antiplatelet / Anticoagulant | | Antiplatelet | 1997 |
| 178 | Clotrimazole | Toxic | Not screened | Ergosterol synthesis inhibitor (azole) | Ergosterol synthesis inhibitor | | Antifungal | 1975 |
| 179 | Cloxacillin·Na | No hit | Not screened | Penicillin-binding protein inhibitor (β-lactam) | β-lactam (PBP inhibitor) | | Antibiotic | Unknown |

| | | | | | | | | |
|---|---|---|---|---|---|---|---|---|
| 180 | Clozapine | No hit | Toxic | Dopamine receptor antagonist; 5-HT2 receptor antagonist | Dopaminergic inhibitor | 5-HT receptor modulator; Adrenergic receptor antagonist; H1 receptor antagonist; Muscarinic receptor antagonist | Antipsychotic | 1989 |
| 181 | Colchicine | No hit | Not screened | Microtubule destabiliser | Microtubule modulator | | Anti-gout | 1961 |
| 182 | Colistimethate·Na | No hit | Not screened | Bacterial cell membrane disruptor | Bacterial cell membrane disruptor | | Antibiotic | 1970 |
| 183 | Colistin Sulfate | Toxic | Not screened | Bacterial cell membrane disruptor | Bacterial cell membrane disruptor | | Antibiotic | 1962 |
| 184 | Cortisone Acetate | No hit | Not screened | Glucocorticoid receptor agonist | Glucocorticoid receptor agonist | | Anti-inflammatory | 1950 |
| 185 | Cromolyn·Na (Disodium Cromoglycate) | No hit | No hit | Mast cell stabiliser | Leukotriene / arachidonate pathway | | Mast cell stabilizer | 1982 |
| 186 | Crotamiton | Initial hit, not retested | No hit | Protease-activated receptor 2 (PAR-2) antagonist; H1 receptor antagonist; 5-HT2 receptor antagonist | Other / Unclassified | 5-HT receptor modulator; H1 receptor antagonist | Scabicidal | 1949 |
| 187 | Cyclobenzaprine·HCl | No hit | Not screened | 5-HT2 receptor antagonist | 5-HT receptor modulator | H1 receptor antagonist; Muscarinic receptor antagonist | Muscle relaxant | 1977 |
| 188 | Cyclopentolate | No hit | Not screened | Muscarinic receptor antagonist | Muscarinic receptor antagonist | | Mydriatic | Unknown |
| 189 | Cyclophosphamide monohydrate | No hit | No hit | DNA alkylating agent | DNA alkylating agent | | Antineoplastic | 1959 |
| 190 | Cycloserine | No hit | Not screened | NMDA receptor glycine-site partial agonist; bacterial alanine racemase inhibitor | Mycobacterial cell wall inhibitor | NMDA receptor antagonist | Antibiotic | 1964 |

| | | | | | | | | |
|---|---|---|---|---|---|---|---|---|
| 191 | Cyclosporine A | No hit | No hit | Calcineurin inhibitor | Calcineurin inhibitor | | Immunosuppressant | 1983 |
| 192 | Cyproheptadine·HCl Sesquihydrate | Toxic | Not screened | H1 receptor antagonist (antihistamine); also serotonin antagonist | H1 receptor antagonist | 5-HT receptor modulator; Muscarinic receptor antagonist | Antihistamine | 1961 |
| 193 | Cysteamine·HCl | No hit | Not screened | Cystine-depleting agent | Other / Unclassified | | Cystine-depleting agent | 1994 |
| 194 | Cytarabine | No hit | No hit | DNA polymerase inhibitor / chain terminator | Antimetabolite | | Antineoplastic | 1969 |
| 195 | Dacarbazine | No hit | No hit | DNA alkylating agent | DNA alkylating agent | | Antineoplastic | 1975 |
| 196 | Dactinomycin (= Actinomycin D) | No hit | Not screened | DNA intercalator; RNA polymerase inhibitor | RNA polymerase inhibitor | | Antineoplastic | 1964 |
| 197 | Dalfampridine (4-Aminopyridine) | No hit | Not screened | Voltage-gated $K^+$ channel blocker | Voltage-gated $K^+$ channel blocker | | Potassium channel blocker | 2010 |
| 198 | Danazol | Toxic | Not screened | Gonadotropin inhibitor; binds androgen | Sex hormone receptor modulator | Glucocorticoid receptor agonist | Estrogen Antagonist | 1976 |
| 199 | Dantrolene·Na | No hit | Not screened | Ryanodine receptor inhibitor | Ryanodine receptor inhibitor | | Muscle relaxant | 1974 |
| 200 | Dapsone | No hit | Not screened | Bacterial dihydropteroate synthetase (DHFR) inhibitor; also inhibits myeloperoxidase and LTB4 | Folate pathway inhibitor | DHFR / folate synthesis inhibitor; Myeloperoxidase inhibitor | Antibacterial | Unknown |
| 201 | Daptomycin | No hit | Not screened | Bacterial cell membrane disruptor | Bacterial cell membrane disruptor | | Antibiotic | 2003 |
| 202 | Darifenacin·HBr | No hit | Not screened | Muscarinic receptor antagonist | Muscarinic receptor antagonist | | Overactive bladder treatment | 2004 |
| 203 | Darunavir | No hit | Not screened | HIV protease inhibitor | HIV protease inhibitor | | Antiviral | 2006 |
| 204 | Dasatinib | Toxic | Not screened | Tyrosine kinase inhibitor | Tyrosine kinase inhibitor | | Antineoplastic | 2006 |

| | | | | | | | | |
|---|---|---|---|---|---|---|---|---|
| 205 | Daunorubicin·HCl | No hit | No hit | Topoisomerase II inhibitor; DNA intercalator | Topoisomerase inhibitor | | Antineoplastic | 1979 |
| 206 | Decitabine | No hit | Not screened | DNA methyltransferase inhibitor | DNA methyltransferase inhibitor | | Antineoplastic | 2006 |
| 207 | Deferasirox | No hit | Not screened | Iron chelator | Iron chelator | | Iron chelator | 2005 |
| 208 | Deferoxamine Mesylate | No hit | Not screened | Iron chelator | Iron chelator | | Iron chelator | 1968 |
| 209 | Delavirdine Mesylate | No hit | No hit | Non-nucleoside reverse transcriptase inhibitor (NNRTI) | Reverse transcriptase inhibitor | | Antiviral | 1997 |
| 210 | Demeclocycline·HCl | No hit | Not screened | 30S ribosomal subunit inhibitor | Ribosomal subunit inhibitor | | Antibiotic | 1960 |
| 211 | Desipramine·HCl | No hit | Not screened | NE / 5-HT / DA reuptake inhibitor | Serotonin reuptake inhibitor (SSRI/SNRI) | Muscarinic receptor antagonist | Antidepressant | 1964 |
| 212 | Desloratadine | Not confirmed in 2nd round | No hit | H1 receptor antagonist (antihistamine) | H1 receptor antagonist | | Antihistamine | 2001 |
| 213 | Desogestrel | Toxic | Not screened | Estrogen receptor modulator; Progesterone receptor agonist | Sex hormone receptor modulator | | Contraceptive | 1992 |
| 214 | Desonide | No hit | Not screened | Glucocorticoid receptor agonist | Glucocorticoid receptor agonist | | Anti-inflammatory | 1994 |
| 215 | Desoximetasone | No hit | Not screened | Glucocorticoid receptor agonist | Glucocorticoid receptor agonist | | Anti-inflammatory | 1977 |
| 216 | Desvenlafaxine Succinate Hydrate | No hit | Not screened | Serotonin reuptake inhibitor (SSRI/SNRI) | Serotonin reuptake inhibitor (SSRI/SNRI) | | Antidepressant | 2008 |
| 217 | Dexamethasone | No hit | No hit | Glucocorticoid receptor agonist | Glucocorticoid receptor agonist | | Anti-inflammatory | 1958 |
| 218 | Dexchlorpheniramine Maleate | No hit | Not screened | H1 receptor antagonist (antihistamine) | H1 receptor antagonist | Muscarinic receptor antagonist | Antihistamine | 1981 |

| | | | | | | | | |
|---|---|---|---|---|---|---|---|---|
| 219 | Dexmedetomidine·HCl | Toxic | Not screened | α2-adrenoceptor agonist | Adrenergic receptor agonist | | Sedative | 1999 |
| 220 | Dexrazoxane | Not confirmed in 2nd round | No hit | Iron chelator; catalytic topoisomerase II inhibitor | Iron chelator | | Cardioprotective agent | 1995 |
| 221 | Dextromethorphan | No hit | No hit | N-methyl-D-aspartate (NMDA) receptor antagonist; also sigma-1 agonist | NMDA receptor antagonist | | Antitussive | 1950's |
| 222 | Diatrizoate Meglumine | No hit | Not screened | Iodinated radiocontrast agent | Diagnostic agent | | Contrast agent | 1954 |
| 223 | Diazoxide | No hit | No hit | ATP-sensitive $K^+$ channel opener | ATP-sensitive $K^+$ channel modulator | | Vasodilator | 1973 |
| 224 | Diclofenac·Na Salt | No hit | No hit | COX inhibitor (NSAID); also blocks voltage-gated sodium channels | COX inhibitor (NSAID) | | NSAID | 1988 |
| 225 | Dicloxacillin·Na Salt Monohydrate | No hit | Not screened | Penicillin-binding protein inhibitor (β-lactam) | β-lactam (PBP inhibitor) | | Antibiotic | 1968 |
| 226 | Dicyclomine·HCl | Hit | Hit | Muscarinic receptor antagonist | Muscarinic receptor antagonist | | Antispasmodic | 1950 |
| 227 | Didanosine | No hit | No hit | Nucleoside Reverse Transcriptase Inhibitor (NRTI) | Reverse transcriptase inhibitor | | Antiviral | 1991 |
| 228 | Dienogest | Not confirmed in 2nd round | Not confirmed in 2nd round | Progesterone receptor agonist | Sex hormone receptor modulator | | Contraceptive | 2010 |
| 229 | Diflunisal | No hit | No hit | COX inhibitor (NSAID) | COX inhibitor (NSAID) | | NSAID | 1982 |
| 230 | Difluprednate | No hit | Not screened | Glucocorticoid receptor agonist | Glucocorticoid receptor agonist | | Corticosteroid | 2008 |
| 231 | Digoxin | No hit | Not screened | $Na^+/K^+$-ATPase inhibitor (cardiac glycoside) | $Na^+/K^+$-ATPase inhibitor (cardiac glycoside) | | Antiarrhythmic | Unknown |

| | | | | | | | | |
|---|---|---|---|---|---|---|---|---|
| 232 | Dihydroergotamine Mesylate | No hit | No hit | 5-HT1B/1D receptor agonist | 5-HT receptor modulator | Adrenergic receptor agonist; Dopaminergic potentiator | Antimigraine | 1946 |
| 233 | Diltiazem·HCl | No hit | No hit | Voltage-gated $Ca^{2+}$ channel (VGCC) blocker | Voltage-gated $Ca^{2+}$ channel (VGCC) blocker | | Antihypertensive | 1982 |
| 234 | Dimenhydrinate | No hit | Not screened | H1 receptor antagonist (antihistamine) | H1 receptor antagonist | Muscarinic receptor antagonist | Antiemetic | 1949 |
| 235 | Dinoprostone | No hit | No hit | Prostaglandin EP/FP receptor agonist | Prostaglandin receptor agonist | | Prostaglandin | 1977 |
| 236 | Diphenhydramine·HCl | No hit | No hit | H1 receptor antagonist (antihistamine) | H1 receptor antagonist | Muscarinic receptor antagonist | Antihistamine | 1946 |
| 237 | Dipyridamole | No hit | No hit | Thromboxane synthase inhibitor; Adenosine reuptake inhibitor; Phosphodiesterase (PDE) inhibitor | Antiplatelet / Anticoagulant | Adenosine receptor modulator; Phosphodiesterase (PDE) inhibitor | Vasodilator / Antiplatelet | 1961 |
| 238 | Disopyramide | Not confirmed in 2nd round | Hit | Voltage-gated $Na^{+}$ channel blocker; muscarinic receptor antagonist | Voltage-gated $Na^{+}$ channel blocker | Muscarinic receptor antagonist | Antiarrhythmic | Unknown |
| 239 | Disulfiram | Toxic | Not screened | Aldehyde dehydrogenase inhibitor | Alcohol metabolism modulator | | Alcohol Deterrent | 1951 |
| 240 | Dobutamine·HCl | No hit | No hit | β1-adrenoceptor agonist | Adrenergic receptor agonist | | Sympathomimetic | 1978 |
| 241 | Docetaxel (Taxotere) | No hit | No hit | Microtubule stabiliser | Microtubule modulator | | Antineoplastic | 1996 |
| 242 | Dofetilide | No hit | No hit | Voltage-gated $K^{+}$ channel blocker | Voltage-gated $K^{+}$ channel blocker | | Antiarrhythmic | 1999 |
| 243 | Dolasetron | No hit | No hit | 5-HT3 receptor antagonist | 5-HT receptor modulator | | Antiemetic | Unknown |
| 244 | Donepezil·HCl | No hit | No hit | Acetylcholinesterase inhibitor | Cholinergic agonist | | Parasympathomimetic | 1996 |

| | | | | | | | | |
|---|---|---|---|---|---|---|---|---|
| 245 | Dopamine·HCl | Not confirmed in 2nd round | No hit | Dopamine receptor agonist; α- and β-adrenoceptor agonist | Adrenergic receptor agonist | Dopaminergic potentiator | Neurotransmitter | 1974 |
| 246 | Doripenem | No hit | Not screened | Penicillin-binding protein inhibitor (β-lactam) | β-lactam (PBP inhibitor) | | Antibiotic | 2007 |
| 247 | Dorzolamide·HCl | No hit | No hit | Carbonic anhydrase inhibitor | Diuretic | | Antihypertensive (ocular) | 1994 |
| 248 | Doxapram·HCl H2O | No hit | Not screened | Two-pore-domain $K^+$ (K2P) channel blocker | Voltage-gated $K^+$ channel blocker | | Respiratory stimulant | 1965 |
| 249 | Doxazosin Mesylate | No hit | No hit | α1-adrenoceptor antagonist | Adrenergic receptor antagonist | | Antihypertensive | 1990 |
| 250 | Doxepin·HCl | No hit | Not screened | NE / 5-HT / DA reuptake inhibitor; H1 receptor antagonist | Serotonin reuptake inhibitor (SSRI/SNRI) | Adrenergic receptor antagonist; H1 receptor antagonist; Muscarinic receptor antagonist | Antidepressant | 1969 |
| 251 | Doxorubicin·HCl | No hit | No hit | Topoisomerase II inhibitor; DNA intercalator | Topoisomerase inhibitor | | Antineoplastic | 1974 |
| 252 | Doxycycline Monohydrate | No hit | Initial hit, not retested | 30S ribosomal subunit inhibitor | Ribosomal subunit inhibitor | | Antibiotic | 1967 |
| 253 | Droperidol | No hit | Not screened | Dopamine receptor antagonist | Dopaminergic inhibitor | | Sedative | 1970 |
| 254 | Drospirenone | No hit | Not screened | Progesterone receptor agonist | Sex hormone receptor modulator | | Contraceptive | 2001 |
| 255 | Duloxetine·HCl | Toxic | Not screened | Serotonin reuptake inhibitor (SSRI/SNRI) | Serotonin reuptake inhibitor (SSRI/SNRI) | | Antidepressant | 2004 |
| 256 | Dutasteride | No hit | Not screened | 5α-reductase inhibitor | Sex hormone receptor modulator | | 5α-reductase inhibitor | 2001 |
| 257 | Dyphylline | No hit | Not screened | Phosphodiesterase (PDE) inhibitor; | Phosphodiesterase (PDE) inhibitor | Adenosine receptor modulator | Bronchodilator | 1951 |

| | | | | | | | | |
|---|---|---|---|---|---|---|---|---|
| | | | | Adenosine receptor antagonist | | | | |
| 258 | Econazole Nitrate | Toxic | Not screened | Ergosterol synthesis inhibitor (azole) | Ergosterol synthesis inhibitor | | Antifungal | 1982 |
| 259 | Efavirenz | Toxic | Not screened | Non-nucleoside reverse transcriptase inhibitor (NNRTI) | Reverse transcriptase inhibitor | | Antiviral | 1998 |
| 260 | Eflornithine·HCl | No hit | Not screened | Ornithine decarboxylase inhibitor | Antiparasitic | | Antiprotozoal | 1990 |
| 261 | Emtricitabine | No hit | Initial hit, not retested | Nucleoside Reverse Transcriptase Inhibitor (NRTI) | Reverse transcriptase inhibitor | | Antiviral | 2003 |
| 262 | Enalapril | Initial hit, not retested | No hit | Angiotensin-converting enzyme (ACE) inhibitor | RAAS modulator | | Antihypertensive | 2000's |
| 263 | Enalaprilat Maleate | No hit | No hit | Angiotensin-converting enzyme (ACE) inhibitor | RAAS modulator | | Antihypertensive | 1985 |
| 264 | Entacapone | No hit | No hit | Catechol-O-methyltransferase (COMT) inhibitor | Dopaminergic potentiator | | Antiparkinsonian | 1999 |
| 265 | Epinastine·HCl | Not confirmed in 2nd round | Not confirmed in 2nd round | 5-HT2 receptor antagonist; H1 antagonist | 5-HT receptor modulator | H1 receptor antagonist | Antihistamine | 2003 |
| 266 | Epinephrine (L-(-)-Epinephrine-(+)-Bitartrate) | No hit | No hit | Adrenergic receptor agonist (α and β) | Adrenergic receptor agonist | | Vasoconstrictor | Unknown |
| 267 | Epirubicin·HCl | No hit | Not screened | Topoisomerase II inhibitor; DNA intercalator | Topoisomerase inhibitor | | Antineoplastic | 1999 |
| 268 | Eplerenone | No hit | Not screened | Mineralocorticoid receptor (MR) antagonist | Diuretic | | Antihypertensive | 2002 |
| 269 | Eprosartan Mesylate | No hit | No hit | Angiotensin II receptor antagonist (ARB) | RAAS modulator | | Antihypertensive | 1997 |
| 270 | Eptifibatide | No hit | Not screened | GPIIb/IIIa antagonist | Antiplatelet / Anticoagulant | | Antiplatelet | 1998 |

| | | | | | | | | |
|---|---|---|---|---|---|---|---|---|
| 271 | Ergotamine Tartrate | No hit | No hit | 5-HT1B/1D receptor agonist; α-adrenoceptor agonist | 5-HT receptor modulator | Adrenergic receptor agonist; Dopaminergic potentiator | Vasoconstrictor / Antimigraine | 1948 |
| 272 | Erlotinib | No hit | No hit | Tyrosine kinase inhibitor | Tyrosine kinase inhibitor | | Antineoplastic | 2004 |
| 273 | Erythromycin | No hit | Not screened | 50S ribosomal subunit inhibitor | Ribosomal subunit inhibitor | | Antibiotic | 1964 |
| 274 | Escitalopram | No hit | No hit | Serotonin reuptake inhibitor (SSRI/SNRI) | Serotonin reuptake inhibitor (SSRI/SNRI) | | Antidepressant | Unknown |
| 275 | Esmolol | No hit | Not screened | β1-adrenoceptor antagonist (selective) | Adrenergic receptor antagonist | | Antihypertensive | 1987 |
| 276 | Esomeprazole Potassium | No hit | No hit | $H^+/K^+$-ATPase inhibitor (PPI) | $H^+/K^+$-ATPase inhibitor (PPI) | | Antiulcer | Unknown |
| 277 | Estradiol | No hit | No hit | Estrogen receptor modulator | Sex hormone receptor modulator | | Hormonal replacement | Unknown |
| 278 | Estramustine Phosphate·Na | No hit | Not screened | DNA alkylating agent; Estradiol analog | DNA alkylating agent | Sex hormone receptor modulator | Antineoplastic | 1981 |
| 279 | Estrone | No hit | No hit | Estrogen receptor modulator | Sex hormone receptor modulator | | Estrogen | Unknown |
| 280 | Estropipate | Not confirmed in 2nd round | Not confirmed in 2nd round | Estrogen receptor modulator; OATP1B1 inhibitor | Sex hormone receptor modulator | | Hormonal replacement | 1977 |
| 281 | Eszopiclone | No hit | Not screened | GABAA receptor modulator | GABAergic modulator | | Sedative | 2004 |
| 282 | Ethacrynic Acid | No hit | No hit | Loop diuretic (NKCC2 inhibitor); also inhibits glutathione S-transferase | Diuretic | | Diuretic | 1967 |
| 283 | Ethambutol Dihydrochloride | Not confirmed in 2nd round | Hit | Arabinosyl transferase inhibitor | Mycobacterial cell wall inhibitor | | Antitubercular | 1967 |

| | | | | | | | | |
|---|---|---|---|---|---|---|---|---|
| 284 | Ethinyl Estradiol | No hit | Not screened | Estrogen receptor modulator | Sex hormone receptor modulator | | Contraceptive | 1943 |
| 285 | Ethionamide | No hit | Not screened | Mycolic acid synthesis inhibitor | Mycobacterial cell wall inhibitor | | Antitubercular | 1965 |
| 286 | Ethosuximide | Not confirmed in 2nd round | No hit | T-type $Ca^{2+}$ channel blocker | Voltage-gated $Ca^{2+}$ channel (VGCC) blocker | | Anticonvulsant | 1960 |
| 287 | Etidronate Disodium | Initial hit, not retested | No hit | Farnesyl pyrophosphate synthase inhibitor (bisphosphonate) | Bisphosphonate | | Bone resorption inhibitor | 1977 |
| 288 | Etodolac | Not confirmed in 2nd round | Hit | COX inhibitor (NSAID) | COX inhibitor (NSAID) | | NSAID | 1991 |
| 289 | Etomidate | No hit | Not screened | GABAA receptor modulator | GABAergic modulator | | Anesthetic | 1982 |
| 290 | Etonogestrel | No hit | Not screened | Progesterone receptor agonist; also estrogen receptor modulator | Sex hormone receptor modulator | | Contraceptive | 2001 |
| 291 | Etoposide | No hit | No hit | Topoisomerase II inhibitor | Topoisomerase inhibitor | | Antineoplastic | 1983 |
| 292 | Everolimus | No hit | Not screened | mTOR inhibitor | mTOR inhibitor | | Immunosuppressant | 2009 |
| 293 | Exemestane | No hit | No hit | Aromatase inhibitor | Aromatase inhibitor | | Antineoplastic | 1999 |
| 294 | Ezetimibe | No hit | Not screened | NPC1L1 inhibitor | Cholesterol absorption inhibitor | | Lipid-lowering agent | 2002 |
| 295 | Famciclovir | No hit | No hit | Viral DNA polymerase inhibitor | Viral DNA polymerase inhibitor | | Antiviral | 1994 |
| 296 | Famotidine | No hit | No hit | H2 receptor antagonist | H2 receptor antagonist | | Antiulcer | 1986 |
| 297 | Febuxostat | No hit | Not screened | Xanthine oxidase inhibitor | Xanthine oxidase inhibitor | | Antigout | 2009 |
| 298 | Felbamate | No hit | No hit | N-methyl-D-aspartate (NMDA) receptor | NMDA receptor antagonist | | Anticonvulsant | 1993 |

| | | | | | | | | |
|---|---|---|---|---|---|---|---|---|
| | | | | antagonist; also modulates GABA | | | | |
| 299 | Felodipine | Toxic | Not screened | Voltage-gated $Ca^{2+}$ channel (VGCC) blocker | Voltage-gated $Ca^{2+}$ channel (VGCC) blocker | | Antihypertensive | 1991 |
| 300 | Fenofibrate | Toxic | Not screened | PPARα agonist (fibrate) | PPAR agonist | | Lipid-lowering agent | 1993 |
| 301 | Fenoldopam Mesylate | No hit | No hit | Dopamine D1 receptor agonist | Dopaminergic potentiator | | Antihypertensive | 1997 |
| 302 | Fenoprofen calcium salt dihydrate | No hit | No hit | COX inhibitor (NSAID) | COX inhibitor (NSAID) | | Analgesic | 1976 |
| 303 | Fexofenadine·HCl | Hit | No hit | H1 receptor antagonist (antihistamine) | H1 receptor antagonist | | Antihistamine | 1996 |
| 304 | Finasteride | No hit | No hit | 5α-reductase inhibitor | Sex hormone receptor modulator | | 5α-reductase inhibitor | 1992 |
| 305 | Fingolimod | No hit | Not screened | Sphingosine kinase substrate / S1P receptor modulator | Sphingosine / S1P modulator | | Immunomodulator | 2010 |
| 306 | Flavoxate·HCl | No hit | Not screened | Muscarinic receptor antagonist | Muscarinic receptor antagonist | | Muscle relaxant | 1970 |
| 307 | Flecainide Acetate | No hit | No hit | Voltage-gated $Na^{+}$ channel blocker | Voltage-gated $Na^{+}$ channel blocker | | Antiarrhythmic | 1985 |
| 308 | Floxuridine | No hit | No hit | Thymidylate synthase inhibitor | Antimetabolite | | Antineoplastic | 1970 |
| 309 | Fluconazole | No hit | Hit | Ergosterol synthesis inhibitor (azole) | Ergosterol synthesis inhibitor | | Antifungal | 1990 |
| 310 | Flucytosine | No hit | Not screened | Fungal cytosine deaminase prodrug (5-FU) | Other antifungal | | Antifungal | 1971 |
| 311 | Fludarabine Phosphate | No hit | Not screened | Purine nucleoside analogue; DNA polymerase inhibitor | Antimetabolite | | Antineoplastic | 1991 |
| 312 | Fludrocortisone Acetate | No hit | Not screened | Glucocorticoid receptor agonist; also mineralocorticoid | Glucocorticoid receptor agonist | Mineralocorticoid receptor agonist | Hormonal replacement | 1955 |

| | | | | | | | | |
|---|---|---|---|---|---|---|---|---|
| 313 | Flumazenil | No hit | No hit | GABAA receptor modulator (antagonist) | GABAergic modulator | | Benzodiazepine antagonist | 1991 |
| 314 | Flunisolide | No hit | Not screened | Glucocorticoid receptor agonist | Glucocorticoid receptor agonist | | Anti-inflammatory | 1981 |
| 315 | Fluocinolone Acetonide | No hit | No hit | Glucocorticoid receptor agonist | Glucocorticoid receptor agonist | | Anti-inflammatory | 1963 |
| 316 | Fluocinonide | No hit | Not screened | Glucocorticoid receptor agonist | Glucocorticoid receptor agonist | | Anti-inflammatory | 1971 |
| 317 | Fluorometholone | No hit | Not screened | Glucocorticoid receptor agonist | Glucocorticoid receptor agonist | | Anti-inflammatory | 1972 |
| 318 | Fluorouracil (5-Fluorouracil) | No hit | No hit | Thymidylate synthase inhibitor | Antimetabolite | | Antineoplastic | 1970 |
| 319 | Fluoxetine·HCl | Toxic | Not screened | Serotonin reuptake inhibitor (SSRI/SNRI) | Serotonin reuptake inhibitor (SSRI/SNRI) | | Antidepressant | 1987 |
| 320 | Fluphenazine· 2HCl | Toxic | Not screened | Dopamine receptor antagonist | Dopaminergic inhibitor | | Antipsychotic | 1959 |
| 321 | Flurandrenolide | No hit | Not screened | Glucocorticoid receptor agonist | Glucocorticoid receptor agonist | | Anti-inflammatory | 1965 |
| 322 | Flurbiprofen | No hit | No hit | COX inhibitor (NSAID) | COX inhibitor (NSAID) | | NSAID | 1988 |
| 323 | Flutamide | No hit | No hit | Androgen receptor antagonist | Sex hormone receptor modulator | | Antineoplastic | 1989 |
| 324 | Fluticasone Propionate | No hit | Not screened | Glucocorticoid receptor agonist | Glucocorticoid receptor agonist | | Anti-inflammatory | 1990 |
| 325 | Fluvastatin·Na | No hit | Toxic | HMG-CoA reductase inhibitor (statin) | HMG-CoA reductase inhibitor (statin) | | Lipid-lowering agent | 1993 |
| 326 | Fluvoxamine Maleate | No hit | Not screened | Serotonin reuptake inhibitor (SSRI/SNRI) | Serotonin reuptake inhibitor (SSRI/SNRI) | | Antidepressant | 1990's |
| 327 | Fomepizole | No hit | Not screened | Alcohol dehydrogenase inhibitor | Alcohol metabolism modulator | | Antidote | 1997 |
| 328 | Formoterol | No hit | Not screened | β2-adrenoceptor agonist (long-acting) | Adrenergic receptor agonist | | Bronchodilator | 2001 |

| | | | | | | | | |
|---|---|---|---|---|---|---|---|---|
| 329 | Fosinopril·Na | No hit | No hit | Angiotensin-converting enzyme (ACE) inhibitor | RAAS modulator | | Antihypertensive | 1991 |
| 330 | Fulvestrant | Not confirmed in 2nd round | Not screened | Estrogen receptor degrader (SERD) | Sex hormone receptor modulator | | Antineoplastic | 2002 |
| 331 | Furosemide | No hit | Not confirmed in 2nd round | Loop diuretic (NKCC2 inhibitor) | Diuretic | | Diuretic | 1966 |
| 332 | Gabapentin | No hit | No hit | Voltage-gated $Ca^{2+}$ channel (VGCC) blocker (α2δ subunit) | Voltage-gated $Ca^{2+}$ channel (VGCC) blocker | | Anticonvulsant | 1993 |
| 333 | Galantamine·HBr | No hit | No hit | Acetylcholinesterase inhibitor | Cholinergic agonist | Nicotinic receptor modulator (allosteric) | Acetylcholinesterase inhibitor | Unknown |
| 334 | Ganciclovir | Initial hit, not retested | No hit | Viral DNA polymerase inhibitor | Viral DNA polymerase inhibitor | | Antiviral | 1989 |
| 335 | Gatifloxacin | No hit | No hit | DNA gyrase / topoisomerase IV inhibitor | DNA gyrase / topoisomerase IV inhibitor | | Antibiotic | 1999 |
| 336 | Gefitinib | No hit | No hit | Tyrosine kinase inhibitor | Tyrosine kinase inhibitor | | Antineoplastic | 2003 |
| 337 | Gemcitabine·HCl | No hit | No hit | Ribonucleotide reductase inhibitor; DNA chain terminator | Antimetabolite | | Antineoplastic | 1996 |
| 338 | Gemfibrozil | No hit | No hit | PPARα agonist (fibrate) | PPAR agonist | | Lipid-lowering agent | 1981 |
| 339 | Gentamycin Sulfate | No hit | No hit | 30S ribosomal subunit inhibitor | Ribosomal subunit inhibitor | | Antibiotic | 1970 |
| 340 | Glimepiride | No hit | No hit | ATP-sensitive $K^+$ channel blocker (insulin secretagogue) | ATP-sensitive $K^+$ channel modulator | | Antidiabetic | 1995 |
| 341 | Glipizide | No hit | Not confirmed in 2nd round | ATP-sensitive $K^+$ channel blocker (insulin secretagogue) | ATP-sensitive $K^+$ channel modulator | | Antidiabetic | 1984 |

| | | | | | | | | |
|---|---|---|---|---|---|---|---|---|
| 342 | Glyburide | No hit | No hit | ATP-sensitive $K^+$ channel blocker (insulin secretagogue) | ATP-sensitive $K^+$ channel modulator | | Antidiabetic | 1984 |
| 343 | Glycopyrrolate Iodide | Hit | Hit | Muscarinic receptor antagonist | Muscarinic receptor antagonist | | Anticholinergic | 1961 |
| 344 | Goserelin Acetate | Initial hit, not retested | No hit | GnRH agonist | Sex hormone receptor modulator | | Antineoplastic | 1989 |
| 345 | Granisetron·HCl | No hit | No hit | 5-HT3 receptor antagonist | 5-HT receptor modulator | | Antiemetic | 1993 |
| 346 | Griseofulvin | No hit | Not screened | Fungal tubulin inhibitor | Other antifungal | | Antifungal | 1962 |
| 347 | Guanabenz Acetate | No hit | No hit | α2-adrenoceptor agonist | Adrenergic receptor agonist | | Antihypertensive | 1982 |
| 348 | Guanfacine·HCl | No hit | No hit | α2A-adrenoceptor agonist | Adrenergic receptor agonist | | Antihypertensive | 1986 |
| 349 | Guanidine·HCl | No hit | Not screened | Voltage-gated $K^+$ channel blocker | Voltage-gated $K^+$ channel blocker | | Other | 1939 |
| 350 | Haloperidol | Toxic | Not screened | Dopamine D2 receptor antagonist | Dopaminergic inhibitor | Adrenergic receptor antagonist | Antipsychotic | 1967 |
| 351 | Hydrocortisone | No hit | No hit | Glucocorticoid receptor agonist | Glucocorticoid receptor agonist | | Anti-inflammatory | Unknown |
| 352 | Hydrocortisone Acetate | No hit | No hit | Glucocorticoid receptor agonist | Glucocorticoid receptor agonist | | Anti-inflammatory | Unknown |
| 353 | Ibandronate·Na Monohydrate | No hit | No hit | Farnesyl pyrophosphate synthase inhibitor (bisphosphonate) | Bisphosphonate | | Bone resorption inhibitor | 2005 |
| 354 | Ibuprofen | No hit | Toxic | COX inhibitor (NSAID) | COX inhibitor (NSAID) | | NSAID | 1974 |
| 355 | Idarubicin·HCl | Toxic | Not screened | Topoisomerase II inhibitor; DNA intercalator | Topoisomerase inhibitor | | Antineoplastic | 1990 |
| 356 | Idoxuridine | No hit | No hit | Viral DNA polymerase inhibitor | Viral DNA polymerase inhibitor | | Antiviral | 1963 |

| | | | | | | | | |
|---|---|---|---|---|---|---|---|---|
| 357 | Ifosfamide | No hit | No hit | DNA alkylating agent | DNA alkylating agent | | Antineoplastic | 1988 |
| 358 | Imatinib Mesylate | No hit | No hit | Tyrosine kinase inhibitor | Tyrosine kinase inhibitor | | Antineoplastic | 2001 |
| 359 | Imipenem | No hit | No hit | Penicillin-binding protein inhibitor (β-lactam) | β-lactam (PBP inhibitor) | | Antibiotic | 1985 |
| 360 | Imipramine·HCl | No hit | No hit | NE / 5-HT / DA reuptake inhibitor | Serotonin reuptake inhibitor (SSRI/SNRI) | H1 receptor antagonist; Muscarinic receptor antagonist | Antidepressant | 1959 |
| 361 | Imiquimod | No hit | No hit | TLR7 agonist | Innate immune modulator | | Immunomodulant | 1997 |
| 362 | Indapamide | No hit | No hit | Thiazide-like diuretic (NCC inhibitor) | Diuretic | | Antihypertensive | 1983 |
| 363 | Indomethacin | No hit | Toxic | COX inhibitor (NSAID) | COX inhibitor (NSAID) | | NSAID | 1965 |
| 364 | Ipratropium·Br | No hit | No hit | Muscarinic receptor antagonist | Muscarinic receptor antagonist | | Bronchodilator | 1986 |
| 365 | Isoniazid | No hit | No hit | Mycolic acid synthesis inhibitor | Mycobacterial cell wall inhibitor | | Antitubercular | Unknown |
| 366 | Itraconazole | Toxic | Not screened | Ergosterol synthesis inhibitor (azole) | Ergosterol synthesis inhibitor | | Antifungal | 1992 |
| 367 | Ivermectin | Toxic | Not screened | Glutamate-gated $Cl^{-}$ channel agonist | Antiparasitic | | Anthelmintic | 1996 |
| 368 | Ketoconazole | No hit | Toxic | Ergosterol synthesis inhibitor (azole) | Ergosterol synthesis inhibitor | | Antifungal | 1981 |
| 369 | Ketoprofen | Hit | Hit | COX inhibitor (NSAID) | COX inhibitor (NSAID) | | NSAID | 1986 |
| 370 | Ketotifen Fumarate | No hit | Not screened | H1 receptor antagonist (antihistamine); also mast cell stabilizer | H1 receptor antagonist | Leukotriene / arachidonate pathway | Antihistamine | 2006 |
| 371 | L-Ascorbic Acid | No hit | No hit | Antioxidant / enzyme cofactor | Antioxidant | | Vitamin | Unknown |

| | | | | | | | | |
|---|---|---|---|---|---|---|---|---|
| 372 | Lamotrigine | No hit | No hit | Voltage-gated $Na^{+}$ channel blocker; also inhibits voltage-gated calcium channels | Voltage-gated $Na^{+}$ channel blocker | Voltage-gated $Ca^{2+}$ channel (VGCC) blocker | Anticonvulsant | 1994 |
| 373 | Lapatinib Ditosylate | No hit | No hit | Tyrosine kinase inhibitor | Tyrosine kinase inhibitor | | Antineoplastic | 2007 |
| 374 | Latanoprost | No hit | No hit | Prostaglandin FP receptor agonist | Prostaglandin receptor agonist | | Antiglaucoma | 1996 |
| 375 | Leflunomide | No hit | No hit | Dihydroorotate dehydrogenase (DHODH) inhibitor | DHODH inhibitor | | Antirheumatic | 1998 |
| 376 | Letrozole | No hit | No hit | Aromatase inhibitor | Aromatase inhibitor | | Antineoplastic | 1997 |
| 377 | Levetiracetam | No hit | No hit | Synaptic vesicle protein SV2A ligand | Synaptic vesicle protein SV2A ligand | | Anticonvulsant | 1999 |
| 378 | Levofloxacin·HCl | No hit | No hit | DNA gyrase / topoisomerase IV inhibitor | DNA gyrase / topoisomerase IV inhibitor | | Antibiotic | 1996 |
| 379 | Levonorgestrel | No hit | No hit | Progesterone receptor agonist; also estrogen receptor modulator | Sex hormone receptor modulator | | Contraceptive | 1990 |
| 380 | Lidocaine·HCl·H2O | No hit | No hit | Voltage-gated $Na^{+}$ channel blocker | Voltage-gated $Na^{+}$ channel blocker | | Anesthetic / Antiarrhythmic | 1948 |
| 381 | Lincomycin·HCl monohydrate | No hit | No hit | 50S ribosomal subunit inhibitor | Ribosomal subunit inhibitor | | Antibiotic | 1964 |
| 382 | Linezolid | No hit | No hit | 50S ribosomal subunit inhibitor (oxazolidinone, 23S rRNA) | Ribosomal subunit inhibitor | | Antibiotic | 2000 |
| 383 | Lisinopril·2H2O | No hit | No hit | Angiotensin-converting enzyme (ACE) inhibitor | RAAS modulator | | Antihypertensive | 1987 |
| 384 | Lomustine | No hit | No hit | DNA alkylating agent | DNA alkylating agent | | Antineoplastic | 1976 |
| 385 | Loperamide·HCl | No hit | No hit | μ-opioid receptor agonist (peripherally restricted) | Opioid receptor modulator | | Antidiarrheal | 1976 |
| 386 | Loratadine | Toxic | Not screened | Peripheral H1 receptor antagonist; also inhibits | H1 receptor antagonist | | Antihistamine | 2003 |

| | | | | | | | | |
|---|---|---|---|---|---|---|---|---|
| | | | | mast cell mediator release | | | | |
| 387 | Losartan Potassium | No hit | Not confirmed in 2nd round | Angiotensin II receptor antagonist (ARB) | RAAS modulator | | Antihypertensive | 1995 |
| 388 | Lovastatin | Toxic | Not screened | HMG-CoA reductase inhibitor (statin) | HMG-CoA reductase inhibitor (statin) | | Lipid-lowering agent | 1987 |
| 389 | Maprotiline·HCl | Toxic | Not screened | NE / 5-HT / DA reuptake inhibitor | Serotonin reuptake inhibitor (SSRI/SNRI) | | Antidepressant | 1980 |
| 390 | Mebendazole | Toxic | Not screened | Helminth tubulin inhibitor | Antiparasitic | | Anthelmintic | 1974 |
| 391 | Medroxyprogesterone Acetate | No hit | No hit | Progesterone receptor agonist | Sex hormone receptor modulator | | Contraceptive | 1959 |
| 392 | Mefenamic Acid | Toxic | Not screened | COX inhibitor (NSAID) | COX inhibitor (NSAID) | | NSAID | 1967 |
| 393 | Megestrol Acetate | No hit | Not screened | Progesterone receptor agonist | Sex hormone receptor modulator | | Contraceptive / Hormonal | 1971 |
| 394 | Meloxicam | Not confirmed in 2nd round | No hit | COX inhibitor (NSAID) | COX inhibitor (NSAID) | | NSAID | 2000 |
| 395 | Melphalan | No hit | Not confirmed in 2nd round | DNA alkylating agent | DNA alkylating agent | | Antineoplastic | Unknown |
| 396 | Memantine·HCl | No hit | Toxic | N-methyl-D-aspartate (NMDA) receptor antagonist | NMDA receptor antagonist | | Antiparkinsonian | 2003 |
| 397 | Meropenem | No hit | No hit | Penicillin-binding protein inhibitor (β-lactam) | β-lactam (PBP inhibitor) | | Antibiotic | 1996 |
| 398 | Mesalamine (5-Aminosalicylic Acid) | No hit | No hit | COX inhibitor (NSAID) | COX inhibitor (NSAID) | | Anti-inflammatory | 1987 |

| | | | | | | | | |
|---|---|---|---|---|---|---|---|---|
| 399 | Metformin·HCl | No hit | No hit | Mitochondrial complex I inhibitor / AMPK activator | Mitochondrial complex I inhibitor | | Antidiabetic | 1995 |
| 400 | Methimazole | No hit | Not confirmed in 2nd round | Thyroid peroxidase inhibitor | Thyroid modulator | | Antithyroid | 1950 |
| 401 | Methyldopa Sesquihydrate (L-A-Methyl-Dopa Sesquihyrate) | No hit | Hit | α-adrenoceptor agonist (via α-methylnorepinephrine); false neurotransmitter (catecholamine depletion) | Adrenergic receptor agonist | Dopaminergic inhibitor | Antihypertensive | 1962 |
| 402 | Methylprednisolone | No hit | No hit | Glucocorticoid receptor agonist | Glucocorticoid receptor agonist | | Anti-inflammatory | 1957 |
| 403 | Metoclopramide·HCl | No hit | No hit | Dopamine D2 receptor antagonist; 5-HT4 receptor agonist; 5-HT3 antagonist | Dopaminergic inhibitor | 5-HT receptor modulator | Antiemetic | Unknown |
| 404 | Metoprolol Tartrate | No hit | Hit | β1-adrenoceptor antagonist (selective) | Adrenergic receptor antagonist | | Antihypertensive | 1978 |
| 405 | Metronidazole | No hit | No hit | Nitroimidazole (DNA damage) | Antiprotozoal | | Antiprotozoal | 1963 |
| 406 | Mifepristone | No hit | Not screened | Progesterone receptor antagonist; also glucocorticoid receptor antagonist | Sex hormone receptor modulator | Glucocorticoid receptor antagonist | Contraceptive | 2000 |
| 407 | Miglustat (N-Butyldeoxynojirimycin·HCl) | No hit | Not screened | Glucosylceramide synthase inhibitor | Glycosphingolipid synthesis inhibitor | | Antiviral | Unknown |
| 408 | Milrinone | No hit | No hit | Phosphodiesterase 3 (PDE3) inhibitor | Phosphodiesterase (PDE) inhibitor | | Vasodilator | Unknown |
| 409 | Minocycline | No hit | Not confirmed in 2nd round | 30S ribosomal subunit inhibitor | Ribosomal subunit inhibitor | | Antibiotic | 1970;s |
| 410 | Minoxidil | No hit | No hit | ATP-sensitive $K^+$ channel opener | ATP-sensitive $K^+$ channel modulator | | Vasodilator | 1979 |

| | | | | | | | | |
|---|---|---|---|---|---|---|---|---|
| 411 | Misoprostol | No hit | No hit | Prostaglandin EP receptor agonist | Prostaglandin receptor agonist | | Antiulcer | 1988 |
| 412 | Mitomycin C | No hit | No hit | DNA alkylating agent | DNA alkylating agent | | Antineoplastic | 1981 |
| 413 | Mitoxantrone·2HCl | Not screened | Toxic | Topoisomerase II inhibitor; DNA intercalator | Topoisomerase inhibitor | | Antineoplastic | 1987 |
| 414 | Montelukast·Na | Toxic | Not screened | CysLT1 receptor antagonist | Leukotriene / arachidonate pathway | | Antiasthmatic | 1998 |
| 415 | Moxifloxacin·HCl | No hit | Initial hit, not retested | DNA gyrase / topoisomerase IV inhibitor | DNA gyrase / topoisomerase IV inhibitor | | Antibiotic | 1999 |
| 416 | Mycophenolate Mofetil | No hit | No hit | Inosine monophosphate dehydrogenase (IMPDH) inhibitor | IMPDH inhibitor | | Antineoplastic / Immunosuppressant | 1995 |
| 417 | Mycophenolic Acid | No hit | No hit | Inosine monophosphate dehydrogenase (IMPDH) inhibitor | IMPDH inhibitor | | Antibiotic / Antineoplastic | 2004 |
| 418 | Nabumetone | No hit | No hit | COX inhibitor (NSAID) | COX inhibitor (NSAID) | | NSAID | 1991 |
| 419 | Nalbuphine·HCl | No hit | Not screened | Opioid receptor agonist/antagonist | Opioid receptor modulator | | Analgesic | Unknown |
| 420 | Naloxone·HCl | No hit | No hit | Opioid receptor agonist/antagonist | Opioid receptor modulator | | Opioid antagonist | 1971 |
| 421 | Naltrexone·HCl | No hit | No hit | Opioid receptor agonist/antagonist | Opioid receptor modulator | | Opioid antagonist | 1984 |
| 422 | Naphazoline·HCl | No hit | Not confirmed in 2nd round | α1-adrenoceptor agonist | Adrenergic receptor agonist | | Vasoconstrictor | Unknown |
| 423 | Naproxen | No hit | Toxic | COX inhibitor (NSAID) | COX inhibitor (NSAID) | | NSAID | 1976 |
| 424 | Nateglinide | No hit | No hit | ATP-sensitive $K^+$ channel blocker (insulin secretagogue) | ATP-sensitive $K^+$ channel modulator | | Antidiabetic | 2000 |

| | | | | | | | | |
|---|---|---|---|---|---|---|---|---|
| 425 | Nefazodone·HCl | Toxic | Not screened | 5-HT2 receptor antagonist; also serotonin uptake inhibitor | 5-HT receptor modulator | Serotonin reuptake inhibitor (SSRI/SNRI) | Antidepressant | 1994 |
| 426 | Nelfinavir Mesylate | No hit | Not screened | HIV protease inhibitor | HIV protease inhibitor | | Antiviral | 1997 |
| 427 | Neomycin trisulfate | No hit | No hit | 30S ribosomal subunit inhibitor | Ribosomal subunit inhibitor | | Antibiotic | Unknown |
| 428 | Nicardipine·HCl | Toxic | Not screened | Voltage-gated $Ca^{2+}$ channel (VGCC) blocker | Voltage-gated $Ca^{2+}$ channel (VGCC) blocker | | Antihypertensive | 1988 |
| 429 | Nifedipine | Not confirmed in 2nd round | No hit | Voltage-gated $Ca^{2+}$ channel (VGCC) blocker | Voltage-gated $Ca^{2+}$ channel (VGCC) blocker | | Antihypertensive | 1981 |
| 430 | Nimodipine | Toxic | Not screened | Voltage-gated $Ca^{2+}$ channel (VGCC) blocker | Voltage-gated $Ca^{2+}$ channel (VGCC) blocker | | Antihypertensive | 1988 |
| 431 | Nisoldipine | Toxic | Not screened | Voltage-gated $Ca^{2+}$ channel (VGCC) blocker | Voltage-gated $Ca^{2+}$ channel (VGCC) blocker | | Antihypertensive | 1995 |
| 432 | Norepinephrine Bitartrate Monohydrate | No hit | No hit | Adrenergic receptor agonist (α and β) | Adrenergic receptor agonist | | Vasoconstrictor | Unknown |
| 433 | Norethindrone | No hit | No hit | Progesterone receptor agonist | Sex hormone receptor modulator | | Contraceptive | 1973 |
| 434 | Norfloxacin | No hit | No hit | DNA gyrase / topoisomerase IV inhibitor | DNA gyrase / topoisomerase IV inhibitor | | Antibiotic | 1986 |
| 435 | Nystatin | Not confirmed in 2nd round | No hit | Ergosterol-binding polyene antifungal | Ergosterol-binding antifungal | | Antifungal | Unknown |
| 436 | Ofloxacin | Not confirmed in 2nd round | No hit | DNA gyrase / topoisomerase IV inhibitor | DNA gyrase / topoisomerase IV inhibitor | | Antibiotic | 1990 |

| | | | | | | | | |
|---|---|---|---|---|---|---|---|---|
| 437 | Olanzapine | Not confirmed in 2nd round | No hit | Dopamine D2 receptor antagonist; 5-HT2A receptor antagonist | Dopaminergic inhibitor | 5-HT receptor modulator; H1 receptor antagonist; Muscarinic receptor antagonist | Antipsychotic | 1996 |
| 438 | Olmesartan | No hit | No hit | Angiotensin II receptor antagonist (ARB) | RAAS modulator | | Antihypertensive | Unknown |
| 439 | Olopatadine | No hit | No hit | H1 receptor antagonist (antihistamine) | H1 receptor antagonist | Muscarinic receptor antagonist | Antihistamine | Unknown |
| 440 | Omeprazole | No hit | No hit | $H^+/K^+$-ATPase inhibitor (PPI) | $H^+/K^+$-ATPase inhibitor (PPI) | | Antiulcer | 1989 |
| 441 | Ondansetron | Not confirmed in 2nd round | No hit | 5-HT3 receptor antagonist | 5-HT receptor modulator | | Antiemetic | Unknown |
| 442 | Orphenadrine Citrate | No hit | No hit | H1 receptor antagonist (antihistamine); N-methyl-D-aspartate (NMDA) receptor antagonist | H1 receptor antagonist | Muscarinic receptor antagonist; NMDA receptor antagonist | Muscle relaxant | 1959 |
| 443 | Oseltamivir Phosphate | No hit | No hit | Viral neuraminidase inhibitor | Other antiviral | | Antiviral | 1999 |
| 444 | Oxacillin sodium salt monohydrate | No hit | No hit | Penicillin-binding protein inhibitor (β-lactam) | β-lactam (PBP inhibitor) | | Antibiotic | Unknown |
| 445 | Oxaliplatin | No hit | No hit | DNA crosslinker (platinum) | DNA crosslinker | | Antineoplastic | 2002 |
| 446 | Oxcarbazepine | No hit | No hit | Voltage-gated $Na^+$ channel blocker | Voltage-gated $Na^+$ channel blocker | | Anticonvulsant | 2000 |
| 447 | Oxiconazole Nitrate | Toxic | Not screened | Ergosterol synthesis inhibitor (azole) | Ergosterol synthesis inhibitor | | Antifungal | 1988 |
| 448 | Paclitaxel (Taxol) | Toxic | Not screened | Microtubule stabiliser | Microtubule modulator | | Antineoplastic | 1992 |
| 449 | Pamidronate Disodium Pentahydrate (Pamidronic Acid) | No hit | No hit | Farnesyl pyrophosphate synthase inhibitor (bisphosphonate) | Bisphosphonate | | Bone resorption inhibitor | Unknown |

| | | | | | | | | |
|---|---|---|---|---|---|---|---|---|
| 450 | Pancuronium·2Br | No hit | No hit | Nicotinic receptor antagonist (NMJ) | Cholinergic antagonist | | Neuromuscular blocking agent | 1972 |
| 451 | Pantoprazole | No hit | No hit | $H^+/K^+$-ATPase inhibitor (PPI) | $H^+/K^+$-ATPase inhibitor (PPI) | | Antiulcer | Unknown |
| 452 | Paroxetine·HCl | Toxic | No hit | Serotonin reuptake inhibitor (SSRI/SNRI) | Serotonin reuptake inhibitor (SSRI/SNRI) | | Antidepressant | 1992 |
| 453 | Penciclovir | No hit | No hit | Viral DNA polymerase inhibitor | Viral DNA polymerase inhibitor | | Antiviral | 2000's |
| 454 | Penicillin V Potassium | Not confirmed in 2nd round | No hit | Penicillin-binding protein inhibitor (β-lactam) | β-lactam (PBP inhibitor) | | Antibiotic | Unknown |
| 455 | Pentoxifylline | Hit | No hit | Phosphodiesterase (PDE) inhibitor; non-selective adenosine receptor antagonist | Phosphodiesterase (PDE) inhibitor | Adenosine receptor modulator | Hematologic agent | 1984 |
| 456 | Phenoxybenzamine·HCl | No hit | Toxic | α-adrenoceptor antagonist (irreversible) | Adrenergic receptor antagonist | | Antihypertensive | 1953 |
| 457 | Phentolamine·HCl | No hit | No hit | α1/α2-adrenoceptor antagonist | Adrenergic receptor antagonist | | Vasodilator | Unknown |
| 458 | Phenytoin | No hit | No hit | Voltage-gated $Na^+$ channel blocker | Voltage-gated $Na^+$ channel blocker | | Anticonvulsant | 1953 |
| 459 | Pilocarpine·HCl | No hit | No hit | Muscarinic receptor agonist | Muscarinic receptor agonist | | Parasympathomimetic | Unknown |
| 460 | Pimozide | Toxic | Not screened | Dopamine D2 receptor antagonist | 5-HT receptor modulator | Dopaminergic inhibitor | Antipsychotic | 1984 |
| 461 | Pindolol | No hit | No hit | β-adrenoceptor antagonist; also 5-HT1A partial agonist | Adrenergic receptor antagonist | 5-HT receptor modulator | Antihypertensive | 1982 |
| 462 | Pioglitazone·HCl | No hit | No hit | PPARγ agonist | PPAR agonist | | Antidiabetic | 1999 |
| 463 | Piperacillin | Hit | No hit | Penicillin-binding protein inhibitor (β-lactam) | β-lactam (PBP inhibitor) | | Antibiotic | Unknown |
| 464 | Piroxicam | No hit | No hit | COX inhibitor (NSAID) | COX inhibitor (NSAID) | | NSAID | 1982 |

| | | | | | | | | |
|---|---|---|---|---|---|---|---|---|
| 465 | Pramipexole Dihydrochloride Monohydrate | No hit | No hit | Dopamine receptor agonist | Dopaminergic potentiator | | Antiparkinsonian | 2006 |
| 466 | Praziquantel | No hit | Not screened | Schistosome TRPM_PZQ channel agonist | Antiparasitic | | Anthelmintic | 1982 |
| 467 | Prazosin·HCl | No hit | No hit | α1-adrenoceptor antagonist | Adrenergic receptor antagonist | | Antihypertensive | 1976 |
| 468 | Prednisolone | No hit | Not screened | Glucocorticoid receptor agonist | Glucocorticoid receptor agonist | | Anti-inflammatory | 1955 |
| 469 | Prednisone | No hit | Not screened | Glucocorticoid receptor agonist | Glucocorticoid receptor agonist | | Anti-inflammatory | 1955 |
| 470 | Primaquine diphosphate | Hit | Not screened | Reactive oxygen species generator (haem-activated) | Antimalarial | | Antimalarial | 1952 |
| 471 | Procainamide·HCl | No hit | No hit | Voltage-gated $Na^+$ channel blocker | Voltage-gated $Na^+$ channel blocker | | Antiarrhythmic | 1950 |
| 472 | Procarbazine·HCl | Not confirmed in 2nd round | Not confirmed in 2nd round | DNA alkylating agent | DNA alkylating agent | | Antineoplastic | 1969 |
| 473 | Progesterone | No hit | Not screened | Progesterone receptor agonist; also estrogen receptor modulator | Sex hormone receptor modulator | | Contraceptive | Unknown |
| 474 | Promethazine·HCl | Toxic | No hit | H1 receptor antagonist (antihistamine) | H1 receptor antagonist | Dopaminergic inhibitor; Muscarinic receptor antagonist | Antihistamine | 1951 |
| 475 | Propafenone·HCl | No hit | No hit | Voltage-gated $Na^+$ channel blocker | Voltage-gated $Na^+$ channel blocker | Adrenergic receptor antagonist | Antiarrhythmic | 1989 |
| 476 | Propofol | No hit | No hit | GABAA receptor modulator; also sodium channel blocker | GABAergic modulator | | Anesthetic | 1989 |
| 477 | Propranolol·HCl | No hit | Not screened | β-adrenoceptor antagonist (non-selective) | Adrenergic receptor antagonist | | Antihypertensive | 1967 |

| | | | | | | | | |
|---|---|---|---|---|---|---|---|---|
| 478 | Quetiapine Fumarate | Not confirmed in 2nd round | No hit | Dopamine D2 receptor antagonist; 5-HT2A receptor antagonist | Dopaminergic inhibitor | 5-HT receptor modulator; Adrenergic receptor antagonist; H1 receptor antagonist; Muscarinic receptor antagonist | Antipsychotic | 1997 |
| 479 | Quinapril·HCl | No hit | Not screened | Angiotensin-converting enzyme (ACE) inhibitor | RAAS modulator | | Antihypertensive | 1991 |
| 480 | Quinine·HCl·2H2O | No hit | Not confirmed in 2nd round | Heme polymerisation inhibitor | Antimalarial | | Antimalarial | Unknown |
| 481 | Raloxifene·HCl | No hit | Toxic | Estrogen receptor modulator (SERM) | Sex hormone receptor modulator | | Bone resorption inhibitor | 1997 |
| 482 | Ramipril | No hit | Not screened | Angiotensin-converting enzyme (ACE) inhibitor | RAAS modulator | | Antihypertensive | 1991 |
| 483 | Ranitidine·HCl | Initial hit, not retested | No hit | H2 receptor antagonist | H2 receptor antagonist | | Antiulcer | 1983 |
| 484 | Ranolazine·2HCl | Not confirmed in 2nd round | No hit | Voltage-gated $Na^+$ channel blocker (late sodium current) | Voltage-gated $Na^+$ channel blocker | | Antianginal | 2006 |
| 485 | Ribavirin | No hit | Not screened | IMPDH inhibitor; viral RNA polymerase inhibitor | IMPDH inhibitor | | Antiviral | 1985 |
| 486 | Rifampin (Rifampicin) | No hit | No hit | Bacterial DNA-dependent RNA polymerase inhibitor | RNA polymerase inhibitor | | Antibiotic | 1971 |
| 487 | Riluzole·HCl | Not confirmed in 2nd round | Toxic | Voltage-gated $Na^+$ channel blocker; also inhibits glutamate release | Voltage-gated $Na^+$ channel blocker | | Neuroprotective | 1995 |
| 488 | Rimantadine·HCl | No hit | Not screened | Viral M2 ion channel inhibitor | Viral ion channel inhibitor | | Antiviral | 1993 |

| | | | | | | | | |
|---|---|---|---|---|---|---|---|---|
| 489 | Risedonic Acid | No hit | No hit | Farnesyl pyrophosphate synthase inhibitor (bisphosphonate) | Bisphosphonate | | Bone resorption inhibitor | Unknown |
| 490 | Risperidone | No hit | No hit | Dopamine D2 receptor antagonist; 5-HT2A receptor antagonist | Dopaminergic inhibitor | 5-HT receptor modulator | Antipsychotic | 1993 |
| 491 | Rivastigmine Tartrate | No hit | No hit | Acetylcholinesterase inhibitor | Cholinergic agonist | | Parasympathomimetic | 2000 |
| 492 | Rocuronium Bromide | No hit | No hit | Nicotinic receptor antagonist (NMJ) | Cholinergic antagonist | | Muscle relaxant | 1994 |
| 493 | Rosiglitazone | No hit | No hit | PPARγ agonist | PPAR agonist | | Antidiabetic | Unknown |
| 494 | S(-)-Carbidopa . monohydrate | No hit | Hit | Aromatic L-amino acid decarboxylase (AADC) inhibitor | Dopaminergic inhibitor | | Antiparkinsonian | 1975 |
| 495 | Salbutamol Hemisulfate | No hit | No hit | β2-adrenoceptor agonist (selective) | Adrenergic receptor agonist | | Bronchodilator | 1981 |
| 496 | Salmeterol | No hit | No hit | β2-adrenoceptor agonist (long-acting) | Adrenergic receptor agonist | | Bronchodilator | 2000's |
| 497 | Scopolamine·HBr | Hit | No hit | Muscarinic receptor antagonist | Muscarinic receptor antagonist | | Antiemetic / Anticholinergic | 1979 |
| 498 | Sertaconazole | Toxic | Not screened | Ergosterol synthesis inhibitor (azole) | Ergosterol synthesis inhibitor | | Antifungal | Unknown |
| 499 | Sildenafil Citrate | No hit | No hit | Phosphodiesterase 5 (PDE5) inhibitor | Phosphodiesterase (PDE) inhibitor | | Vasodilator | 1998 |
| 500 | Simvastatin | Toxic | Not screened | HMG-CoA reductase inhibitor (statin) | HMG-CoA reductase inhibitor (statin) | | Lipid-lowering agent | 1991 |
| 501 | Sirolimus (Rapamycin) | No hit | No hit | mTOR inhibitor | mTOR inhibitor | | Immunosuppressant | 1999 |
| 502 | Sodium Phenylbutyrate | Not screened | Not confirmed in 2nd round | Ammonia scavenger (phenylacetylglutamine conjugation); HDAC inhibitor | Urea cycle modulator | | Antineoplastic | 1996 |
| 503 | Sotalol·HCl | No hit | No hit | β-adrenoceptor antagonist; also blocks IKr potassium channels | Adrenergic receptor antagonist | Voltage-gated $K^+$ channel blocker | Antiarrhythmic | 1992 |

| | | | | | | | | |
|---|---|---|---|---|---|---|---|---|
| 504 | Spectinomycin·2HCl Pentahydrate | Not screened | No hit | 30S ribosomal subunit inhibitor | Ribosomal subunit inhibitor | | Antibiotic | 1971 |
| 505 | Spironolactone | No hit | Not screened | Mineralocorticoid receptor (MR) antagonist | Diuretic | | Diuretic | 1960 |
| 506 | Streptomycin Sulfate | No hit | No hit | 30S ribosomal subunit inhibitor | Ribosomal subunit inhibitor | | Antibiotic | 1946 |
| 507 | Succinylcholine Chloride·2H2O | No hit | Not screened | Nicotinic receptor agonist (depolarising) | Cholinergic agonist | | Neuromuscular blocking agent | Unknown |
| 508 | Sulfadiazine | Not confirmed in 2nd round | Not confirmed in 2nd round | Bacterial dihydropteroate synthetase (DHPS) inhibitor | Folate pathway inhibitor | | Antibacterial | 1941 |
| 509 | Sulfasalazine | No hit | Not screened | Prodrug releasing 5-aminosalicylic acid and sulfapyridine | COX inhibitor (NSAID) | | Antirheumatic | 1950 |
| 510 | Sulindac | No hit | No hit | COX inhibitor (NSAID) | COX inhibitor (NSAID) | | NSAID | 1978 |
| 511 | Sumatriptan Succinate | Hit | No hit | 5-HT1B/1D receptor agonist | 5-HT receptor modulator | | Antimigraine | 1992 |
| 512 | Tacrine·HCl | No hit | No hit | Acetylcholinesterase inhibitor | Cholinergic agonist | | Parasympathomimetic | 1993 |
| 513 | Tamoxifen Citrate | Toxic | Not screened | Estrogen receptor modulator (SERM) | Sex hormone receptor modulator | | Antineoplastic | 1977 |
| 514 | Tamsulosin·HCl | No hit | Not screened | α1A/α1D-adrenoceptor antagonist | Adrenergic receptor antagonist | | Antineoplastic? (BPH treatment) | 1997 |
| 515 | Telmisartan | No hit | Not screened | Angiotensin II receptor antagonist (ARB) | RAAS modulator | | Antihypertensive | 2000 |
| 516 | Temozolomide | No hit | Not screened | DNA alkylating agent | DNA alkylating agent | | Antineoplastic | 1999 |
| 517 | Terazosin·HCl | No hit | Not screened | α1-adrenoceptor antagonist | Adrenergic receptor antagonist | | Antihypertensive | 1998 |
| 518 | Terbinafine·HCl | Toxic | Not screened | Squalene epoxidase inhibitor | Ergosterol synthesis inhibitor | | Antifungal | 1992 |
| 519 | Tetracycline | No hit | Not screened | 30S ribosomal subunit inhibitor | Ribosomal subunit inhibitor | | Antibiotic | 1950's |

| | | | | | | | | |
|---|---|---|---|---|---|---|---|---|
| 520 | Thalidomide | No hit | No hit | Cereblon (CRBN) E3 ubiquitin ligase modulator | Innate immune modulator | | Immunomodulant | 1998 |
| 521 | Thioridazine·HCl | Toxic | Not screened | Dopamine receptor antagonist | Dopaminergic inhibitor | H1 receptor antagonist; Muscarinic receptor antagonist | Antipsychotic | 1962 |
| 522 | Ticlopidine·HCl | No hit | No hit | P2Y12 receptor antagonist | Antiplatelet / Anticoagulant | | Antiplatelet | 1991 |
| 523 | Tinidazole | No hit | Not screened | Nitroimidazole (DNA damage) | Antiprotozoal | | Antiprotozoal | 2004 |
| 524 | Tiotropium Bromide | Hit | Hit | Muscarinic receptor antagonist | Muscarinic receptor antagonist | | Bronchodilator | 2004 |
| 525 | Tizanidine·HCl | No hit | No hit | α2-adrenoceptor agonist | Adrenergic receptor agonist | | Muscle relaxant | 1996 |
| 526 | Tobramycin | No hit | Not screened | 30S ribosomal subunit inhibitor | Ribosomal subunit inhibitor | | Antibiotic | 1975 |
| 527 | Tolazamide | No hit | No hit | ATP-sensitive $K^+$ channel blocker (insulin secretagogue) | ATP-sensitive $K^+$ channel modulator | | Antidiabetic | 1966 |
| 528 | Tolbutamide | No hit | No hit | ATP-sensitive $K^+$ channel blocker (insulin secretagogue) | ATP-sensitive $K^+$ channel modulator | | Antidiabetic | 1961 |
| 529 | Tolcapone | No hit | Hit | Catechol-O-methyltransferase (COMT) inhibitor | Dopaminergic potentiator | | Antiparkinsonian | 1998 |
| 530 | Tolmetin sodium dihydrate | No hit | Not screened | COX inhibitor (NSAID) | COX inhibitor (NSAID) | | NSAID | 1976 |
| 531 | Topotecan·HCl | No hit | Not screened | Topoisomerase I inhibitor | Topoisomerase inhibitor | | Antineoplastic | 1996 |
| 532 | Toremifene Base | Toxic | Not screened | Estrogen receptor modulator (SERM) | Sex hormone receptor modulator | | Antineoplastic | Unknown |
| 533 | Tramadol·HCl | No hit | Not screened | μ-opioid receptor agonist; also inhibits serotonin/norepinephrine reuptake | Opioid receptor modulator | Serotonin reuptake inhibitor (SSRI/SNRI) | Analgesic | 1995 |

| | | | | | | | | |
|---|---|---|---|---|---|---|---|---|
| 534 | Tranexamic Acid | No hit | No hit | Antifibrinolytic (plasminogen inhibitor) | Antifibrinolytic | | Antifibrinolytic | 1986 |
| 535 | Tranylcypromine Hemisulfate | No hit | No hit | MAO inhibitor | MAO inhibitor | | Antidepressant | 1961 |
| 536 | Trifluoperazine· HCl | Toxic | Not screened | Dopamine receptor antagonist | Dopaminergic inhibitor | | Antipsychotic | 1959 |
| 537 | Trimethoprim | Hit | Hit | Bacterial dihydrofolate reductase (DHFR) inhibitor | Folate pathway inhibitor | | Antibiotic | Unknown |
| 538 | Triptorelin Acetate | No hit | No hit | GnRH agonist | Sex hormone receptor modulator | | Hormonal therapy | Unknown |
| 539 | Tropicamide | No hit | No hit | Muscarinic receptor antagonist | Muscarinic receptor antagonist | | Mydriatic | 1960 |
| 540 | Valacyclovir·HCl | No hit | Not screened | Viral DNA polymerase inhibitor | Viral DNA polymerase inhibitor | | Antiviral | 1995 |
| 541 | Valproate·Na | No hit | No hit | Voltage-gated $Na^+$ channel blocker; also inhibits GABA transaminase and HDAC | Voltage-gated $Na^+$ channel blocker | HDAC inhibitor | Anticonvulsant | 1996 |
| 542 | Vardenafil | No hit | Initial hit, not retested | Phosphodiesterase 5 (PDE5) inhibitor | Phosphodiesterase (PDE) inhibitor | | Vasodilator | Unknown |
| 543 | Vecuronium Bromide | No hit | Not screened | Nicotinic receptor antagonist (NMJ) | Cholinergic antagonist | | Muscle relaxant | 1984 |
| 544 | Venlafaxine·HCl | No hit | Not screened | Serotonin reuptake inhibitor (SSRI/SNRI) | Serotonin reuptake inhibitor (SSRI/SNRI) | | Antidepressant | 1993 |
| 545 | Verapamil·HCl | No hit | No hit | Voltage-gated $Ca^{2+}$ channel (VGCC) blocker | Voltage-gated $Ca^{2+}$ channel (VGCC) blocker | | Antihypertensive | 1981 |
| 546 | Vinblastine Sulfate | No hit | No hit | Microtubule destabiliser | Microtubule modulator | | Antineoplastic | 1965 |
| 547 | Vincristine Sulfate | No hit | No hit | Microtubule destabiliser | Microtubule modulator | | Antineoplastic | 1963 |
| 548 | Vinorelbine | No hit | No hit | Microtubule destabiliser | Microtubule modulator | | Antineoplastic | 1994 |

| | | | | | | | | |
|---|---|---|---|---|---|---|---|---|
| 549 | Vorinostat | No hit | No hit | Histone deacetylase (HDAC) inhibitor | HDAC inhibitor | | Antineoplastic | 2006 |
| 550 | Zafirlukast | No hit | No hit | CysLT1 receptor antagonist | Leukotriene / arachidonate pathway | | Antiasthmatic | 1996 |
| 551 | Zalcitabine (2',3'-Dideoxycytidine) | Hit | No hit | Nucleoside Reverse Transcriptase Inhibitor (NRTI) | Reverse transcriptase inhibitor | | Antiviral, Anti-HIV Agent | 1992 |
| 552 | Zidovudine (3'-Azido-3'-Deoxythymidine) | No hit | No hit | Nucleoside Reverse Transcriptase Inhibitor (NRTI) | Reverse transcriptase inhibitor | | Antiviral | 1987 |
| 553 | Zileuton | No hit | No hit | 5-Lipoxygenase inhibitor | Leukotriene / arachidonate pathway | | Antiasthmatic | 1996 |
| 554 | Zoledronic Acid Monohydrate | Not confirmed in 2nd round | No hit | Farnesyl pyrophosphate synthase inhibitor (bisphosphonate) | Bisphosphonate | | Bone resorption inhibitor | Unknown |
| 555 | Zolmitriptan | No hit | No hit | 5-HT1B/1D receptor agonist | 5-HT receptor modulator | | Antimigrane | 1997 |
| 556 | Zonisamide | No hit | No hit | Voltage-gated $Na^+$ channel blocker; also reduces T-type calcium currents | Voltage-gated $Na^+$ channel blocker | | Anticonvulsant | Unknown |

**Supplementary Data 2 - Mechanism-of-action enrichment analysis.** Enrichment of mechanism-of-action (MOA) classes among screen hit compounds relative to the full library, assessed using a two-sided Fisher's exact test. Secondary MOA classes were assigned to compounds when said activity was independently responsible for a documented clinical or adverse effect or was a recognized mechanistic contributor to the approved pharmacological action. The total number of compounds in the library and among the hits, the odds ratio, and the nominal p-value are shown for each class, together with the identity of the hit compounds. Classes containing fewer than three compounds are omitted. The analysis was repeated using the primary MOA class alone as a sensitivity control.

| MOA class | Primary + secondary MOA classes | | | | | | | Primary MOA class only | | | | Hit compounds |
|---|---|---|---|---|---|---|---|---|---|---|---|---|
| | N (library) | % of library | N (hits) | % of hits | N (non hits) | Odds ratio | p-value (Fisher) | N (library) | N (hits) | Odds ratio | p-value (Fisher) | |
| Muscarinic receptor antagonist | 36 | 6.47 | 5 | 21.7 | 31 | 4.5 | 0.0123 | 12 | 4 | 13.82 | 0.0009 | Dicyclomine·HCl, Disopyramide, Glycopyrrolate Iodide, Scopolamine·HBr, Tiotropium Bromide |
| Phosphodiesterase (PDE) inhibitor | 11 | 1.98 | 2 | 8.7 | 9 | 5.54 | 0.0718 | 9 | 2 | 7.16 | 0.0494 | Amrinone, Pentoxifylline |
| Dopaminergic inhibitor | 18 | 3.24 | 2 | 8.7 | 16 | 3.08 | 0.1677 | 14 | 1 | 1.82 | 0.4504 | Methyldopa Sesquihydrate (L-A-Methyl-Dopa Sesquihyrate), S(-)-Carbidopa . monohydrate |
| COX inhibitor (NSAID) | 22 | 3.96 | 2 | 8.7 | 20 | 2.44 | 0.2294 | 22 | 2 | 2.44 | 0.2294 | Etodolac, Ketoprofen |
| β-lactam (PBP inhibitor) | 30 | 5.4 | 2 | 8.7 | 28 | 1.72 | 0.3559 | 30 | 2 | 1.72 | 0.3559 | Ceftizoxim·Na, Piperacillin |
| Folate pathway inhibitor | 4 | 0.72 | 1 | 4.3 | 3 | 8.03 | 0.1559 | 4 | 1 | 8.03 | 0.1559 | Trimethoprim |
| Mycobacterial cell wall inhibitor | 4 | 0.72 | 1 | 4.3 | 3 | 8.03 | 0.1559 | 4 | 1 | 8.03 | 0.1559 | Ethambutol Dihydrochloride |
| Antimalarial | 5 | 0.9 | 1 | 4.3 | 4 | 6.01 | 0.191 | 5 | 1 | 6.01 | 0.191 | Primaquine diphosphate |
| Adenosine receptor modulator | 7 | 1.26 | 1 | 4.3 | 6 | 3.99 | 0.2572 | 2 | 0 | 0 | 1 | Pentoxifylline |
| Reverse transcriptase inhibitor | 8 | 1.44 | 1 | 4.3 | 7 | 3.42 | 0.2883 | 8 | 1 | 3.42 | 0.2883 | Zalcitabine (2',3'-Dideoxycytidine) |

| | | | | | | | | | | | |
|---|---|---|---|---|---|---|---|---|---|---|---|
| Ergosterol synthesis inhibitor | 10 | 1.8 | 1 | 4.3 | 9 | 2.65 | 0.3469 | 10 | 1 | 2.65 | 0.3469 | Fluconazole |
| Dopamine potentiator | 11 | 1.98 | 1 | 4.3 | 10 | 2.38 | 0.3744 | 7 | 1 | 3.99 | 0.2572 | Tolcapone |
| DNA alkylating agent | 14 | 2.52 | 1 | 4.3 | 13 | 1.82 | 0.4504 | 14 | 1 | 1.82 | 0.4504 | Bendamustine·HCl |
| Voltage-gated $Na^+$ channel blocker | 15 | 2.7 | 1 | 4.3 | 14 | 1.69 | 0.4738 | 15 | 1 | 1.69 | 0.4738 | Disopyramide |
| Adrenergic receptor agonist | 18 | 3.24 | 1 | 4.3 | 17 | 1.38 | 0.5382 | 16 | 1 | 1.57 | 0.4961 | Methyldopa Sesquihydrate (L-A-Methyl-Dopa Sesquihyrate) |
| 5-HT receptor modulator | 27 | 4.86 | 1 | 4.3 | 26 | 0.89 | 1 | 14 | 1 | 1.82 | 0.4504 | Sumatriptan Succinate |
| Adrenergic receptor antagonist | 28 | 5.04 | 1 | 4.3 | 27 | 0.85 | 1 | 18 | 1 | 1.38 | 0.5382 | Metoprolol Tartrate |
| H1 receptor antagonist | 31 | 5.58 | 1 | 4.3 | 30 | 0.76 | 1 | 18 | 1 | 1.38 | 0.5382 | Fexofenadine·HCl |
| Sex hormone receptor modulator | 27 | 4.86 | 0 | 0 | 27 | 0 | 0.6194 | 26 | 0 | 0 | 0.6176 | |
| Glucocorticoid receptor agonist | 24 | 4.32 | 0 | 0 | 24 | 0 | 0.6159 | 23 | 0 | 0 | 0.6161 | |
| Ribosomal subunit inhibitor | 19 | 3.42 | 0 | 0 | 19 | 0 | 1 | 19 | 0 | 0 | 1 | |
| Serotonin reuptake inhibitor (SSRI/SNRI) | 17 | 3.06 | 0 | 0 | 17 | 0 | 1 | 15 | 0 | 0 | 1 | |
| RAAS modulator | 13 | 2.34 | 0 | 0 | 13 | 0 | 1 | 13 | 0 | 0 | 1 | |
| Diuretic | 12 | 2.16 | 0 | 0 | 12 | 0 | 1 | 12 | 0 | 0 | 1 | |
| Voltage-gated $Ca^{2+}$ channel (VGCC) blocker | 11 | 1.98 | 0 | 0 | 11 | 0 | 1 | 10 | 0 | 0 | 1 | |
| ATP-sensitive $K^+$ channel modulator | 10 | 1.8 | 0 | 0 | 10 | 0 | 1 | 10 | 0 | 0 | 1 | |
| Antimetabolite | 9 | 1.62 | 0 | 0 | 9 | 0 | 1 | 9 | 0 | 0 | 1 | |
| Topoisomerase inhibitor | 7 | 1.26 | 0 | 0 | 7 | 0 | 1 | 7 | 0 | 0 | 1 | |
| Viral DNA polymerase inhibitor | 7 | 1.26 | 0 | 0 | 7 | 0 | 1 | 7 | 0 | 0 | 1 | |
| Bisphosphonate | 6 | 1.08 | 0 | 0 | 6 | 0 | 1 | 6 | 0 | 0 | 1 | |

| | | | | | | | | | | | |
|---|---|---|---|---|---|---|---|---|---|---|---|
| Voltage-gated $K^+$ channel blocker | 6 | 1.08 | 0 | 0 | 6 | 0 | 1 | 5 | 0 | 0 | 1 |
| GABAergic modulator | 6 | 1.08 | 0 | 0 | 6 | 0 | 1 | 5 | 0 | 0 | 1 |
| Microtubule modulator | 6 | 1.08 | 0 | 0 | 6 | 0 | 1 | 6 | 0 | 0 | 1 |
| Leukotriene / arachidonate pathway | 6 | 1.08 | 0 | 0 | 6 | 0 | 1 | 4 | 0 | 0 | 1 |
| NMDA receptor antagonist | 6 | 1.08 | 0 | 0 | 6 | 0 | 1 | 4 | 0 | 0 | 1 |
| DNA gyrase / topoisomerase IV inhibitor | 6 | 1.08 | 0 | 0 | 6 | 0 | 1 | 6 | 0 | 0 | 1 |
| Antiparasitic | 5 | 0.9 | 0 | 0 | 5 | 0 | 1 | 5 | 0 | 0 | 1 |
| Tyrosine kinase inhibitor | 5 | 0.9 | 0 | 0 | 5 | 0 | 1 | 5 | 0 | 0 | 1 |
| Other / Unclassified | 5 | 0.9 | 0 | 0 | 5 | 0 | 1 | 5 | 0 | 0 | 1 |
| Opioid receptor modulator | 5 | 0.9 | 0 | 0 | 5 | 0 | 1 | 5 | 0 | 0 | 1 |
| Cholinergic antagonist | 5 | 0.9 | 0 | 0 | 5 | 0 | 1 | 5 | 0 | 0 | 1 |
| Cholinergic agonist | 5 | 0.9 | 0 | 0 | 5 | 0 | 1 | 5 | 0 | 0 | 1 |
| Antiplatelet / Anticoagulant | 5 | 0.9 | 0 | 0 | 5 | 0 | 1 | 5 | 0 | 0 | 1 |
| Prostaglandin receptor agonist | 5 | 0.9 | 0 | 0 | 5 | 0 | 1 | 5 | 0 | 0 | 1 |
| Bacterial cell membrane disruptor | 4 | 0.72 | 0 | 0 | 4 | 0 | 1 | 4 | 0 | 0 | 1 |
| PPAR agonist | 4 | 0.72 | 0 | 0 | 4 | 0 | 1 | 4 | 0 | 0 | 1 |
| Retinoid receptor agonist | 4 | 0.72 | 0 | 0 | 4 | 0 | 1 | 4 | 0 | 0 | 1 |
| HMG-CoA reductase inhibitor (statin) | 4 | 0.72 | 0 | 0 | 4 | 0 | 1 | 4 | 0 | 0 | 1 |
| Other antimicrobial | 4 | 0.72 | 0 | 0 | 4 | 0 | 1 | 4 | 0 | 0 | 1 |
| Muscarinic receptor agonist | 4 | 0.72 | 0 | 0 | 4 | 0 | 1 | 4 | 0 | 0 | 1 |
| Aromatase inhibitor | 3 | 0.54 | 0 | 0 | 3 | 0 | 1 | 3 | 0 | 0 | 1 |

| | | | | | | | | | | | |
|---|---|---|---|---|---|---|---|---|---|---|---|
| DNA crosslinker | 3 | 0.54 | 0 | 0 | 3 | 0 | 1 | 3 | 0 | 0 | 1 |
| $H^{+}/K^{+}$-ATPase inhibitor (PPI) | 3 | 0.54 | 0 | 0 | 3 | 0 | 1 | 3 | 0 | 0 | 1 |
| Monoamine reuptake inhibitor (NE/DA) | 3 | 0.54 | 0 | 0 | 3 | 0 | 1 | 2 | 0 | 0 | 1 |
| Innate immune modulator | 3 | 0.54 | 0 | 0 | 3 | 0 | 1 | 3 | 0 | 0 | 1 |
| Iron chelator | 3 | 0.54 | 0 | 0 | 3 | 0 | 1 | 3 | 0 | 0 | 1 |
| IMPDH inhibitor | 3 | 0.54 | 0 | 0 | 3 | 0 | 1 | 3 | 0 | 0 | 1 |
| H2 receptor antagonist | 3 | 0.54 | 0 | 0 | 3 | 0 | 1 | 3 | 0 | 0 | 1 |
| HIV protease inhibitor | 3 | 0.54 | 0 | 0 | 3 | 0 | 1 | 3 | 0 | 0 | 1 |
| Other antifungal | 3 | 0.54 | 0 | 0 | 3 | 0 | 1 | 3 | 0 | 0 | 1 |
| Thioredoxin reductase inhibitor | 3 | 0.54 | 0 | 0 | 3 | 0 | 1 | 3 | 0 | 0 | 1 |

**Supplementary Data - Enrichment of neurotransmitter systems among screen hits.** Compounds were grouped by the neurotransmitter system their molecular target belongs to, irrespective of mechanism-of-action class or clinical indication, and each system was tested against the remainder of the library by two-sided Fisher's exact test. Compounds acting on more than one system are counted in each, so totals exceed 556. The number of compounds in the library and among the 23 hits, the hit rate, the odds ratio and the nominal p-value are shown for each system, together with the identity of the hit compounds.

| Neurotransmitter system | N (library) | % of library | N (hits) | % of hits | N (non-hits) | Hit rate (%) | OR | p-value | Hit compounds |
|---|---|---|---|---|---|---|---|---|---|
| Cholinergic | 40 | 7.19 | 5 | 21.7 | 35 | 12.5 | 3.95 | 0.01916 | Dicyclomine·HCl; Disopyramide; Glycopyrrolate Iodide; Scopolamine·HBr; Tiotropium Bromide |
| Dopaminergic | 29 | 5.22 | 3 | 13.0 | 26 | 10.3 | 2.92 | 0.11171 | Methyldopa Sesquihydrate; S(-)-Carbidopa monohydrate; Tolcapone |
| Adrenergic | 46 | 8.27 | 2 | 8.7 | 44 | 4.3 | 1.06 | 1.00000 | Methyldopa Sesquihydrate; Metoprolol Tartrate |
| Purinergic | 7 | 1.26 | 1 | 4.3 | 6 | 14.3 | 3.99 | 0.25723 | Pentoxifylline |
| Histaminergic | 34 | 6.12 | 1 | 4.3 | 33 | 2.9 | 0.69 | 1.00000 | Fexofenadine·HCl |
| Serotonergic | 39 | 7.01 | 1 | 4.3 | 38 | 2.6 | 0.59 | 1.00000 | Sumatriptan Succinate |
| Glutamatergic | 6 | 1.08 | 0 | 0.0 | 6 | 0.0 | 0.00 | 1.00000 | — |
| GABAergic | 5 | 0.90 | 0 | 0.0 | 5 | 0.0 | 0.00 | 1.00000 | — |
| Opioidergic | 5 | 0.90 | 0 | 0.0 | 5 | 0.0 | 0.00 | 1.00000 | — |
